# An Alzheimer’s disease risk variant in *TTC3* modifies the actin cytoskeleton organization and the PI3K-Akt signaling pathway in iPSC-derived forebrain neurons

**DOI:** 10.1101/2023.05.25.542316

**Authors:** Holly N. Cukier, Carolina L. Duarte, Mayra J. Laverde-Paz, Shaina A. Simon, Derek J. Van Booven, Amanda T. Miyares, Patrice L. Whitehead, Kara L. Hamilton-Nelson, Larry D. Adams, Regina M. Carney, Michael L. Cuccaro, Jeffery M. Vance, Margaret A. Pericak-Vance, Anthony J. Griswold, Derek M. Dykxhoorn

**Affiliations:** John P. Hussman Institute for Human Genomics, 1501 NW 10th Avenue, University of Miami Miller School of Medicine, Miami, FL, USA 33136; Department of Neurology, University of Miami Miller School of Medicine, 1120 NW 14th Street, Miami, FL, USA 33136; John T. Macdonald Foundation Department of Human Genetics, 1501 NW 10^th^ Avenue, University of Miami Miller School of Medicine, Miami, FL, USA 33136; JJ Vance Memorial Summer Internship in Biological and Computational Sciences, 1501 NW 10^th^ Avenue, University of Miami Miller School of Medicine, Miami, FL, USA 33136; Mental Health & Behavioral Science Service, Bruce W. Carter VA Medical Center, Miami, FL, USA, 1201 Northwest 16th Street, Miami, FL 33125

**Keywords:** Alzheimer’s disease (AD), induced pluripotent stem cells (iPSCs), *tetratricopeptide repeat domain 3* (TTC3) gene

## Abstract

A missense variant in the *tetratricopeptide repeat domain 3* (*TTC3*) gene (rs377155188, p.S1038C, NM_003316.4:c.3113C>G) was found to segregate with disease in a multigenerational family with late onset Alzheimer’s disease. This variant was introduced into induced pluripotent stem cells (iPSCs) derived from a cognitively intact individual using CRISPR genome editing and the resulting isogenic pair of iPSC lines were differentiated into cortical neurons. Transcriptome analysis showed an enrichment for genes involved in axon guidance, regulation of actin cytoskeleton, and GABAergic synapse. Functional analysis showed that the TTC3 p.S1038C iPSC-derived neuronal progenitor cells had altered 3D morphology and increased migration, while the corresponding neurons had longer neurites, increased branch points, and altered expression levels of synaptic proteins. Pharmacological treatment with small molecules that target the actin cytoskeleton could revert many of these cellular phenotypes, suggesting a central role for actin in mediating the cellular phenotypes associated with the TTC3 p.S1038C variant.

**Highlights:**

- The AD risk variant TTC3 p.S1038C reduces the expression levels of *TTC3*
- The variant modifies the expression of AD specific genes *BACE1*, *INPP5F*, and *UNC5C*
- Neurons with the variant are enriched for genes in the PI3K-Akt pathway
- iPSC-derived neurons with the alteration have increased neurite length and branching
- The variant interferes with actin cytoskeleton and is ameliorated by Cytochalasin D

## 1. Introduction

Alzheimer’s disease (AD) is the leading form of dementia in older adults (Qiu et al., 2009). The underlying genetics contributing to AD are complex, with dozens of genes and loci being associated with disease through both family and population based studies (Lambert et al, 2013; Kunkle et al, 2019; Sims et al, 2020; Bellenguez et al, 2022). We previously identified the *tetratricopeptide repeat domain 3* (*TTC3*) gene as a candidate contributing to AD risk (Kohli et al, 2016; Beecham, et al, 2018). A rare, nonsynonymous *TTC3* variant, rs377155188, was the only alteration that segregated in all 11 AD individuals in a non-Hispanic white late onset AD (LOAD) family (mean age at onset = 75.7 years, Kohli et al, 2016). This alteration results in a missense change, p.S1038C, that is predicted to be deleterious and extremely rare in the gnomAD database (allele frequency=3.231×10^-5^). Thus, we hypothesized that this variant was contributing to the AD genetic burden in this LOAD family, and that *TTC3* potentially plays a wider role in LOAD etiology. In addition to being involved in AD, it is possible that *TTC3* plays a role in other dementias as well. Indeed, a patient with frontotemporal dementia was reported who carried a TTC3 p.V1893M alteration, along with an *APOE* ε4 allele and *ADAM10* variant (Cochran, et al, 2019). The *TTC3* gene is located on chromosome 21q22.2 in the Down syndrome critical region (DCR, Ohira et al, 1996; Tsukahara et al, 1996). It encodes a 2025 amino acid protein that contains four tetratricopeptide repeat (TPR) domains in the amino terminus, followed by a potential coiled-coil domain, Citron binding region, and a carboxy terminal E3 ubiquitin ligase (E3) ring finger domain (Tsukahara et al, 1996).

Additional evidence supports the role of *TTC3* in AD pathogenesis. While *TTC3* is ubiquitously expressed, it has been shown to have elevated levels in the brain (Rachidi, et al, 2000; Fagerberg, et al, 2014). Studies reported that cortical *TTC3* expression is reduced in LOAD patients and negatively correlated with AD neuropathology (Webster et al, 2009; Zhang et al, 2013). TTC3 interacts with and mediates the ubiquitination and subsequent degradation of multiple target proteins, including heat shock proteins, polymerase gamma (POLG), and AKT. Phosphorylated AKT (AKT1, AKT2 and AKT3) is a primary target of TTC3 (Suizu et al, 2009; Kim et al, 2019). The E3 ubiquitin ligase activity of TTC3 is dependent on its phosphorylation by AKT, thus creating a negative feedback loop (Suizu et al, 2009). The serine-threonine kinase AKT plays a crucial role in the phosphatidylinositol 3-kinase (PI3K)-AKT pathway, which is activated by a wide variety of cellular signals to regulate multiple fundamental cellular processes, including cellular proliferation, survival, cell cycle arrest, cytoskeletal organization, vesicle trafficking, migration, and glucose transport, among others (Manning and Toker, 2017). Disruption of the PI3-K/Akt/mTOR pathway has been implicated in AD (O’ Neill, 2013). Additionally, *TTC3* was shown to be upregulated in neurons from mice in which the ribosome quality control pathway gene, *listerin E3 ubiquitin protein ligase 1* (*Ltn1*), had been deleted (Endo et al, 2023). These *Ltn1* knockout mice showed cognitive deficits that could be restored by the silencing of *TTC3*.

To understand the breadth of pathways impacted by a rare AD associated variant in *TTC3* (p.S1038C), isogenic iPSC lines that either carry or lack the TTC3 p.S1038C variant were derived and differentiated into forebrain neurons (Laverde-Paz et al, 2021). Transcriptome analysis (RNA-seq) of day 70 iPSC-derived neurons showed an alteration in the expression of established AD risk genes (*BACE1*, *A2M*, *INPP5F*, and *UNC5C*) as well as genes located in AD GWAS loci (*ADAMTS1*, *MAF*, and *NCK2*). The differentially expressed genes (DEGs) between the TTC3 p.S1038C variant bearing and the isogenic parental iPSC-derived neurons showed an enrichment for KEGG pathways associated with focal adhesion, PI3K-Akt signalling, axon guidance, regulation of actin cytoskeleton, and GABAergic synapse. Previous studies had shown that the silencing of *TTC3* expression led to increased neurite growth, while overexpression of *TTC3* inhibited neurite extension and interfered with the compactness of Golgi (Berto et al, 2007; Berto et al, 2014). The TTC3 p.S1038C early neurons (D30) showed increased neurite length and branch points, similar to the phenotype that was previously reported upon *TTC3* silencing. Treatment of the neurons with the selective actin polymerization inhibitor Cytochalasin D led to the reduction of neurite length and neurite branch point number in the variant-bearing neurons to levels similar to those seen in the control neurons. TTC3 p.S1038C bearing neuronal progenitor cells (NPCs) exhibited increased migration that was also ameliorated by Cytochalasin D treatment, suggesting that aberrations in the actin cytoskeleton may be driving TTC3 p.S1038C mediated neuronal irregularities.

## 2. Materials and Methods

### 2.1 Derivation of iPSCs into Forebrain Neurons

The isogenic set of iPSCs with the homozygous p.S1038C change, TTC3 clone 50 (UMi028-A-2), and the unedited parental line were both differentiated into forebrain neurons (Laverde-Paz et al, 2021). To begin, the iPSC lines were grown in mTeSR1 medium (STEMCELL Technologies, #85850) on plates coated in Matrigel hESC-Qualified Matrix (Corning, #354277) and passaged with Gentle Cell Dissociation Reagent (STEMCELL Technologies, #100-0485). To initiate differentiation and the formation of embryoid bodies on day 0, the iPSCs were washed with PBS and dissociated into single cells with Accutase (STEMCELL Technologies, #07920), filtered with a 40 μm cell strainer to remove any clumps, and counted. The iPSCs were resuspended in STEMdiff Neural Induction Medium with the SMADi Neural Induction Supplement (STEMCELL Technologies, #08581), and 10 μm Y-27623 (Biogems, #1293823) at a concentration of 3×10^6^ cells/mL. iPSCs were plated into AggreWell800 Microwell Plates (STEMCELL Technologies, #34811) with approximately 3×10^6^ cells/well or 10,000 cells/microwell, spun down at 100 x *g* for 3 minutes, and incubated at 37°C. The media in the AggreWells was partially changed (3/4 replaced) for days 1-4. On day 5, the embryoid bodies were dislodged from each AggreWell using a wide-bore P1000 tip with DMEM/F12 (Gibco/ThermoFisher Scientific, #11320033) filtered with a 40 μm cell strainer to remove single cells, replated onto 1 well of a 6 well plate in STEMdiff Neural Induction Medium with the SMADi

Neural Induction Supplement and 10 μm Y-27623, and incubated at 37°C. For days 6-11, full media changes were performed. On day 12, the cells were washed with DMEM/F12 and then incubated for ∼1.5 hours with 1 mL of STEMdiff Neural Rosette Selection Reagent (STEMCELL Technologies, #05832) in each well. Neural rosette clusters were then dislodged, resuspended in STEMdiff Neural Induction Medium with the SMADi Neural Induction Supplement and 10 μm Y-27623, and replated onto Matrigel. Cells were maintained in STEMdiff Neural Induction Medium with the SMADi Neural Induction Supplement for days 12-19 and passaged as required, at about 90% confluency. On days 20-29, the cells were grown using the daily medium changes of STEMdiff Forebrain Neuron Differentiation kit (STEMCELL Technologies, #08600) and then transitioned to the STEMdiff Forebrain Neuron Maturation kit (STEMCELL Technologies, #08605) for days 30-70 with full medium changes every 2-3 days.

For cytoskeleton rearrangement assays, the culture media was supplemented with 1 µM Cytochalasin D (Cayman Chemical, #11330), 10 nM Jasplakinolide (Cayman Chemical, # 11705) or 20 µM Y27632 (Biogems, # 1293823) and cells were cultured for 24 hrs with the supplemented medium. All were reconstituted in DMSO and the final concentration of DMSO in the culture media was between 0.2% to 0.05%.

### 2.2 Immunocytochemistry (ICC)

Cells were grown in coated 8-well chamber slides or black wall optical bottom 96-well plates and cultured in modified media as described above. Cell monolayers were washed with PBS, fixed and permeabilized using the BD Cytofix/Cytoperm™ Fixation/Permeabilization Solution Kit (BD Bioscience, #BD554715) following the manufacturer’s recommendations and incubated in blocking buffer for 1hr. Following blocking, the primary antibodies were added [Recombinant Anti-PAX6 antibody [EPR15858] (Abcam, # ab195045), Human/Mouse/Rat SOX1 Antibody (R&D Systems, # AF3369), Purified anti-Tubulin β-3 (TUBB3) Antibody 1:1000 (Biolegend, #PRB-435P), Anti-MAP2 antibody 1:1000 (Abcam, # ab5392), Monoclonal Anti-Synaptophysin 1:200 (Sigma Aldrich, # S5768), Recombinant Anti-Synapsin I antibody [EPR23531-50] 1:200 (Abcam, # ab254349), TTC3 Polyclonal Antibody 1:200 (Abcam, ab80061)] and samples incubated overnight at 4°C. The cells were washed with PBS and incubated with the appropriate secondary antibody in blocking buffer for 1 hour (donkey anti-rabbit IgG AF594 1:500, goat anti-chicken IgY AF647 1:500, goat anti-mouse IgG AF488 1:500). Phalloidin FITC Conjugate 1:1000 (Abcam, #ab235137) was added along with the secondary antibodies and incubated for 1 hour for the appropriate wells. Finally, the cells were washed with PBS and incubated with 4′,6-diamidino-2-phenylindole (DAPI) for 10 minutes and, either mounted or stored in PBS prior to imaging. The stained cells were visualized using the Zeiss Axio Observer Z.1 Confocal Microscope with Spinning Disk and images analysed with Zen 2 Blue Edition (Carl Zeiss) or the EVOS™ FL Auto 2 Imaging System (Invitrogen) and images analysed using Celleste Image Analysis Software (Thermo Fisher Scientific). For the cytoskeleton rearrangement assays, the cells were imaged in the ArrayScan XTI (Thermo Scientific) and analysed using the Cellomics Morphology Explorer V4 in HCS Studio (Thermo Scientific).

### 2.3 Quantitative Polymerase Chain Reaction (qPCR)

Cells were collected at day 0, day 30, and day 70 and stored in RNAprotect (Qiagen, #76106) at −20°C. To extract the RNA, the samples were thawed and processed using the Qiagen RNeasy Mini Kit (Qiagen, Cat #74104) according to the manufacturer’s protocol. Cell lysates were homogenized using QIAshredder spin columns (Qiagen, Cat #79656) and genomic DNA was removed by an on-column DNase treatment (Qiagen, Cat #79254). 500 ng of total RNA was used for each sample for reverse transcription (RT) into cDNA using the iScript Reverse Transcription Supermix for RT-qPCR (Bio-Rad, #1708840). RNA expression levels for *TTC3* (Hs01598175_m1) and *GAPDH* (Hs99999905_m1) were measured in triplicate using TaqMan Gene Expression assays (Thermo Fisher Scientific) on the QuantStudio 12 k Flex Real-Time PCR system machine.

### 2.4 Western blot analysis

Cells were lysed at day 0, day 30, and day 70 in RIPA Lysis and Extraction Buffer (Thermo Fisher Scientific, Cat # 89901) supplemented with Halt™ Protease and Phosphatase Inhibitor Cocktail, EDTA-Free (100X, Thermo Scientific™, Cat# 78441). The lysates were denatured, separated in 4–15% Mini-PROTEAN® TGX™ Precast Protein Gels (Bio-Rad, # 4561084DC) and transferred to a membrane in a Trans-Blot® Turbo™ Mini PVDF Transfer Pack (Bio-Rad, Cat# 1704156EDU). The membrane was blocked 1 hr with 5% Blotting Grade Blocker Non-Fat Dry Milk (Bio-Rad, Cat# 1706404XTU) and incubated overnight with the primary antibodies [TTC3 Polyclonal Antibody (Abcam, # AB80061), Recombinant Anti-AKT1 + AKT2 + AKT3 antibody [EPR16798] (Abcam, # AB179463), Recombinant Anti-AKT1 + AKT2 + AKT3 (phospho S472 + S473 + S474, Purified anti-Tubulin β-3 (TUBB3) Antibody 1:1000 (Biolegend, # PRB-435P), anti-GAPDH antibody 1:1000 (Sigma Aldrich, #SAB1405848), Monoclonal Anti-Synaptophysin 1:200 (Sigma Aldrich, # S5768), Recombinant Anti-Synapsin I antibody [EPR23531-50] 1:200 (Abcam, # ab254349),) antibody [EPR18853], (Abcam, # AB192623)] in blocking buffer. The cells were incubated with HRP-linked secondary antibodies, and the protein bands were detected using the chemiluminescent Clarity Max™ Western ECL Substrate (Bio-Rad, Cat# 1705062S). The bands were visualized in the ChemiDoc Imaging System (Bio-Rad) and quantified in ImageLab 6.0.1 (Bio-Rad).

### 2.5 Neurite Tracking/Neurogenesis

Cells were grown in PLO/laminin coated 96-well plates, cultured in modified media as described above, and imaged live using the IncuCyte® Zoom system for automated live cell analysis (Sartorius). Time-lapse images were acquired every six hours. The growth rate of neurites in each well was obtained by measuring the length of neurites that extend from the cell bodies expressed as neurite length/surface area (mm/mm^2^), and the intersection of two masked neurites in an image expressed as branch points/cell body cluster count (1/cell body cluster).

### 2.6 Cell Migration

A monolayer scratch assay was performed on a 96-well Essen ImageLock plate (Sartorius) using the IncuCyte® WoundMaker Tool (Sartorius) and cells were imaged live every 6 hours using the IncuCyte® ZOO system for automated live cell analysis (Sartorius). Wound closure was analysed by measuring the spatial cell density in the wound area relative to the spatial cell density outside of the wound expressed as the Relative Wound Density (%) and the confluence of cells within the wound region expressed as Wound Confluence (%).

### 2.7 Chemotaxis Assay

Chemotaxis was assessed using a PLO/laminin coated µ-Slide Chemotaxis (iBidi, #80236) following the manufacturer’s recommended protocol and using 500µM Dibutyryl-cAMP (Stemcell Technologies, #73882) as the chemotactic agent. The EVOS™ FL Auto 2 Imaging System (Invitrogen) was used for time lapse image acquisition and cell tracking was performed using the FastTrack AI image analysis software (MetaVi Labs).

### 2.8 RNA sequencing (RNA-Seq) and analysis

The RNA samples used for quantitative RT-PCR were the same used for the transcriptomic analysis. The concentration and quality of RNA samples from day 70 was assessed using a Bioanalyzer RNA kit (Agilent Technologies). Total RNA was prepped with the NuGEN Universal with AnyDeplete Human Ribo kit from ∼100 ng of total RNA input. Libraries were sequenced on the Illumina NovaSeq 6000, yielding a minimum of 35 million reads/sample. Raw FASTQs were processed by a bioinformatics pipeline including adapter trimming by TrimGalore (v0.6.1) (https://github.com/FelixKrueger/TrimGalore), alignment with the STAR alignerv2.5.0a to the GRCh38 human reference, and gene counts quanitified against the GENCODEv35 gene annotation release using the GeneCounts module implemented in STAR.

Differential expression analysis was conducted using edgeR with a false discovery rate (FDR) cut-off of 0.05 to correct for the multiple comparison testing and fold change (FC) of +/- 1.25 (Robinson et al, 2010). Pathway enrichment with differentially expressed protein coding genes was performed using the clusterProfiler package (v4.4.4) which supports the latest online version of KEGG data from the KEGG website (Yu et al, 2012; Chen et al, 2013).

### 2.9 Statistical Analysis

Two means were compared using the Student’s t-test (unpaired, two tailed). Three or more means were analysed by one-way or two-way analysis of variance, followed by Dunnett’s test for pairwise comparisons. Time lapse assays were analysed using repeated measures analysis of variance followed by Dunnett’s test for pairwise comparisons or by Mann-Whitney test for frequencies and ratios. A p-value ofL0.05 was considered significant. All statistical analyses were performed in GraphPad Prism version 9 (GraphPad Software).

## 3. Results and Discussion

### 3.1 iPSC-derived neurons bearing the TTC3 variant showed differences in known and novel AD genes compared to the isogenic unedited neurons

Global transcriptional analysis of day 70 neurons demonstrated a wide range of differentially expressed genes, including those previously implicated in AD, as well as novel genes. 978 protein coding genes were differentially expressed (FDR<0.05, FC +/-1.25) between the isogenic cells with and without the *TTC3* alteration (Table 1, Supplemental Table 1). The volcano plot show the 381 upregulated and 597 down regulated genes and the heat map also demonstrate that there is a distinct transcriptional profile between the edited and unedited neurons (Fig. 1A). Hierarchal clustering demonstrated that the neurons derived from the edited and unedited cells are distinct from each other (Fig. 1B). In addition, the TTC3 p.S1038C variant-bearing neurons showed a −1.31-fold change (FDR=8.70×10^-9^) in *TTC3* expression compared to the unedited cell line. This reduction in *TTC3* expression was confirmed by qRT-PCR analysis on day 70, as well as iPSC and day 30 neuronal progenitor cells (NPCs) (Supplemental Fig. 1A). Immunoblot analysis confirmed this decrease at the protein level (Supplementary Fig. 1B).

**Figure 1.**
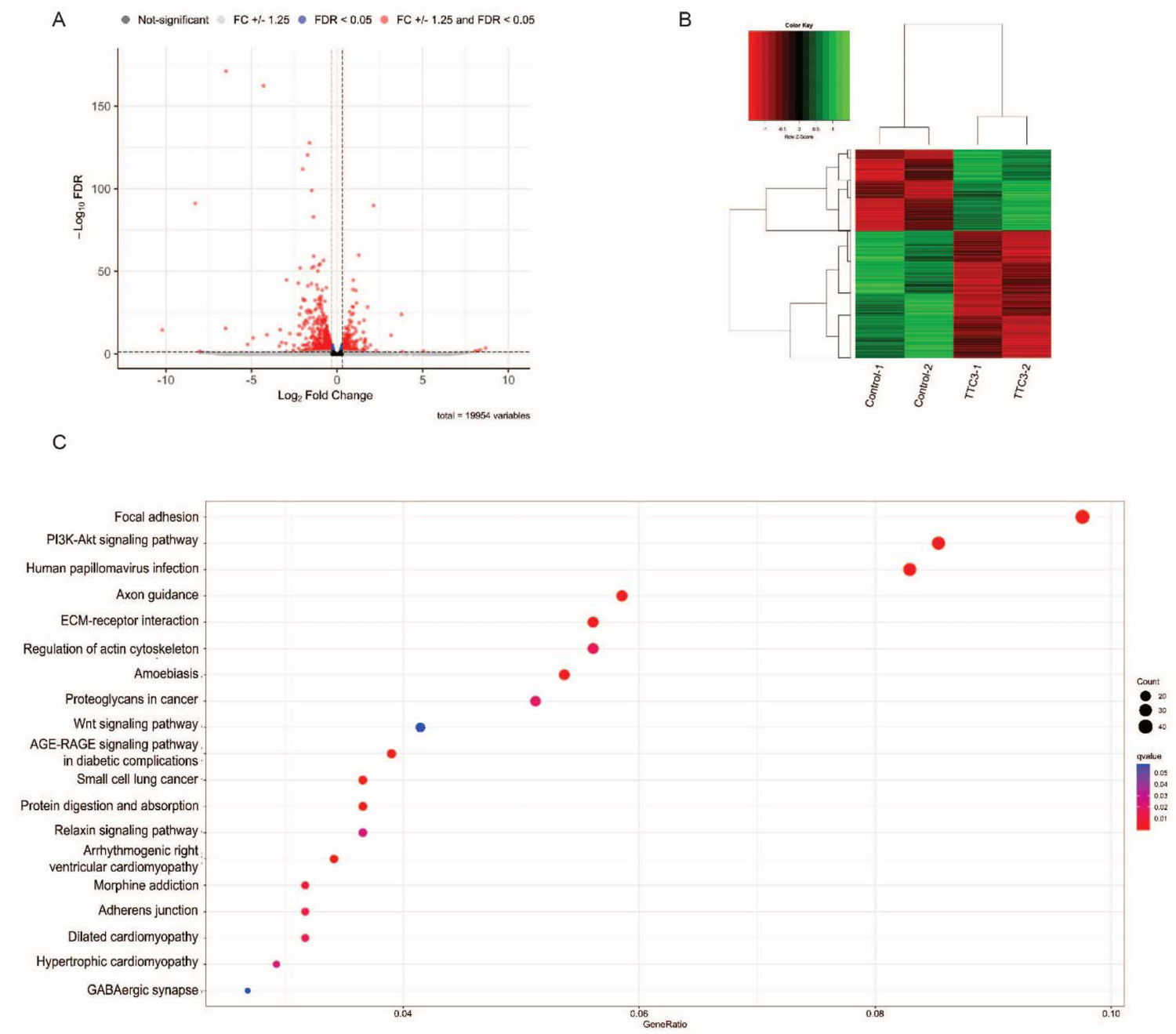
Transcriptional analysis of day 70 neurons. **A**. Volcano plot of the 381 upregulated and 597 down regulated genes that were significant between the neurons with and without the *TTC3* alteration. Genes marked in red have a FDR < 0.05 and FC +/-1.25. **B.** Heat map of the 978 significantly differentially expressed genes with a false discovery rate (FDR) < 0.05 and fold change (FC) +/-1.25 **C.** KEGG pathway analysis showing cellular functions overrepresented in the DEGs from the parental compared to TTC3 p.S1038C derived neurons.

**Table 1.**
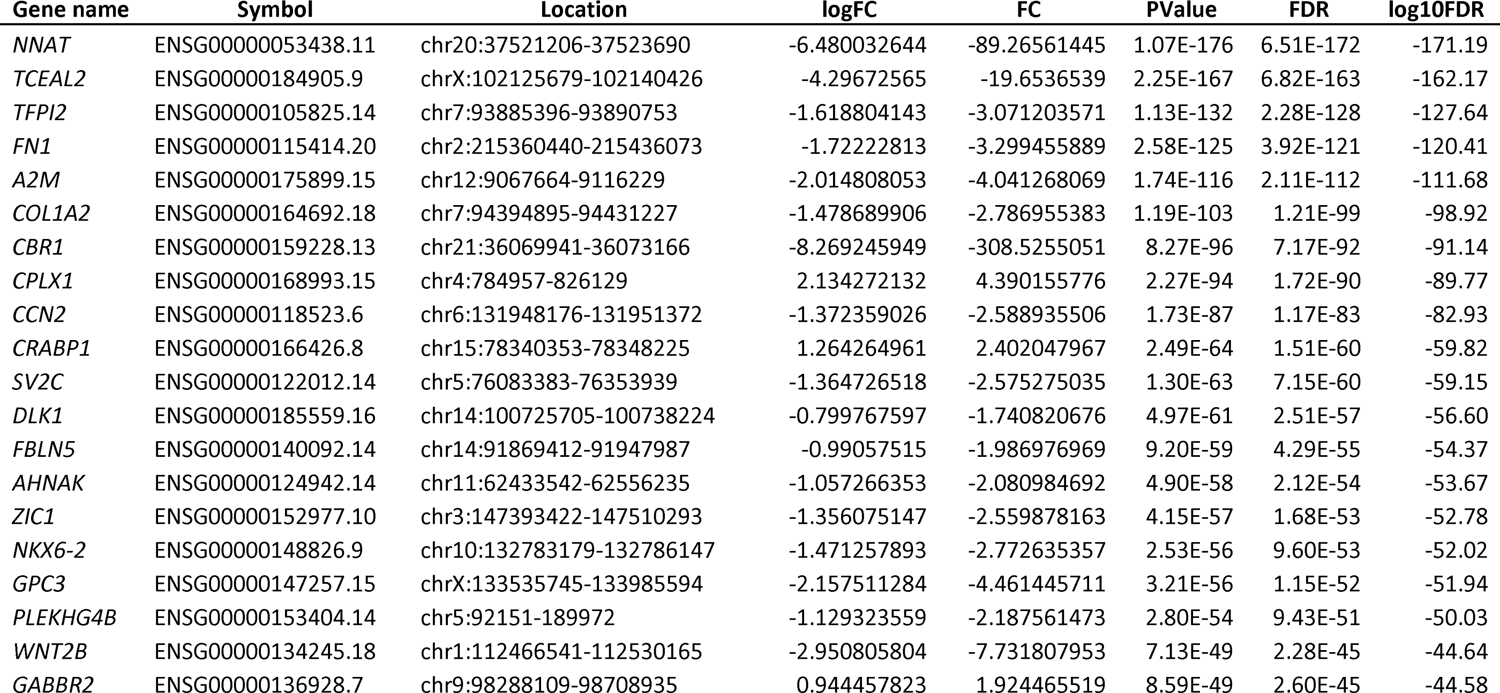
Top 20 significantly differentially expressed genes

Significant differentially expressed genes were also overlaid with known AD genes and loci (Lambert, et al, 2013, Kunkle, et al, 2019, Bellenguez, et al, 2022). Although no overall enrichment was seen for AD GWAS genes, four of the significantly differentially expressed genes identified in this study have been previously implicated in AD: *beta-secretase 1* (*BACE1*), *alpha-2-macroglobulin* (*A2M*), *unc-5 netrin receptor C* (*UNC5C*) and *inositol polyphosphate-5-phosphatase F* (*INPP5F*, Wetzel-Smith, et al, 2014, Jiao, et al, 2014, Korvatska, et al, 2015, Xue, et al, 2022). Differential expression was identified for *BACE1* with an increase of 1.28-fold change in the *TTC3* variant-bearing neurons compared to the control neurons (FDR=2.39×10^-4^, Supplemental Table 1). *BACE1* is one of the first genes found to cause AD (Vassar, et al, 1999). *BACE1* encodes for a secretase that cleaves the amyloid precursor protein (APP), which is the product from which beta amyloid fragments are generated. *A2M* encodes a pan-protease inhibitor and chaperone that is involved in innate immunity which can clear and degrade amyloid beta and has been suggested to be a preclinical marker of AD (Blacker, et al, 1998, Varma, et al, 2017). More recently, studies identified numerous rare *UNC5C* alterations that segregate with disease in LOAD families, as well as a broader association with four large case-control datasets (Wetzel-Smith, et al, 2014, Jiao, et al, 2014, Korvatska, et al, 2015). In addition, imaging studies have demonstrated that polymorphisms in *UNC5C* can influence cognition over time, as well as the volume and atrophy rate of key brain regions, including the hippocampus, middle temporal, and precuneus areas (Sun, et al, 2015, Yang, et al, 2019). *INPP5F* was also associated with AD, in addition to being connected to Parkinson’s Disease (Nalls, et al, 2014, Blauwendraat, et al, 2019, Xue, et al, 2022). Moreover, *INPP5F* is in the same gene family as *INPP5D*, a significant AD GWAS locus (Kunkle, et al, 2019, Bellenguez, et al, 2022).

In addition, three genes located within established genome wide association study (GWAS) regions were found to be differentially regulated: *ADAM metallopeptidase with thrombospondin type 1 motif 1* (*ADAMTS1*), *MAF bZIP transcription factor* (*MAF*), and *NCK adaptor protein 2* (*NCK2*, Supplemental Table 1, Kunkle, et al, 2019, Schwartzentruber, et al, 2021, Bellenguez, et al, 2022). *ADAMTS1* has been shown to be upregulated in the brains of AD patients and has been specifically connected to both brain amyloidosis and neurodegeneration (Ferrando Miguel, et al, 2005, Medoro, et al, 2019, Tan, et al, 2021). Variants in *MAF* have also been associated with limbic-predominant age-related TDP-43 encephalopathy neuropathological changes (LATE-NC), a type of dementia similar to AD (Nelson, et al, 2019, Dugan, et al, 2022). Intriguingly, *NCK2* has been shown to be involved in actin cytoskeleton dynamics as well as vascular remodeling (Rohatgi, et al, 2001, Alfaidi, et al, 2021) Additionally, some of the most significantly differentially expressed genes have already been shown to be critical for neuronal function. *Neuronatin* (*NNAT*) was the most significant differentially expressed gene (FDR=6.51×10^-176^) and was strongly downregulated in the edited neurons (FC=-89.3). *NNAT* is an imprinted gene that encodes a proteolipid critical during brain development to regulate ion channels and can act in an anti-inflammatory manner (Joseph, et al, 1995, Oyang, et al, 2011, Ka, et al, 2017). *Complexin 1* (*CPLX1*, FDR=1.72×10^-90^, FC=4.4), the highest gene that was upregulated in the edited neurons, encodes a protein involved in synaptic vesicle exocytosis and was previously shown to influence cognitive decline in humans, as well it was shown to be differentially expressed in a mouse AD model (Chang, et al, 2015, Ramons-Miguel, et al, 2018, Jesko, et al, 2021). *Synaptic Vesicle Glycoprotein 2C* (*SV2C*, FDR=7.15×10^-60^, FC=-2.6) encodes a synaptic vesicle protein involved in dopamine regulation and is altered in post-mortem tissue from patients with Parkinson’s disease (Dunn, et al, 2017, Stout, et al, 2019). *SV2C* has a relatively narrow expression in dopaminergic neurons, and some GABAergic and cholinergic neurons (Dardou, et al, 2011). Intriguingly, the closely related paralogs *SV2A* and *SV2B* both have altered expression in other AD models, with *SV2B* specifically being upregulated in the presence of Aβ42, reviewed by Bartholome, et al, 2017 (Heese, et al, 2001, Gomez Ravetti, et al, 2010, Stockburger, et al, 2016, Chen, et al, 2018).

### 3.2 Pathway analysis demonstrated alterations in multiple pathways

KEGG pathway analysis of the 979 significantly differentially regulated genes identified 23 pathways with an adjusted P-value < 0.05 (Fig. 1C, Table 2). This includes the PI3K-Akt signalling pathway, in which *TTC3* has been previously implicated (adjusted p=7.99×10-5), as well as the axon guidance pathway, the regulation of actin cytoskeleton pathway, the GABAergic synapse pathway, and the Wnt signalling pathway (Fig. 1C). 39 genes in the PI3K-Akt signalling pathway, or 11% of the 354 genes recognized by KEGG, were significantly differentially expressed. Moreover, the most differentially expressed gene, *C-X-C Motif Chemokine Ligand 11* (*CXCL11*, FDR=4.08×10^-15^, FC=-1187), encodes a pro-inflammatory chemokine that depends on the AKT pathway (Callahan, et al, 2021). AKT signalling has been associated with a wide variety of cellular phenotypes from proliferation and cell cycle progression to cell survival. metabolism, and actin reorganization (Franke 2008). In neurons, AKT signalling has been shown to be important for protection against trophic factor deprivation, ischemic injury, and oxidative stress (Dudek et al, 1997; Salinas et al, 2001; Noshita et al, 2002). Dysregulation of AKT signalling has also been observed in Alzheimer disease with multiple possible means of action being suggested, including alteration to GSK3β signalling and Tau phosphorylation, to the response to insulin and induction of neuroinflammation (Rickle et al, 2004; Ryder et al, 2004; Wang et al, 2018; Hoyer 2002; van der Heide et al, 2005; Chiu et al, 2008; Capiralla et al, 2012). In fact, many of the pathways identified in our transcriptome analysis can be related back to AKT signalling.

**Table 2.** Significant KEGG pathways and genes

| Term | Overlap | P-value | Adjusted P-value | Odds Ratio |
| --- | --- | --- | --- | --- |
| Focal adhesion | 45/201 | 4.62E-18 | 1.32E-15 | 5.832921098 |
| ECM-receptor interaction | 27/88 | 5.53E-15 | 7.91E-13 | 8.824998707 |
| Amoebiasis | 24/102 | 9.43E-11 | 8.99E-09 | 6.109982261 |
| Proteoglycans in cancer | 28/205 | 8.68E-07 | 5.53E-05 | 3.138031519 |
| Human papillomavirus infection | 38/331 | 9.67E-07 | 5.53E-05 | 2.584060707 |
| AGE-RAGE signaling pathway in diabetic complications | 18/100 | 1.50E-06 | 7.17E-05 | 4.330792683 |
| PI3K-Akt signaling pathway | 39/354 | 1.96E-06 | 7.99E-05 | 2.466565242 |
| Axon guidance | 25/182 | 2.98E-06 | 1.06E-04 | 3.152131051 |
| Small cell lung cancer | 16/92 | 9.01E-06 | 2.82E-04 | 4.146186673 |
| Protein digestion and absorption | 17/103 | 9.86E-06 | 2.82E-04 | 3.895070542 |
| Arrhythmogenic right ventricular cardiomyopathy | 14/77 | 1.92E-05 | 4.99E-04 | 4.370447211 |
| Morphine addiction | 15/91 | 3.30E-05 | 7.86E-04 | 3.883013609 |
| Regulation of actin cytoskeleton | 24/218 | 1.79E-04 | 0.003772994 | 2.441548337 |
| Relaxin signaling pathway | 17/129 | 1.85E-04 | 0.003772994 | 2.986751152 |
| GABAergic synapse | 13/89 | 3.76E-04 | 0.007173163 | 3.358303791 |
| Dilated cardiomyopathy | 13/96 | 7.89E-04 | 0.014097705 | 3.0739372 |
| Hippo signaling pathway | 18/163 | 0.001059654 | 0.017827118 | 2.440991379 |
| Wnt signaling pathway | 18/166 | 0.001307032 | 0.020767286 | 2.391131757 |
| Hypertrophic cardiomyopathy | 12/90 | 0.001420274 | 0.02137886 | 3.01704093 |
| Tight junction | 18/169 | 0.001601787 | 0.022905551 | 2.343253311 |
| TGF-beta signaling pathway | 12/94 | 0.002072614 | 0.028227031 | 2.869262233 |
| Adherens junction | 10/71 | 0.002291585 | 0.029790608 | 3.21111638 |
| Complement and coagulation cascades | 11/85 | 0.002809557 | 0.034936235 | 2.912714162 |

### 3.3 Migration and chemotaxis are altered in TTC3 p.S1038C variant bearing neuronal progenitor cells compared to control NPCs

TTC3 activates the RhoA-PIIA actin polymerization and stabilization pathway that can alter NPC migration (Berto et al, 2007, Berto, et al, 2014). Therefore, actin cytoskeleton irregularities can be expected in cases of *TTC3* mutation. To test cell migration, we performed wound healing assays on neuronal progenitor cells (NPCs) from the TTC3 p.S1038C and isogenic control lines. In day 30 NPCs, cell cultures with the *TTC3* variant were able to recover more quickly from a scratch wound than unedited NPCs (Fig. 2 A-C). This was shown from the higher relative wound density (i.e. faster closing of the wound) as measured by time lapse imaging on the Incucyte® Zoom (Sartorius, Fig. 2B). In addition, the relative wound confluence (a measure of cell confluence in the wound area) was found to be modestly, but statistically significantly, higher in the TTC3 S.1038C variant bearing compared to the control NPCs (Fig. 2C). Migration of the NPCs towards a chemoattractant (Dibutyryl cyclic AMP (DBcAMP)) was assessed using a µ-Slide Chemotaxis assay (iBiDi). Consistent with the wound healing assay, NPCs bearing the p.S1038C variant had faster speed (as measured by the total distance travelled over time) and velocity (as measured by the vectorial cell displacement over time) compared to the isogenic control NPCs (Fig. 2F-H). There appeared to be no difference in the directionality of the migration since both the p.S1038C variant bearing and control NPCs migrated towards the chemoattractant; however, stronger chemotaxis was observed in the p.S1038C variant bearing NPCs as measured by the forward migration index values. The forward migration index on the X-axis represents the lateral displacement of cell movement that is not directed to the region of higher chemoattractant concentration. Therefore, the lower forward migration index on the X-axis seen on the p.S1038C variant bearing NPCs represents a stronger response to the chemoattractant and less lateral displacement during movement as evidenced in the endpoint tracking diagram (Fig. 2D,E). The morphometric analysis of phallodin-stained day 30 NPCs bearing the *TTC3* p.S1038C variant showed an increase in the 3D structure of the cells as measured by the oblate and prolate volumes (Fig. 2G,H) compared to the unedited control NPCs while no differences in object entropy intensity or fiber length or width was observed (Fig. 2G,H).

**Figure 2.**
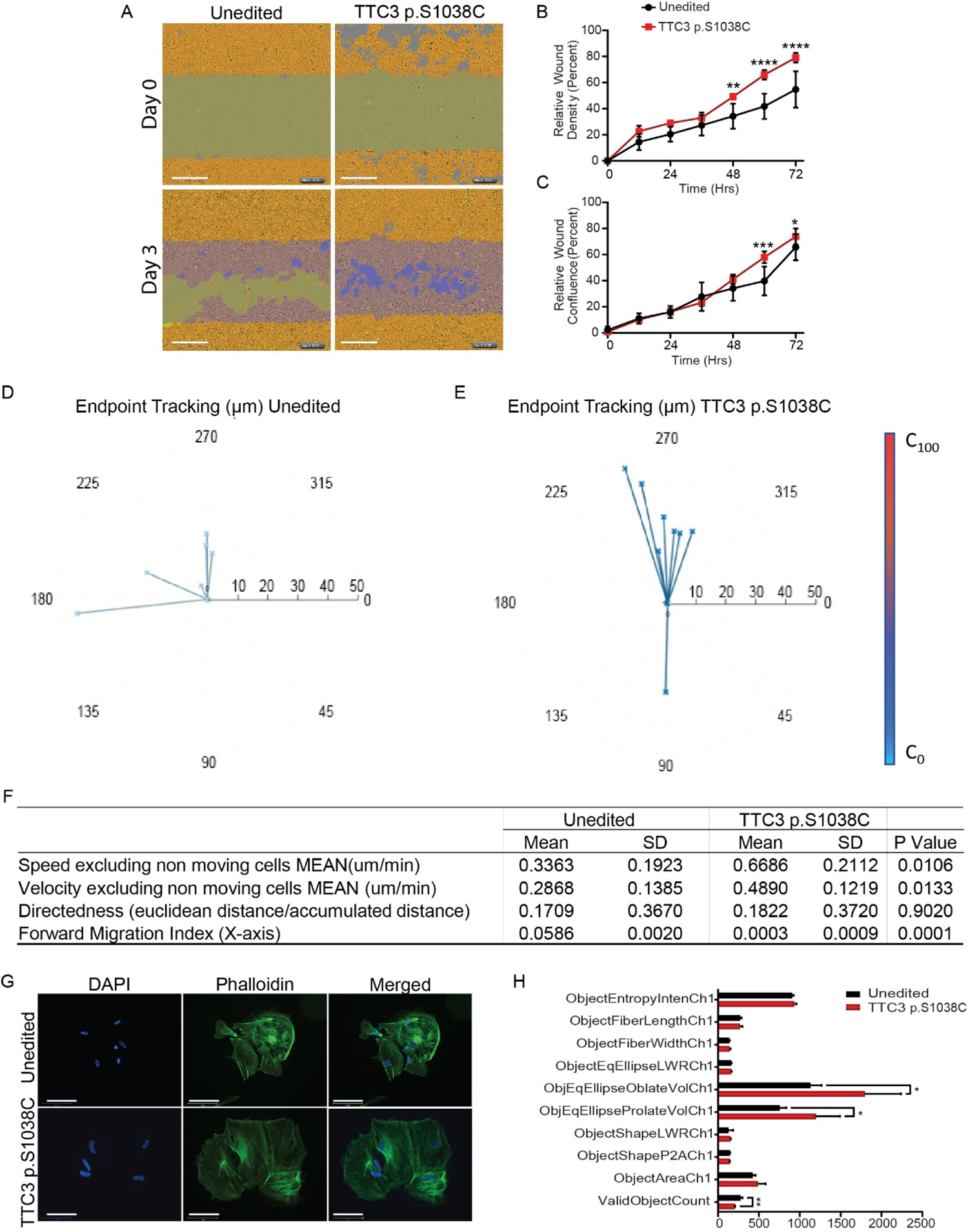
Neuronal Precursor Cell Migration and Cytoskeleton Assessment. **A-C.** Scratch wound assay on D30 neuron precursor cells recording cell migration for three days. Migration rate was measured as the wound area relative to the spatial cell density outside of the wound (Wound Density) and the confluence of cells within the wound region (Wound Confluence). Scale Bars: 300 µm. **D-E.** Endpoint tracking of migrating cells using the µ-Slide Chemotaxis assay. The start point for each tracked cell is located in the center of the diagram and the vectorial displacement is delineated in the direction of movement. The Y-axis represents movement along the chemoattractant gradient, which is higher towards the top of the diagram, and the X-axis represents movement perpendicular to the chemoattractant gradient. **F.** Chemotaxis analysis. **G.** Representative images of D30 neurons. Blue: Nuclear counterstain DAPI, Green: Actin filament stain Phalloidin, Scale Bars: 125 µm. **H.** Immunofluorescent assessment of actin filament organization and cytoskeleton pattern analysis of D30 neurons. Ch1: Phalloidin. *: p-value<0.05, **: p-value<0.01, ***: p-value<0.001, ****: p-value<0.0001

### 3.4 Neurite length and number of branch points were elevated in the TTC3 p.S1038C variant bearing neurons compared to the control neurons

Similar morphometric analysis was performed on the day 70 neurons. Unlike the NPCs, the *TTC3* p.S1038C variant bearing neurons showed no statistically significant differences in the 3D structure of the cell bodies (i,.e. no difference in the oblate or prolate volume). However, the TTC3 p.S1038C variant bearing neurons had higher entropy of the actin microfilaments (as measured by Phalloidin staining, Fig. 3A,B, Ch1) and microtubules (as measured by TUBB3 staining, Fig. 3A,B, Ch3) compared to the control unedited neurons. These results suggest that the ultrastructural organization of the neuronal cytoskeleton is affected by the *TTC3* mutation and aligns with previous reports of *TTC3* silencing (Berto et al, 2014).

**Figure 3.**
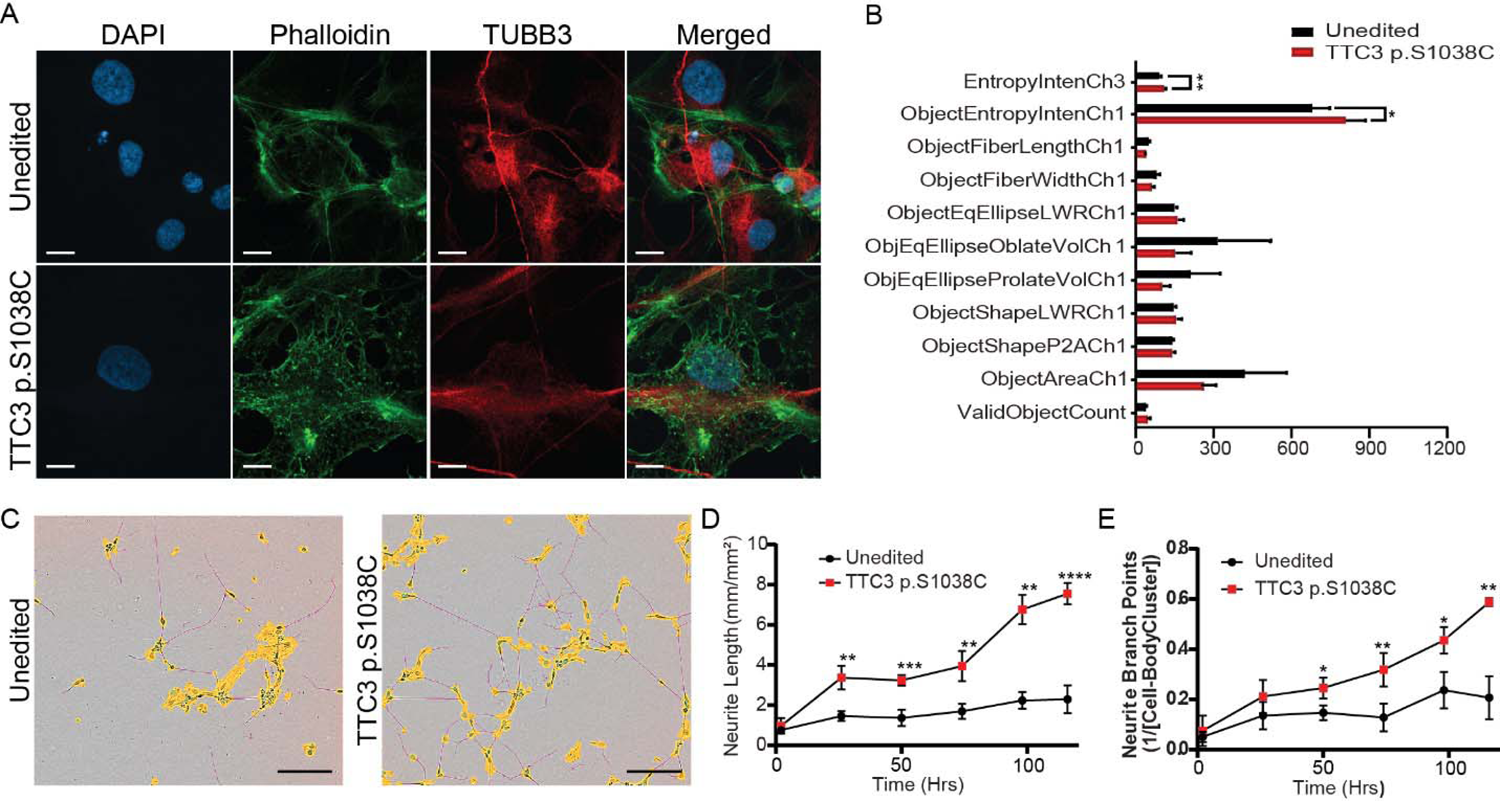
iPSC-Derived Neuron Cytoskeleton Assessment and Neurite Formation. **A.** Representative images of D70 neurons. Blue: Nuclear counterstain DAPI, Green: Actin filament stain Phalloidin, Red: β-tubulin 3 (TUBB3). Scale Bars: 125 µm **C-F.** Immunofluorescent assessment of actin filament organization and cytoskeleton pattern analysis of D70 neurons Ch1: Phalloidin, Ch3: TUBB3. **C-E.** Representative images and quantification of neurite formation recorded from D30 to D35 and presented as neurite length per surface area. The intersection of two masked neurites in an image is presented as branch points/cell body cluster count. Scale Bars: 150 µm. *: p-value<0.05, **: p-value<0.01, ***: p-value<0.001, ****: p-value<0.0001

The transcriptome analysis highlighted differences in axon guidance. In addition, there is evidence that shows that modulation of *TTC3* affects neurite growth (Berto et al, 2007; Berto et al, 2014). Therefore, morphological measures of neurite growth and branching were assessed in differentiating neuronal cultures using live cell imaging on the Incucyte® Zoom (Sartorius). The TTC3 p.S1038C variant bearing neurons had increased neurite length (Fig. 3D), which phenocopies previous studies in which *TTC3* expression was silenced in rat hippocampal neurons using small interfering RNAs (siRNAs, Berto et al, 2007, Berto, et al, 2014). In addition, the edited neurons had significantly more branch points/cell body than the isogenic control neurons (Fig. 3E). Interestingly, the cell body clusters formed in the edited cells were less voluminous than those observed in control cells, with more single cells developing neurites and neuron-to-neuron connections (Fig. 3C).

Defects in neurite outgrowth and branching could affect the transport of molecules down the axon to the synapses. The transcriptome analysis also showed an enrichment for differentially expressed genes corresponding to “GABAergic synapse”. Therefore, we examined the levels of expression of Synapsin 1 (SYN1) and Synaptophysin (SYP). Interestingly, the TTC3 p.S1038C variant bearing neurons had increased expression of SYN1, while the level of SYP was decreased in these same cells compared to the isogenic control neurons (Fig. 4A-G). This dysregulation of synaptic protein expression may affect vesicle transport, fusion, and recycling in TTC3 p.S1038C neurons. SYN1 is a synaptic vesicle marker that can directly bind to PIIA (Witke et al, 1998), and both proteins have been associated with synaptic vesicle traffic and endocytic assembly (Witke et al, 1998; Shah and Rossie, 2018). Similarly, SYP, an integral membrane protein found in pre-synaptic vesicles, is associated with presynaptic vesicle trafficking, and is believed to play a role in pore complex formation during vesicle fusion (Thiel, 1993; Rao et al, 2017). Furthermore, SYP concentration in neurons is reduced during aging and, more remarkably, in AD (Hamos et al, 1989; Masliah and Terry, 1993).

**Figure 4.**
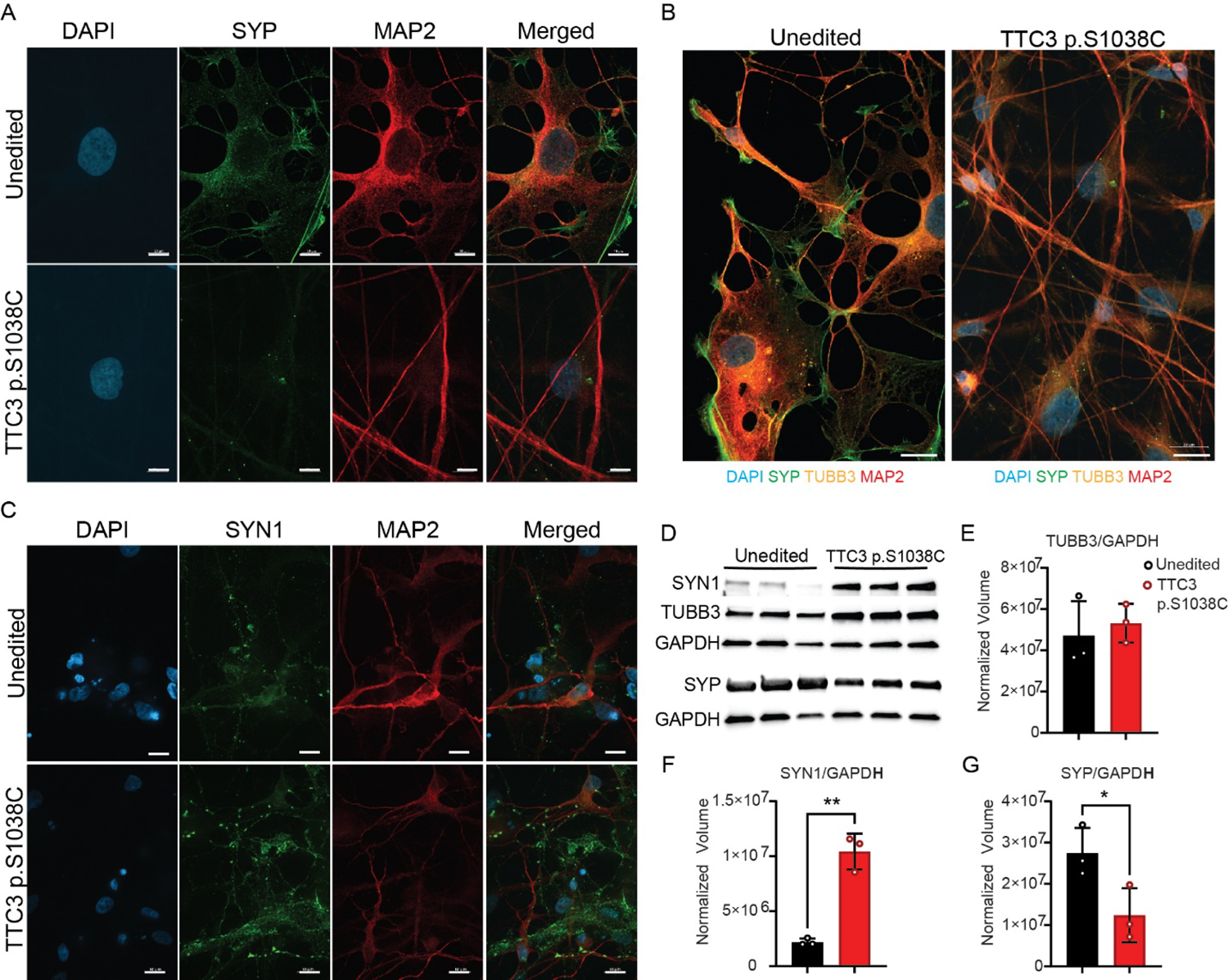
Markers of Neuronal Differentiation and Synaptic Vesicle Formation. **A-C.** Immunofluorescent detection of markers of neuronal differentiation on D70 neurons. Scale Bars A & C: 10 µm. Scale Bars B: 20 µm. **D-G.** Western blot analysis and quantification of the expression of β-tubulin 3 and synaptic vesicle markers Synapsin 1 and Synaptophysin. β-tubulin 3 (TUBB3), Microtubule Associated Protein 2 (MAP2), Synapsin 1 (SYN1) and Synaptophysin (SYP). *: p-value<0.05, **: p-value<0.01, ***: p-value<0.001, ****: p-value<0.0001

### 3.5 Differences in NPC migration and neuronal outgrowth seen in the TTC3 p.S1038C variant bearing cells could be modulated by small molecules that alter the stability of the actin cytoskeleton

The phenotypes identified with the NPCs (morphology and migration) and the neurons (morphology, neurite outgrowth, and branch point number) from the TTC3 p.S1038C variant bearing iPSC can all be connected through the modulation of actin stability. Therefore, we tested the effect of several small molecules that are known to affect actin polymerization, either directly by interacting with actin or by targeting the upstream regulator of actin polymerization, Rho Associated Coiled-Coil Containing Protein (ROCK). Cytochalasin D (CytoD) is a potent inhibitor of actin polymerization which binds to the F-actin polymer and preventing the addition of actin monomers, as well as disrupting the microfilament network organization (Schliwa, 1982). Jasplakinolide (Jaspla) is an actin filament polymerizing and stabilizing small molecule, that can also contribute to neurite extension and microtubule plasticity in dendrites (Holzinger, 2001). Finally, Y27362 is a selective and potent inhibitor of ROCK1 and ROCK2 that contributes to actin cytoskeleton stabilization and neuritogenesis (Da Silva, et al, 2003; Shi et al, 2013).

The cytoskeletal characteristics that were previously found to be significantly different between unedited and *TTC3* NPCs and neurons (Fig. 2C,D; Fig. 3A,B) were assessed after treatment with CytoD, Jaspla or Y27632. Microscopic assessment of the cytoskeleton of edited and unedited cells after treatment with CytoD showed dramatic actin bundling and cytoskeleton collapse; thus, CytoD treatment could disrupt the three-dimensional cell volume (prolate and oblate volume) in day 30 NPCs derived from both the TTC3 p.S1038C variant bearing and isogenic control iPSCs (Fig. 5A,C,D). Microscopic assessment of the Jaspla treated *TTC3* mutants showed improved cytoskeleton organization that approximated the morphological characteristics of control cells (Fig. 5A,C,D). Although the treatment with Jaspla improved the morphology of the day 30 TTC3 variant bearing NPCs, it had no effect on the microtubule entropy (as measured by TUBB3 staining, Ch3) of the day 70 *TTC3* variant bearing neurons (Fig.5B,E,F). On the other hand, Jaspla treatment of the isogenic control D70 neurons had a statistically significant increase in microtubule entropy (Ch3) so that they resemble the *TTC3* variant bearing neurons (Fig. 5B,F). Although Y27632 treatment had no significant effect on the *TTC3* variant bearing neurons, it showed a statistically significant increase in both the actin filament entropy (phallodin staining, Ch1) and microtubule entropy (TUBB3 staining, Ch3) of the unedited neurons so that they more closely resemble the *TTC3* variant bearing neurons. No difference from the untreated NPCs were seen with the *TTC3* variant bearing and control NPCs treated with Y27632 (Fig. 5A,C,D).

**Figure 5.**
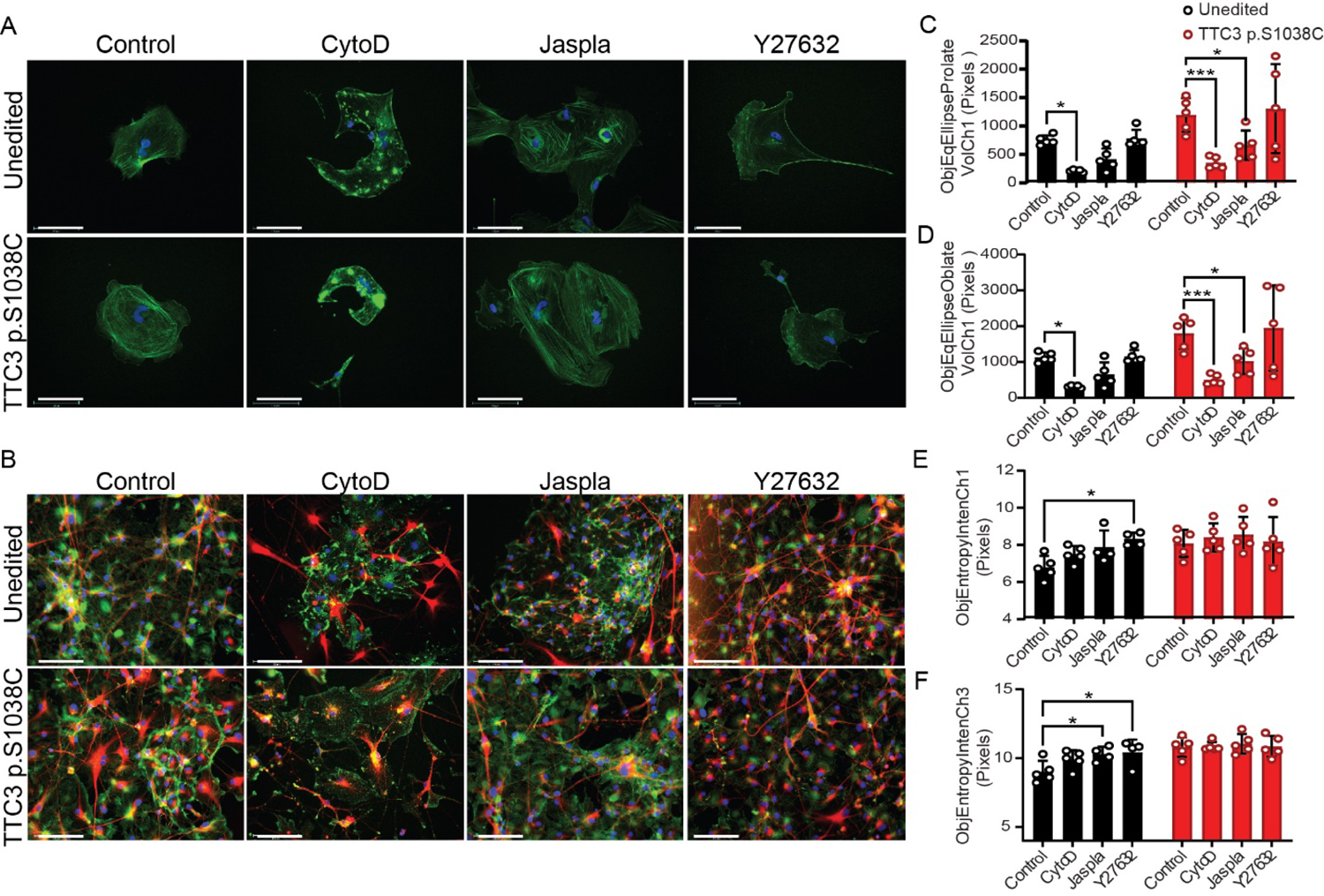
Cytoskeleton Rearrangement Assay of Neuronal Precursor Cells and iPSC-Derived Neurons. **A.** Representative images of D30 neuron progenitor cells. Blue: Nuclear counterstain DAPI, Green: Actin filament stain Phalloidin. Scale Bars: 125 µm **B.** Representative images of D70 neurons. Blue: Nuclear counterstain DAPI, Green: Actin filament stain Phalloidin, Red: β-tubulin 3. Scale Bars: 125 µm **C-D.** Comparison of cytoskeleton characteristics that were significantly different between D30 (**C-D**) and D70 (**E-F**) TTC3 pS1038C cells and unedited cells after treatment with Cytochalasin D (CytoD), Jasplakinolide (Jaspla) and Y27632. Ch1: Phalloidin, Ch3: β-tubulin 3. *: p-value<0.05, ***: p-value<0.001.

We next tested the impact of modulating the stability of the actin cytoskeleton on NPC migration (Fig. 6A-E) and neurite branching and outgrowth (Fig. 6F-H). The treatment with CytoD showed a significant decrease in the migration of both the *TTC3* variant bearing and the isogenic control NPCs. The treatment of the control NPCs with Y27632 had a significant increase in migration into the wound (Relative wound density) that resembled that of the untreated *TTC3* variant bearing NPCs (Fig. 6A-E). Jaspla had no effect on the control NPCs but showed a decrease in migration of the *TTC3* variant bearing NPCs. The treatment of *TTC3* variant bearing neurons with CytoD significantly decreased the neurite branch points/cell body and the neurite length so that they more closely resemble that of the isogenic control neurons (Fig. 6F-H). On the other hand, the treatment of the day 70 neurons (both isogenic control and the *TTC3* variant bearing) with Y27632 led to increases in neurite length. The results of these analysis show that the alteration of NPC morphology and migration and neurite length and branching in the TTC3 p.S1038C variant bearing cells is regulated, at least in part, through the modulation of actin polymerization.

**Figure 6.**
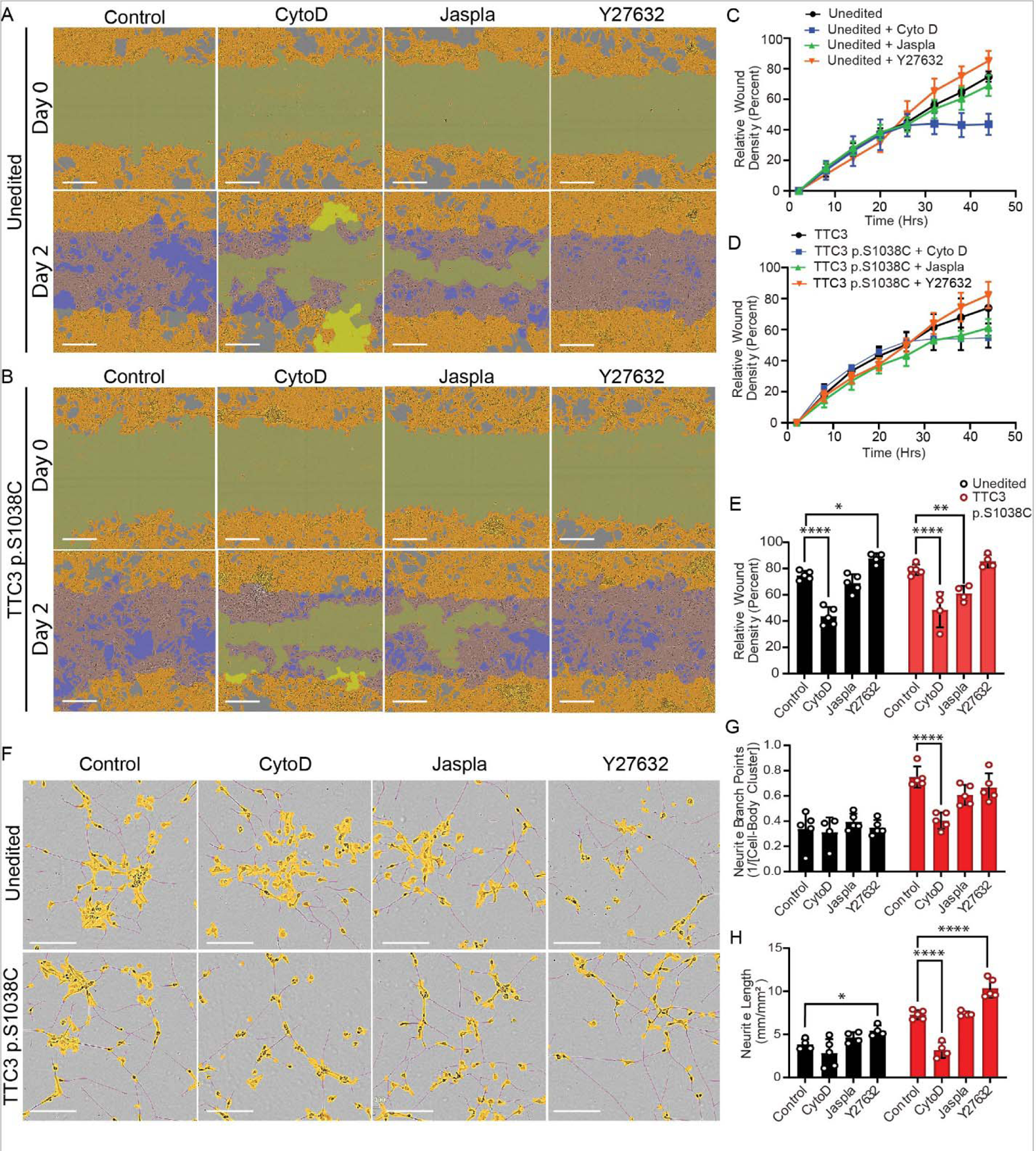
Migration and Neurite Formation of TTC3 pS1038C Bearing Neuronal Precursor Cells after Treatment with Actin Disrupting Agents. **A-E.** Scratch wound assay on D30 neuron precursor cells recording cell migration 24 hours before and 24 hours after treatment with Cytochalasin D (CytoD), Jasplakinolide (Jaspla) and Y27632. Migration rate was measured as the wound area relative to the spatial cell density outside of the wound (Wound Density) throughout the duration of the experiment (C-D) and at endpoint (E). Scale Bars: 300 µm **F-H.** Representative images and quantification of neurite formation after treatment with Cytochalasin D (CytoD), Jasplakinolide (Jaspla) and Y27632, recorded from D30 to D35 and presented as neurite length per surface area. The intersection of two masked neurites in an image presented as branch points/cell body cluster count. Scale Bars: 150 µm. *: p-value<0.05, **: p-value<0.01, ****: p-value<0.0001.

Altogether, these results demonstrate that ROCK inhibition by Y27632 induces phenotypical changes in unedited cells that resemble those observed in the TTC3 p.S1038C cells, which suggest the RhoA-PIIA pathway is, indeed, responsible for structural and physiological changes in these TTC3 mutation bearing cells. These results also suggest that corrections in ATP-Actin availability and actin microfilament stability at specific timepoints may both be necessary to rescue the TTC3 p.S1038 phenotype. Jaspla is an actin microfilament polymerization inducer that acts on thymosin-β4 to increase the availability of ATP-Actin that contributes to microfilament elongation (Bubb et al, 2000). Its effect on NPCs suggests that the availability of ATP-actin for microfilament polymerization is essential for ultrastructural integrity and mobility of NPCs, which is dysregulated by the inhibition of TTC3 activity. The actin microfilament polymerization inhibitor CytoD can bind to the ends of the actin microfilament and prevent monomer association, sever actin filaments and cause actin aggregation (Schliwa, 1982). Although CytoD reduced cell migration and neuritogenesis due to its drastic effects on the actin microfilaments, it did not improve the cytoskeletal patterns of the TTC3 pS1038C neurons. In addition, the pathway analysis also correlated directly with some of the cellular phenotypes that we characterised. Treatment of the neurons with the actin polymerization inhibitor Cytochalasin D reversed the enhanced neurite length and neurite branch points in the edited cells. Moreover, the actin cytoskeleton pathway was also dysregulated in the pathway analysis (adjusted p value = 3.8×10^-3^). This finding adds to the work of previous studies demonstrating a connection of *TTC3* to actin organization (Berto, et al, 2014).

The TTC3 E3 ubiquitin ligase function maybe also be contributing to AD pathogenesis through additional mechanisms. In addition to AKT, both POLG and SMAD ubiquitination regulatory factor 2 (SMURF2) have been identified as targets and others may yet be discovered (Gong et al, 2017; Kim, et al, 2019). POLG is a critical replication and repair enzyme in the mitochondria (Gong et al, 2017). In addition, through the degradation of SMURF2, TTC3 influences TGF-β signalling (Kim, et al, 2019). While just one of hundreds of E3 ligases, TTC3 is an intriguing potential therapeutic target for AD by small molecules (Morreale and Walden, 2016; Gong, et al 2016). Moreover, disruption of the Ubiquitin Proteasome System (UPS) has been suggested as a contributing mechanism of neurodegeneration (Upadhyay et al, 2017). *TTC3* is just one of numerous ubiquitin signalling genes connected to neurodegenerative diseases (Schmidt et al, 2021). Additional ubiquitin ligases implicated in AD include *CHIP*, *HRD1*, *Nedd4*, and *RNF182* (Sahara et al, 2005; Kaneko et al, 2010; Rodrigues et al, 2016; Liu et al, 2008).

Consequences from modulation of *TTC3* may be complex, for in addition to having numerous isoforms expressed, the gene has also been shown to generate circular and long noncoding RNAs (Cai et al, 2019, Zhang et al, 2021). These RNAs have demonstrated a myriad of functions, including regulating apoptosis in retinal ganglion cells and sequestering microRNAs to modulate cardiac function and protect against inflammation, oxidative stress, and hypoxic injury (Cai et al, 2019, Yu et al, 2020, Ma et al, 2021, Zhang, et al, 2021). In addition, *TTC3-AS1* has been shown to be differentially expressed in Parkinson’s disease (Huang et al, 2022). Thus, the involvement of *TTC3* with neurodegeneration may be both nuanced and multifaceted, requiring additional examination.

## 4. Conclusions

The results from this study suggest that the TTC3 p.S1038C variant causes a partial loss of function. This, along with the lower cortical *TTC3* expression in LOAD patients and an inverse relationship between *TTC3* levels and AD neuropathology, may indicate that reduced levels may contribute to AD pathology, and not merely be a consequence of the disease (Webster et al, 2009; Zhang et al, 2013).

## Acknowledgements

Funding: This work was supported by the Florida Department of Health Ed and Ethel Moore Alzheimer’s Disease Research Program [grant number 7AZ20 to HNC], and the National Institutes of Health [grant number R25NS090624 to JMV].

---

Supplemental Figure 1. T*T*C3 Expression **A.** *TTC3* mRNA expression assessed by qPCR. **B.** Western blot analysis and quantification of TTC3 protein expression. **C.** Immunofluorescent detection of TTC3 in cell monolayers. Scale bars: 10 µm. **D.** Western Blot membrane images. Black arrow heads on C-E point to the TTC3 expected molecular weight of ∼229 kDa. Red arrow on C points to a low molecular weight band of ∼125 kDA observed in iPSCs. *: p-value<0.05.

## Antibody List and Dilutions

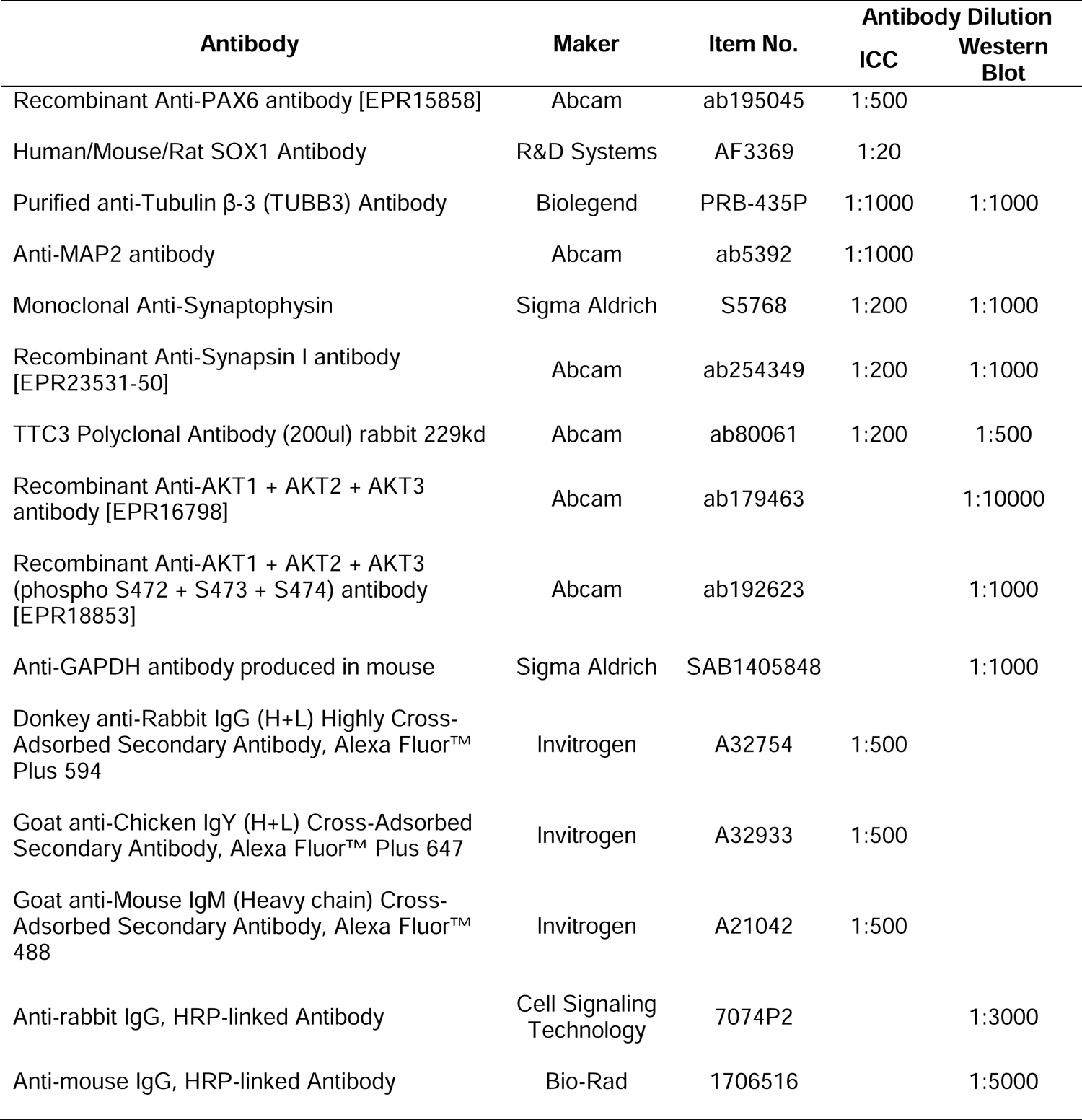

