## Supplementary figures and images for "An Alzheimer’s disease risk variant in *TTC3* modifies the actin cytoskeleton organization and the PI3K-Akt signaling pathway in iPSC-derived forebrain neurons"

### Supplemental Figure 1

Supplemental Figure 1

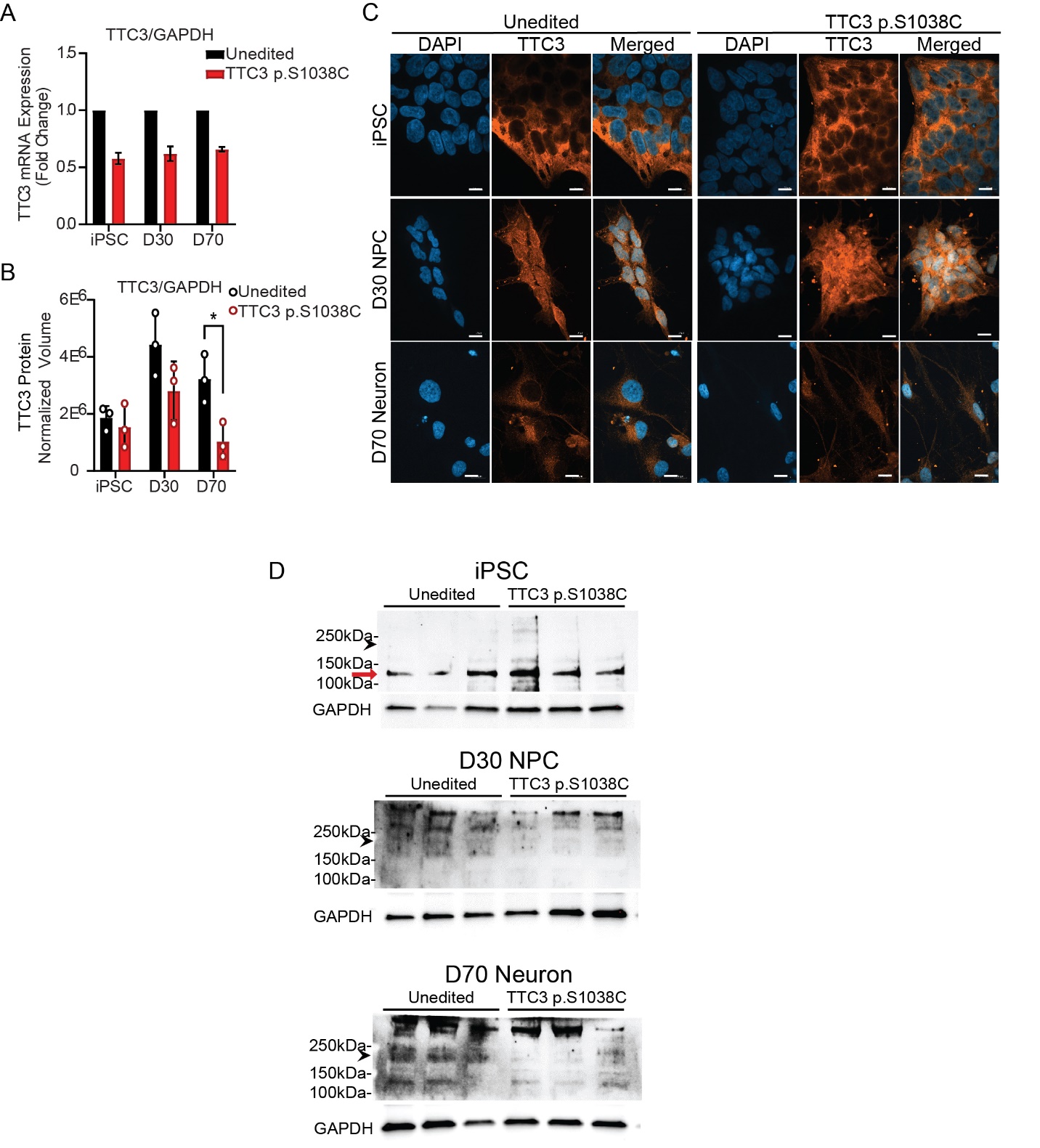
