## Supplemental Table 1 for "An Alzheimer’s disease risk variant in *TTC3* modifies the actin cytoskeleton organization and the PI3K-Akt signaling pathway in iPSC-derived forebrain neurons"

| **Supplemental Table 1. Significant differentially expressed genes** | | | | | | | | | |
| --- | --- | --- | --- | --- | --- | --- | --- | --- | --- |
| **Gene name** | **Symbol** | **Location** | **logFC** | **FC** | **logCPM** | **F** | **PValue** | **FDR** | **log10FDR** |
| *NNAT* | ENSG00000053438.11 | chr20:37521206-37523690 | -6.48 | -89.27 | 4.47 | 805.89 | 1.07E-176 | 6.51E-172 | -171.19 |
| *TCEAL2* | ENSG00000184905.9 | chrX:102125679-102140426 | -4.30 | -19.65 | 5.06 | 762.75 | 2.25E-167 | 6.82E-163 | -162.17 |
| *TFPI2* | ENSG00000105825.14 | chr7:93885396-93890753 | -1.62 | -3.07 | 8.83 | 602.28 | 1.13E-132 | 2.28E-128 | -127.64 |
| *FN1* | ENSG00000115414.20 | chr2:215360440-215436073 | -1.72 | -3.30 | 7.99 | 568.28 | 2.58E-125 | 3.92E-121 | -120.41 |
| *A2M* | ENSG00000175899.15 | chr12:9067664-9116229 | -2.01 | -4.04 | 6.68 | 527.51 | 1.74E-116 | 2.11E-112 | -111.68 |
| *COL1A2* | ENSG00000164692.18 | chr7:94394895-94431227 | -1.48 | -2.79 | 8.45 | 468.28 | 1.19E-103 | 1.21E-99 | -98.92 |
| *CBR1* | ENSG00000159228.13 | chr21:36069941-36073166 | -8.27 | -308.53 | 3.35 | 432.12 | 8.27E-96 | 7.17E-92 | -91.14 |
| *CPLX1* | ENSG00000168993.15 | chr4:784957-826129 | 2.13 | 4.39 | 5.96 | 425.49 | 2.27E-94 | 1.72E-90 | -89.77 |
| *CCN2* | ENSG00000118523.6 | chr6:131948176-131951372 | -1.37 | -2.59 | 8.31 | 393.76 | 1.73E-87 | 1.17E-83 | -82.93 |
| *CRABP1* | ENSG00000166426.8 | chr15:78340353-78348225 | 1.26 | 2.40 | 7.61 | 287.13 | 2.49E-64 | 1.51E-60 | -59.82 |
| *SV2C* | ENSG00000122012.14 | chr5:76083383-76353939 | -1.36 | -2.58 | 7.07 | 283.83 | 1.30E-63 | 7.15E-60 | -59.15 |
| *DLK1* | ENSG00000185559.16 | chr14:100725705-100738224 | -0.80 | -1.74 | 11.67 | 271.95 | 4.97E-61 | 2.51E-57 | -56.60 |
| *FBLN5* | ENSG00000140092.14 | chr14:91869412-91947987 | -0.99 | -1.99 | 9.38 | 261.53 | 9.20E-59 | 4.29E-55 | -54.37 |
| *AHNAK* | ENSG00000124942.14 | chr11:62433542-62556235 | -1.06 | -2.08 | 8.71 | 258.18 | 4.90E-58 | 2.12E-54 | -53.67 |
| *ZIC1* | ENSG00000152977.10 | chr3:147393422-147510293 | -1.36 | -2.56 | 6.71 | 253.92 | 4.15E-57 | 1.68E-53 | -52.78 |
| *NKX6-2* | ENSG00000148826.9 | chr10:132783179-132786147 | -1.47 | -2.77 | 6.34 | 250.31 | 2.53E-56 | 9.60E-53 | -52.02 |
| *GPC3* | ENSG00000147257.15 | chrX:133535745-133985594 | -2.16 | -4.46 | 5.01 | 249.84 | 3.21E-56 | 1.15E-52 | -51.94 |
| *PLEKHG4B* | ENSG00000153404.14 | chr5:92151-189972 | -1.13 | -2.19 | 7.77 | 240.92 | 2.80E-54 | 9.43E-51 | -50.03 |
| *WNT2B* | ENSG00000134245.18 | chr1:112466541-112530165 | -2.95 | -7.73 | 3.96 | 216.08 | 7.13E-49 | 2.28E-45 | -44.64 |
| *GABBR2* | ENSG00000136928.7 | chr9:98288109-98708935 | 0.94 | 1.92 | 8.90 | 215.71 | 8.59E-49 | 2.60E-45 | -44.58 |
| *PDLIM1* | ENSG00000107438.9 | chr10:95237572-95291012 | -2.27 | -4.81 | 4.56 | 207.24 | 6.02E-47 | 1.74E-43 | -42.76 |
| *WIF1* | ENSG00000156076.10 | chr12:65050626-65121305 | -1.49 | -2.81 | 5.86 | 205.56 | 1.39E-46 | 3.84E-43 | -42.42 |
| *FLNB* | ENSG00000136068.15 | chr3:58008400-58172251 | -1.18 | -2.27 | 6.85 | 202.89 | 5.32E-46 | 1.40E-42 | -41.85 |
| *HMCN2* | ENSG00000148357.16 | chr9:130265882-130434123 | -1.70 | -3.24 | 5.34 | 199.62 | 2.75E-45 | 6.94E-42 | -41.16 |
| *PEG10* | ENSG00000242265.5 | chr7:94656325-94669695 | -0.63 | -1.55 | 12.34 | 194.55 | 3.49E-44 | 8.46E-41 | -40.07 |
| *TNKS* | ENSG00000173273.16 | chr8:9555912-9782346 | -1.14 | -2.20 | 6.92 | 190.95 | 2.13E-43 | 4.97E-40 | -39.30 |
| *FAM171B* | ENSG00000144369.13 | chr2:186694060-186765959 | 0.93 | 1.90 | 8.46 | 189.60 | 4.19E-43 | 9.42E-40 | -39.03 |
| *RCAN1* | ENSG00000159200.18 | chr21:34513142-34615113 | -0.87 | -1.82 | 9.05 | 187.38 | 1.27E-42 | 2.76E-39 | -38.56 |
| *PMP2* | ENSG00000147588.7 | chr8:81440326-81447439 | 1.09 | 2.13 | 7.09 | 186.24 | 2.26E-42 | 4.72E-39 | -38.33 |
| *RTL1* | ENSG00000254656.2 | chr14:100879753-100903722 | -0.93 | -1.91 | 7.87 | 170.65 | 5.66E-39 | 1.14E-35 | -34.94 |
| *THBS1* | ENSG00000137801.11 | chr15:39581079-39599466 | -1.04 | -2.05 | 7.06 | 166.43 | 4.73E-38 | 9.25E-35 | -34.03 |
| *COL12A1* | ENSG00000111799.21 | chr6:75084326-75206053 | -1.97 | -3.93 | 4.58 | 163.75 | 1.81E-37 | 3.43E-34 | -33.46 |
| *GNG11* | ENSG00000127920.6 | chr7:93921735-93928610 | -0.94 | -1.92 | 7.63 | 162.90 | 2.78E-37 | 5.11E-34 | -33.29 |
| *MAMDC2* | ENSG00000165072.10 | chr9:70043848-70226972 | -1.07 | -2.09 | 6.74 | 160.60 | 8.83E-37 | 1.57E-33 | -32.80 |
| *TPBG* | ENSG00000146242.9 | chr6:82363206-82367420 | -1.86 | -3.64 | 4.71 | 158.77 | 2.21E-36 | 3.83E-33 | -32.42 |
| *PAPPA* | ENSG00000182752.10 | chr9:116153791-116402321 | -1.96 | -3.89 | 4.55 | 158.56 | 2.47E-36 | 4.15E-33 | -32.38 |
| *ZNF185* | ENSG00000147394.18 | chrX:152914442-152973480 | -1.28 | -2.43 | 5.92 | 158.35 | 2.73E-36 | 4.47E-33 | -32.35 |
| *TAGLN* | ENSG00000149591.17 | chr11:117199370-117207464 | -0.96 | -1.94 | 7.41 | 157.01 | 5.35E-36 | 8.54E-33 | -32.07 |
| *FLNA* | ENSG00000196924.18 | chrX:154348524-154374638 | -0.72 | -1.65 | 9.90 | 155.88 | 9.45E-36 | 1.47E-32 | -31.83 |
| *IRX1* | ENSG00000170549.4 | chr5:3595832-3601403 | 1.14 | 2.20 | 6.30 | 151.03 | 1.08E-34 | 1.64E-31 | -30.78 |
| *FGFBP2* | ENSG00000137441.8 | chr4:15960245-15969309 | -0.97 | -1.96 | 7.09 | 148.35 | 4.16E-34 | 6.15E-31 | -30.21 |
| *EMP2* | ENSG00000213853.10 | chr16:10528422-10580632 | -1.13 | -2.19 | 6.29 | 147.27 | 7.18E-34 | 1.04E-30 | -29.98 |
| *PCDHGB6* | ENSG00000253305.2 | chr5:141408021-141512979 | -1.03 | -2.05 | 6.65 | 146.67 | 9.67E-34 | 1.36E-30 | -29.86 |
| *SYNPO* | ENSG00000171992.13 | chr5:150601080-150659207 | -0.93 | -1.90 | 7.32 | 144.60 | 2.74E-33 | 3.78E-30 | -29.42 |
| *BCAN* | ENSG00000132692.19 | chr1:156641390-156659532 | 0.91 | 1.88 | 7.34 | 140.70 | 1.95E-32 | 2.63E-29 | -28.58 |
| *SALL3* | ENSG00000256463.8 | chr18:78980275-79002677 | 1.78 | 3.43 | 4.62 | 139.84 | 3.01E-32 | 3.97E-29 | -28.40 |
| *PCDHGA3* | ENSG00000254245.3 | chr5:141343829-141512975 | 0.94 | 1.91 | 7.13 | 139.69 | 3.24E-32 | 4.18E-29 | -28.38 |
| *LRP2* | ENSG00000081479.15 | chr2:169127109-169362534 | -1.03 | -2.04 | 6.55 | 138.87 | 4.89E-32 | 6.18E-29 | -28.21 |
| *TCEAL5* | ENSG00000204065.3 | chrX:103273691-103276750 | -2.09 | -4.26 | 4.15 | 137.94 | 7.81E-32 | 9.67E-29 | -28.01 |
| *FBLN2* | ENSG00000163520.14 | chr3:13549125-13638422 | -1.23 | -2.34 | 5.75 | 134.95 | 3.51E-31 | 4.26E-28 | -27.37 |
| *CALD1* | ENSG00000122786.20 | chr7:134744252-134970729 | -0.91 | -1.88 | 7.19 | 134.86 | 3.68E-31 | 4.38E-28 | -27.36 |
| *ALPK2* | ENSG00000198796.7 | chr18:58481247-58629091 | -1.28 | -2.44 | 5.48 | 131.19 | 2.33E-30 | 2.72E-27 | -26.57 |
| *SLC7A2* | ENSG00000003989.18 | chr8:17497088-17570573 | -1.64 | -3.12 | 4.78 | 131.14 | 2.39E-30 | 2.73E-27 | -26.56 |
| *FNDC1* | ENSG00000164694.17 | chr6:159169400-159272108 | -1.78 | -3.44 | 4.48 | 128.31 | 9.93E-30 | 1.12E-26 | -25.95 |
| *CYP1B1* | ENSG00000138061.12 | chr2:38066973-38109902 | -1.55 | -2.93 | 4.90 | 127.35 | 1.61E-29 | 1.77E-26 | -25.75 |
| *CHD8* | ENSG00000100888.15 | chr14:21385194-21456126 | -0.88 | -1.84 | 7.22 | 126.28 | 2.75E-29 | 2.98E-26 | -25.53 |
| *NEDD4L* | ENSG00000049759.19 | chr18:58044226-58401540 | 0.74 | 1.67 | 8.65 | 126.08 | 3.06E-29 | 3.25E-26 | -25.49 |
| *SLIT2* | ENSG00000145147.20 | chr4:20251905-20620561 | -1.51 | -2.85 | 4.95 | 124.32 | 7.40E-29 | 7.74E-26 | -25.11 |
| *BNC2* | ENSG00000173068.18 | chr9:16409503-16870843 | -1.69 | -3.24 | 4.57 | 122.48 | 1.87E-28 | 1.92E-25 | -24.72 |
| *GPNMB* | ENSG00000136235.17 | chr7:23235967-23275108 | 0.94 | 1.92 | 6.63 | 121.62 | 2.88E-28 | 2.91E-25 | -24.54 |
| *HPD* | ENSG00000158104.11 | chr12:121839527-121863596 | -2.19 | -4.56 | 3.83 | 118.76 | 1.21E-27 | 1.21E-24 | -23.92 |
| *PARM1* | ENSG00000169116.11 | chr4:74933095-75050115 | -0.79 | -1.73 | 7.75 | 118.60 | 1.32E-27 | 1.29E-24 | -23.89 |
| *MYH14* | ENSG00000105357.18 | chr19:50188186-50310542 | 3.76 | 13.56 | 2.52 | 118.42 | 1.44E-27 | 1.39E-24 | -23.86 |
| *PCDH7* | ENSG00000169851.15 | chr4:30720415-31146805 | 0.85 | 1.81 | 7.17 | 117.40 | 2.41E-27 | 2.29E-24 | -23.64 |
| *CTNNA2* | ENSG00000066032.18 | chr2:79185231-80648861 | -0.83 | -1.77 | 7.36 | 116.00 | 4.88E-27 | 4.55E-24 | -23.34 |
| *CPNE4* | ENSG00000196353.11 | chr3:131533555-132285410 | -1.05 | -2.07 | 6.02 | 112.46 | 2.91E-26 | 2.67E-23 | -22.57 |
| *DST* | ENSG00000151914.20 | chr6:56457987-56954649 | -0.62 | -1.53 | 9.78 | 111.74 | 4.19E-26 | 3.79E-23 | -22.42 |
| *ARHGAP29* | ENSG00000137962.13 | chr1:94148988-94275068 | -1.38 | -2.61 | 5.02 | 110.23 | 8.97E-26 | 8.00E-23 | -22.10 |
| *ITGB3* | ENSG00000259207.9 | chr17:47253827-47313743 | -1.61 | -3.06 | 4.51 | 108.39 | 2.26E-25 | 1.99E-22 | -21.70 |
| *NKAIN3* | ENSG00000185942.12 | chr8:62248591-63014000 | -0.71 | -1.64 | 8.22 | 107.62 | 3.33E-25 | 2.89E-22 | -21.54 |
| *GATM* | ENSG00000171766.17 | chr15:45361124-45402327 | -0.85 | -1.80 | 6.83 | 106.27 | 6.60E-25 | 5.64E-22 | -21.25 |
| *ZNF469* | ENSG00000225614.4 | chr16:88382959-88440757 | -1.12 | -2.18 | 5.56 | 104.79 | 1.39E-24 | 1.17E-21 | -20.93 |
| *BCKDHB* | ENSG00000083123.15 | chr6:80106647-80346270 | -0.74 | -1.68 | 7.70 | 104.19 | 1.88E-24 | 1.56E-21 | -20.81 |
| *SERPINE1* | ENSG00000106366.9 | chr7:101127104-101139247 | -1.16 | -2.24 | 5.41 | 102.69 | 4.00E-24 | 3.28E-21 | -20.48 |
| *ECEL1* | ENSG00000171551.12 | chr2:232479827-232487834 | -0.97 | -1.96 | 6.14 | 102.64 | 4.11E-24 | 3.32E-21 | -20.48 |
| *SMOC1* | ENSG00000198732.11 | chr14:69854131-70032366 | 0.73 | 1.66 | 7.75 | 101.99 | 5.71E-24 | 4.52E-21 | -20.34 |
| *IRX2* | ENSG00000170561.13 | chr5:2745845-2751677 | 1.57 | 2.97 | 4.45 | 101.98 | 5.74E-24 | 4.52E-21 | -20.34 |
| *IRS1* | ENSG00000169047.5 | chr2:226731317-226799759 | -0.86 | -1.82 | 6.63 | 101.86 | 6.08E-24 | 4.70E-21 | -20.33 |
| *FAT3* | ENSG00000165323.15 | chr11:92352096-92896470 | -0.66 | -1.58 | 8.70 | 101.85 | 6.12E-24 | 4.70E-21 | -20.33 |
| *IFITM1* | ENSG00000185885.16 | chr11:313506-315272 | -1.81 | -3.52 | 4.09 | 101.60 | 6.94E-24 | 5.26E-21 | -20.28 |
| *RHOBTB2* | ENSG00000008853.17 | chr8:22987417-23020199 | -0.70 | -1.62 | 8.11 | 101.37 | 7.79E-24 | 5.83E-21 | -20.23 |
| *SLCO1C1* | ENSG00000139155.9 | chr12:20695332-20753386 | -0.71 | -1.63 | 7.98 | 100.42 | 1.26E-23 | 9.29E-21 | -20.03 |
| *ITGA8* | ENSG00000077943.8 | chr10:15513954-15719922 | -1.84 | -3.58 | 4.03 | 100.32 | 1.32E-23 | 9.66E-21 | -20.02 |
| *CAVIN1* | ENSG00000177469.13 | chr17:42402449-42423256 | -0.91 | -1.87 | 6.37 | 99.17 | 2.36E-23 | 1.69E-20 | -19.77 |
| *ID3* | ENSG00000117318.9 | chr1:23557926-23559501 | -1.29 | -2.45 | 5.05 | 98.80 | 2.85E-23 | 2.01E-20 | -19.70 |
| *CCN3* | ENSG00000136999.5 | chr8:119416446-119424434 | 0.84 | 1.79 | 6.64 | 98.45 | 3.40E-23 | 2.37E-20 | -19.63 |
| *COL5A1* | ENSG00000130635.16 | chr9:134641803-134844843 | -0.83 | -1.78 | 6.66 | 96.37 | 9.74E-23 | 6.71E-20 | -19.17 |
| *IGFBP7* | ENSG00000163453.11 | chr4:57030773-57110385 | -1.73 | -3.32 | 4.11 | 94.58 | 2.40E-22 | 1.64E-19 | -18.79 |
| *COL4A6* | ENSG00000197565.16 | chrX:108155607-108439497 | -1.18 | -2.27 | 5.24 | 94.51 | 2.48E-22 | 1.67E-19 | -18.78 |
| *ACTA2* | ENSG00000107796.13 | chr10:88935074-88991339 | -0.74 | -1.67 | 7.43 | 94.47 | 2.54E-22 | 1.69E-19 | -18.77 |
| *LAMA4* | ENSG00000112769.20 | chr6:112107931-112254939 | 1.29 | 2.44 | 4.97 | 94.37 | 2.67E-22 | 1.76E-19 | -18.76 |
| *PRSS56* | ENSG00000237412.7 | chr2:232520388-232525716 | -1.95 | -3.86 | 3.78 | 93.36 | 4.45E-22 | 2.87E-19 | -18.54 |
| *TRIM9* | ENSG00000100505.13 | chr14:50975262-51096061 | 0.76 | 1.70 | 7.14 | 93.36 | 4.44E-22 | 2.87E-19 | -18.54 |
| *COL9A3* | ENSG00000092758.18 | chr20:62816244-62841159 | -1.14 | -2.20 | 5.32 | 92.36 | 7.36E-22 | 4.70E-19 | -18.33 |
| *COL11A1* | ENSG00000060718.22 | chr1:102876467-103108872 | -0.99 | -1.98 | 5.84 | 91.92 | 9.19E-22 | 5.81E-19 | -18.24 |
| *WNT5A* | ENSG00000114251.14 | chr3:55465715-55490539 | -0.85 | -1.80 | 6.47 | 91.51 | 1.13E-21 | 7.07E-19 | -18.15 |
| *SEL1L3* | ENSG00000091490.11 | chr4:25747433-25863760 | -1.37 | -2.59 | 4.71 | 89.55 | 3.04E-21 | 1.88E-18 | -17.73 |
| *MMRN1* | ENSG00000138722.10 | chr4:89879532-89954629 | -1.89 | -3.71 | 3.77 | 87.95 | 6.82E-21 | 4.18E-18 | -17.38 |
| *FLNC* | ENSG00000128591.15 | chr7:128830377-128859274 | -0.62 | -1.54 | 8.54 | 87.03 | 1.08E-20 | 6.57E-18 | -17.18 |
| *RAB11FIP1* | ENSG00000156675.16 | chr8:37858618-37899497 | -2.20 | -4.61 | 3.29 | 85.96 | 1.87E-20 | 1.12E-17 | -16.95 |
| *ITGB8* | ENSG00000105855.10 | chr7:20330702-20415754 | -0.58 | -1.50 | 9.06 | 85.66 | 2.18E-20 | 1.29E-17 | -16.89 |
| *MFGE8* | ENSG00000140545.15 | chr15:88898683-88913381 | 0.66 | 1.58 | 7.86 | 85.65 | 2.18E-20 | 1.29E-17 | -16.89 |
| *FIGN* | ENSG00000182263.14 | chr2:163593396-163736012 | -0.92 | -1.90 | 5.95 | 84.66 | 3.59E-20 | 2.10E-17 | -16.68 |
| *MYH9* | ENSG00000100345.22 | chr22:36281280-36387967 | -0.62 | -1.54 | 8.35 | 84.03 | 4.95E-20 | 2.86E-17 | -16.54 |
| *PROX1* | ENSG00000117707.16 | chr1:213983181-214041510 | 0.88 | 1.83 | 6.10 | 82.65 | 9.93E-20 | 5.68E-17 | -16.25 |
| *SCD* | ENSG00000099194.6 | chr10:100347233-100364826 | 0.47 | 1.39 | 10.85 | 81.07 | 2.21E-19 | 1.25E-16 | -15.90 |
| *COMT* | ENSG00000093010.14 | chr22:19941733-19969975 | -2.03 | -4.08 | 3.42 | 80.11 | 3.58E-19 | 2.01E-16 | -15.70 |
| *COL4A2* | ENSG00000134871.19 | chr13:110305812-110513209 | -0.59 | -1.50 | 8.62 | 79.70 | 4.42E-19 | 2.46E-16 | -15.61 |
| *PSMD5* | ENSG00000095261.14 | chr9:120815496-120842951 | -6.52 | -91.65 | 1.25 | 79.10 | 5.98E-19 | 3.30E-16 | -15.48 |
| *ID1* | ENSG00000125968.9 | chr20:31605283-31606515 | -1.60 | -3.02 | 4.06 | 78.69 | 7.38E-19 | 4.03E-16 | -15.39 |
| *SEMA3C* | ENSG00000075223.14 | chr7:80742538-80922359 | -1.14 | -2.20 | 5.07 | 78.48 | 8.17E-19 | 4.39E-16 | -15.36 |
| *MOB3B* | ENSG00000120162.10 | chr9:27325209-27529814 | 1.05 | 2.07 | 5.28 | 78.49 | 8.13E-19 | 4.39E-16 | -15.36 |
| *MXRA5* | ENSG00000101825.8 | chrX:3308565-3346652 | -1.13 | -2.19 | 5.08 | 77.89 | 1.11E-18 | 5.88E-16 | -15.23 |
| *CRISPLD1* | ENSG00000121005.9 | chr8:74984505-75034558 | 0.62 | 1.54 | 7.92 | 77.74 | 1.19E-18 | 6.29E-16 | -15.20 |
| *MYL9* | ENSG00000101335.10 | chr20:36541497-36551447 | -0.62 | -1.53 | 8.04 | 77.52 | 1.33E-18 | 6.95E-16 | -15.16 |
| *GLUL* | ENSG00000135821.19 | chr1:182378098-182392206 | 0.60 | 1.52 | 8.26 | 77.47 | 1.37E-18 | 7.08E-16 | -15.15 |
| *PIEZO2* | ENSG00000154864.13 | chr18:10666483-11149569 | -1.41 | -2.65 | 4.36 | 77.32 | 1.47E-18 | 7.57E-16 | -15.12 |
| *SHROOM3* | ENSG00000138771.16 | chr4:76435229-76783253 | -0.71 | -1.63 | 6.97 | 76.57 | 2.15E-18 | 1.09E-15 | -14.96 |
| *COL4A5* | ENSG00000188153.14 | chrX:108439838-108697545 | -0.88 | -1.84 | 5.91 | 76.00 | 2.87E-18 | 1.44E-15 | -14.84 |
| *NPY* | ENSG00000122585.8 | chr7:24284188-24291862 | -3.33 | -10.03 | 2.16 | 74.77 | 5.36E-18 | 2.67E-15 | -14.57 |
| *FOSL2* | ENSG00000075426.12 | chr2:28392448-28417317 | -0.72 | -1.64 | 6.79 | 74.43 | 6.37E-18 | 3.09E-15 | -14.51 |
| *COL4A1* | ENSG00000187498.16 | chr13:110148963-110307157 | -0.58 | -1.49 | 8.50 | 74.43 | 6.36E-18 | 3.09E-15 | -14.51 |
| *ADAMTS9* | ENSG00000163638.13 | chr3:64515654-64688000 | -0.97 | -1.97 | 5.43 | 74.06 | 7.68E-18 | 3.70E-15 | -14.43 |
| *CXCL11* | ENSG00000169248.13 | chr4:76033682-76041415 | -10.21 | -1187.07 | 0.87 | 73.84 | 8.55E-18 | 4.08E-15 | -14.39 |
| *INPP5F* | ENSG00000198825.15 | chr10:119726042-119829147 | 0.69 | 1.61 | 6.96 | 72.88 | 1.39E-17 | 6.59E-15 | -14.18 |
| *SULF2* | ENSG00000196562.14 | chr20:47656348-47786616 | -0.68 | -1.60 | 7.07 | 71.91 | 2.27E-17 | 1.07E-14 | -13.97 |
| *ACTN1* | ENSG00000072110.14 | chr14:68874128-68979440 | -0.61 | -1.53 | 7.69 | 71.18 | 3.29E-17 | 1.54E-14 | -13.81 |
| *COL1A1* | ENSG00000108821.14 | chr17:50184101-50201632 | -0.56 | -1.47 | 8.48 | 70.08 | 5.76E-17 | 2.67E-14 | -13.57 |
| *SLC7A8* | ENSG00000092068.20 | chr14:23125295-23183674 | -1.03 | -2.04 | 5.18 | 69.71 | 6.95E-17 | 3.17E-14 | -13.50 |
| *MT1E* | ENSG00000169715.15 | chr16:56625475-56627112 | 1.12 | 2.17 | 4.91 | 69.72 | 6.92E-17 | 3.17E-14 | -13.50 |
| *IFITM2* | ENSG00000185201.16 | chr11:307631-315272 | -1.45 | -2.74 | 4.13 | 69.40 | 8.11E-17 | 3.67E-14 | -13.44 |
| *RALGAPA2* | ENSG00000188559.15 | chr20:20389530-20712644 | -0.80 | -1.74 | 6.11 | 69.31 | 8.49E-17 | 3.78E-14 | -13.42 |
| *HEG1* | ENSG00000173706.14 | chr3:124965710-125055997 | -0.63 | -1.55 | 7.45 | 69.32 | 8.46E-17 | 3.78E-14 | -13.42 |
| *COL18A1* | ENSG00000182871.16 | chr21:45405165-45513720 | -0.55 | -1.46 | 8.61 | 68.70 | 1.16E-16 | 5.12E-14 | -13.29 |
| *PCSK5* | ENSG00000099139.14 | chr9:75890644-76362975 | -1.30 | -2.46 | 4.42 | 68.50 | 1.28E-16 | 5.58E-14 | -13.25 |
| *GABRQ* | ENSG00000268089.3 | chrX:152637895-152657542 | -1.01 | -2.02 | 5.20 | 68.49 | 1.29E-16 | 5.58E-14 | -13.25 |
| *LTBP1* | ENSG00000049323.16 | chr2:32946953-33399509 | -0.75 | -1.69 | 6.36 | 68.50 | 1.28E-16 | 5.58E-14 | -13.25 |
| *PDE2A* | ENSG00000186642.16 | chr11:72576141-72674591 | 1.54 | 2.90 | 3.94 | 67.95 | 1.70E-16 | 7.30E-14 | -13.14 |
| *DHCR24* | ENSG00000116133.13 | chr1:54849627-54887195 | -0.59 | -1.50 | 7.72 | 65.76 | 5.13E-16 | 2.19E-13 | -12.66 |
| *SDC2* | ENSG00000169439.12 | chr8:96493813-96611790 | -0.63 | -1.55 | 7.20 | 64.80 | 8.35E-16 | 3.54E-13 | -12.45 |
| *FTL* | ENSG00000087086.15 | chr19:48965309-48966879 | 0.42 | 1.34 | 10.76 | 64.35 | 1.05E-15 | 4.43E-13 | -12.35 |
| *CNR1* | ENSG00000118432.12 | chr6:88139864-88166359 | 0.61 | 1.52 | 7.41 | 64.14 | 1.17E-15 | 4.90E-13 | -12.31 |
| *C7* | ENSG00000112936.19 | chr5:40909497-40984643 | -2.76 | -6.76 | 2.37 | 63.78 | 1.40E-15 | 5.82E-13 | -12.24 |
| *LMO1* | ENSG00000166407.14 | chr11:8224309-8268716 | 0.96 | 1.95 | 5.23 | 63.46 | 1.65E-15 | 6.82E-13 | -12.17 |
| *RGS5* | ENSG00000143248.13 | chr1:163111121-163321791 | -2.16 | -4.46 | 2.90 | 62.11 | 3.27E-15 | 1.33E-12 | -11.88 |
| *PODXL* | ENSG00000128567.17 | chr7:131500262-131558217 | -0.53 | -1.44 | 8.49 | 61.99 | 3.48E-15 | 1.39E-12 | -11.86 |
| *KDR* | ENSG00000128052.10 | chr4:55078481-55125595 | -2.28 | -4.84 | 2.78 | 61.94 | 3.56E-15 | 1.41E-12 | -11.85 |
| *GPC4* | ENSG00000076716.9 | chrX:133300103-133415489 | -0.61 | -1.52 | 7.31 | 61.91 | 3.63E-15 | 1.43E-12 | -11.84 |
| *GAB2* | ENSG00000033327.13 | chr11:78215293-78418348 | 0.62 | 1.54 | 7.09 | 61.85 | 3.74E-15 | 1.46E-12 | -11.84 |
| *PTGDS* | ENSG00000107317.13 | chr9:136975092-136981742 | 0.74 | 1.67 | 6.19 | 61.84 | 3.76E-15 | 1.46E-12 | -11.83 |
| *BAMBI* | ENSG00000095739.11 | chr10:28677510-28682932 | -1.04 | -2.06 | 4.95 | 61.60 | 4.25E-15 | 1.64E-12 | -11.78 |
| *LMO4* | ENSG00000143013.13 | chr1:87328880-87348923 | 0.62 | 1.53 | 7.13 | 61.47 | 4.53E-15 | 1.74E-12 | -11.76 |
| *FAT1* | ENSG00000083857.14 | chr4:186587794-186726722 | -0.48 | -1.39 | 9.31 | 61.35 | 4.83E-15 | 1.84E-12 | -11.73 |
| *PPP1R3B* | ENSG00000173281.5 | chr8:9136255-9151574 | -0.95 | -1.94 | 5.21 | 61.07 | 5.54E-15 | 2.10E-12 | -11.68 |
| *ERICH3* | ENSG00000178965.14 | chr1:74568117-74673792 | -4.10 | -17.11 | 1.53 | 61.03 | 5.67E-15 | 2.14E-12 | -11.67 |
| *MKI67* | ENSG00000148773.14 | chr10:128096659-128126423 | -0.80 | -1.74 | 5.84 | 60.83 | 6.27E-15 | 2.35E-12 | -11.63 |
| *TPM1* | ENSG00000140416.23 | chr15:63042632-63071915 | -0.65 | -1.57 | 6.70 | 60.81 | 6.34E-15 | 2.36E-12 | -11.63 |
| *CCDC80* | ENSG00000091986.16 | chr3:112596797-112649530 | -0.72 | -1.65 | 6.28 | 60.45 | 7.60E-15 | 2.81E-12 | -11.55 |
| *TFAP2B* | ENSG00000008196.13 | chr6:50818723-50847619 | 0.85 | 1.80 | 5.53 | 60.37 | 7.91E-15 | 2.91E-12 | -11.54 |
| *C1QL3* | ENSG00000165985.10 | chr10:16513734-16521879 | -1.07 | -2.10 | 4.84 | 60.09 | 9.14E-15 | 3.32E-12 | -11.48 |
| *HDAC4* | ENSG00000068024.17 | chr2:239048168-239401654 | -0.72 | -1.65 | 6.27 | 60.10 | 9.11E-15 | 3.32E-12 | -11.48 |
| *CORO2B* | ENSG00000103647.13 | chr15:68578993-68727806 | 0.55 | 1.46 | 7.89 | 60.03 | 9.40E-15 | 3.40E-12 | -11.47 |
| *IGFBP5* | ENSG00000115461.5 | chr2:216672105-216695549 | -0.54 | -1.46 | 7.99 | 59.94 | 9.87E-15 | 3.53E-12 | -11.45 |
| *TMOD1* | ENSG00000136842.14 | chr9:97501180-97601743 | 0.66 | 1.58 | 6.60 | 59.93 | 9.89E-15 | 3.53E-12 | -11.45 |
| *GNAL* | ENSG00000141404.16 | chr18:11689264-11885685 | -0.55 | -1.47 | 7.80 | 59.53 | 1.21E-14 | 4.30E-12 | -11.37 |
| *C1R* | ENSG00000159403.18 | chr12:7080214-7092540 | -0.62 | -1.54 | 6.97 | 59.00 | 1.59E-14 | 5.62E-12 | -11.25 |
| *C10orf90* | ENSG00000154493.19 | chr10:126424997-126798708 | 3.15 | 8.88 | 1.87 | 58.98 | 1.60E-14 | 5.62E-12 | -11.25 |
| *SMAD6* | ENSG00000137834.15 | chr15:66702236-66782849 | -1.42 | -2.67 | 3.97 | 58.84 | 1.72E-14 | 6.00E-12 | -11.22 |
| *PPFIA1* | ENSG00000131626.18 | chr11:70270700-70385312 | 0.59 | 1.51 | 7.27 | 58.54 | 2.00E-14 | 6.94E-12 | -11.16 |
| *EGR1* | ENSG00000120738.8 | chr5:138465479-138469303 | 0.61 | 1.53 | 6.97 | 58.19 | 2.40E-14 | 8.27E-12 | -11.08 |
| *CRIM1* | ENSG00000150938.10 | chr2:36355778-36551135 | -0.68 | -1.60 | 6.44 | 58.09 | 2.52E-14 | 8.65E-12 | -11.06 |
| *S1PR3* | ENSG00000213694.6 | chr9:88990863-89005155 | -0.61 | -1.53 | 6.97 | 57.16 | 4.05E-14 | 1.38E-11 | -10.86 |
| *ZNF703* | ENSG00000183779.7 | chr8:37695782-37700019 | 0.62 | 1.54 | 6.77 | 56.41 | 5.94E-14 | 2.01E-11 | -10.70 |
| *TENT5B* | ENSG00000158246.8 | chr1:27005020-27012850 | -1.43 | -2.70 | 3.87 | 56.34 | 6.13E-14 | 2.06E-11 | -10.69 |
| *SPTAN1* | ENSG00000197694.18 | chr9:128552558-128633662 | -0.42 | -1.34 | 10.17 | 56.13 | 6.83E-14 | 2.29E-11 | -10.64 |
| *RNF145* | ENSG00000145860.12 | chr5:159157409-159210053 | -0.66 | -1.58 | 6.47 | 55.85 | 7.86E-14 | 2.62E-11 | -10.58 |
| *IGFBP2* | ENSG00000115457.10 | chr2:216632828-216664436 | -0.50 | -1.42 | 8.37 | 55.40 | 9.89E-14 | 3.28E-11 | -10.48 |
| *TLN2* | ENSG00000171914.16 | chr15:62390526-62844631 | -0.62 | -1.54 | 6.74 | 55.20 | 1.09E-13 | 3.61E-11 | -10.44 |
| *GPR37L1* | ENSG00000170075.9 | chr1:202122917-202133592 | 0.71 | 1.63 | 6.14 | 54.94 | 1.25E-13 | 4.10E-11 | -10.39 |
| *MT-ND3* | ENSG00000198840.2 | chrM:10059-10404 | 0.37 | 1.29 | 11.32 | 54.74 | 1.39E-13 | 4.52E-11 | -10.34 |
| *NPY5R* | ENSG00000164129.12 | chr4:163343892-163351934 | -1.50 | -2.83 | 3.68 | 53.92 | 2.11E-13 | 6.83E-11 | -10.17 |
| *IFITM3* | ENSG00000142089.16 | chr11:319676-327537 | -0.73 | -1.66 | 5.96 | 53.69 | 2.37E-13 | 7.64E-11 | -10.12 |
| *GPR37* | ENSG00000170775.3 | chr7:124743885-124765792 | -1.26 | -2.39 | 4.15 | 53.20 | 3.04E-13 | 9.75E-11 | -10.01 |
| *PRDX6* | ENSG00000117592.9 | chr1:173477330-173488815 | -0.65 | -1.57 | 6.41 | 52.59 | 4.14E-13 | 1.32E-10 | -9.88 |
| *SOX8* | ENSG00000005513.10 | chr16:981770-986979 | 0.79 | 1.73 | 5.52 | 52.47 | 4.39E-13 | 1.40E-10 | -9.86 |
| *ZNF462* | ENSG00000148143.13 | chr9:106863166-107013634 | -0.54 | -1.46 | 7.46 | 52.31 | 4.77E-13 | 1.51E-10 | -9.82 |
| *CREB3L2* | ENSG00000182158.15 | chr7:137874979-138002086 | -0.74 | -1.67 | 5.84 | 52.15 | 5.17E-13 | 1.62E-10 | -9.79 |
| *MACF1* | ENSG00000127603.29 | chr1:39081316-39487177 | -0.46 | -1.37 | 8.97 | 52.08 | 5.37E-13 | 1.68E-10 | -9.77 |
| *NNMT* | ENSG00000166741.8 | chr11:114257787-114313536 | -1.51 | -2.85 | 3.60 | 52.00 | 5.58E-13 | 1.73E-10 | -9.76 |
| *SEMA3B* | ENSG00000012171.20 | chr3:50267558-50277546 | -1.01 | -2.01 | 4.79 | 51.81 | 6.14E-13 | 1.89E-10 | -9.72 |
| *CEBPZOS* | ENSG00000218739.10 | chr2:37196488-37216193 | -4.92 | -30.19 | 1.04 | 51.76 | 6.30E-13 | 1.93E-10 | -9.71 |
| *NELL2* | ENSG00000184613.11 | chr12:44508275-44921848 | -0.70 | -1.62 | 6.06 | 51.53 | 7.10E-13 | 2.16E-10 | -9.66 |
| *SERTM1* | ENSG00000180440.4 | chr13:36674020-36697839 | -1.04 | -2.06 | 4.67 | 51.39 | 7.60E-13 | 2.31E-10 | -9.64 |
| *NPHP4* | ENSG00000131697.18 | chr1:5862811-5992473 | -0.81 | -1.76 | 5.41 | 51.28 | 8.04E-13 | 2.43E-10 | -9.61 |
| *SEMA5A* | ENSG00000112902.12 | chr5:9035033-9546075 | -0.54 | -1.46 | 7.40 | 51.12 | 8.74E-13 | 2.62E-10 | -9.58 |
| *DMD* | ENSG00000198947.16 | chrX:31097677-33339441 | -0.57 | -1.48 | 7.04 | 50.98 | 9.40E-13 | 2.81E-10 | -9.55 |
| *COLEC12* | ENSG00000158270.12 | chr18:316737-500722 | -0.98 | -1.97 | 4.84 | 50.76 | 1.05E-12 | 3.12E-10 | -9.51 |
| *QPCT* | ENSG00000115828.17 | chr2:37342827-37373322 | -1.80 | -3.49 | 3.05 | 50.64 | 1.11E-12 | 3.29E-10 | -9.48 |
| *ANOS1* | ENSG00000011201.12 | chrX:8528874-8732137 | -1.28 | -2.43 | 4.03 | 50.63 | 1.12E-12 | 3.30E-10 | -9.48 |
| *EN2* | ENSG00000164778.4 | chr7:155458129-155464831 | 0.52 | 1.44 | 7.58 | 50.34 | 1.30E-12 | 3.81E-10 | -9.42 |
| *TRIL* | ENSG00000255690.3 | chr7:28953358-28958330 | 0.58 | 1.49 | 6.87 | 50.23 | 1.37E-12 | 4.00E-10 | -9.40 |
| *NEDD9* | ENSG00000111859.17 | chr6:11183298-11382348 | -0.58 | -1.49 | 6.86 | 50.02 | 1.53E-12 | 4.45E-10 | -9.35 |
| *F3* | ENSG00000117525.14 | chr1:94529173-94541759 | 0.70 | 1.62 | 5.93 | 48.83 | 2.81E-12 | 8.12E-10 | -9.09 |
| *LIMA1* | ENSG00000050405.13 | chr12:50175788-50283546 | -0.57 | -1.49 | 6.83 | 48.66 | 3.07E-12 | 8.82E-10 | -9.05 |
| *AK2* | ENSG00000004455.17 | chr1:33007940-33080996 | -1.10 | -2.14 | 4.39 | 48.46 | 3.39E-12 | 9.65E-10 | -9.02 |
| *PPDPF* | ENSG00000125534.10 | chr20:63520765-63522206 | 0.45 | 1.37 | 8.77 | 48.46 | 3.39E-12 | 9.65E-10 | -9.02 |
| *RAB3B* | ENSG00000169213.7 | chr1:51907956-51990700 | -0.51 | -1.42 | 7.64 | 48.31 | 3.66E-12 | 1.04E-09 | -8.98 |
| *SLIT1* | ENSG00000187122.17 | chr10:96998038-97185959 | 0.47 | 1.38 | 8.39 | 48.22 | 3.83E-12 | 1.08E-09 | -8.97 |
| *SP9* | ENSG00000217236.2 | chr2:174334954-174338500 | 1.39 | 2.63 | 3.69 | 48.12 | 4.03E-12 | 1.13E-09 | -8.95 |
| *ADAMTS12* | ENSG00000151388.11 | chr5:33523535-33892019 | -0.69 | -1.61 | 5.97 | 48.00 | 4.29E-12 | 1.19E-09 | -8.92 |
| *PCDHA11* | ENSG00000249158.7 | chr5:140868183-141012347 | 1.65 | 3.14 | 3.14 | 47.68 | 5.04E-12 | 1.40E-09 | -8.86 |
| *CDKN1A* | ENSG00000124762.14 | chr6:36676460-36687337 | 0.44 | 1.35 | 8.98 | 47.57 | 5.35E-12 | 1.47E-09 | -8.83 |
| *SCNN1B* | ENSG00000168447.11 | chr16:23278231-23381294 | 1.64 | 3.12 | 3.15 | 47.33 | 6.03E-12 | 1.65E-09 | -8.78 |
| *C12orf49* | ENSG00000111412.6 | chr12:116710171-116738070 | 0.54 | 1.46 | 7.08 | 47.21 | 6.40E-12 | 1.75E-09 | -8.76 |
| *MYLK* | ENSG00000065534.19 | chr3:123610049-123884332 | -0.82 | -1.77 | 5.25 | 46.61 | 8.70E-12 | 2.37E-09 | -8.63 |
| *SCARA3* | ENSG00000168077.14 | chr8:27633868-27676776 | 0.48 | 1.39 | 8.00 | 46.46 | 9.38E-12 | 2.54E-09 | -8.60 |
| *TLN1* | ENSG00000137076.21 | chr9:35696948-35732195 | -0.48 | -1.40 | 7.93 | 46.37 | 9.84E-12 | 2.65E-09 | -8.58 |
| *SOX1* | ENSG00000182968.5 | chr13:112067149-112071706 | 0.58 | 1.49 | 6.63 | 46.23 | 1.06E-11 | 2.84E-09 | -8.55 |
| *P3H2* | ENSG00000090530.10 | chr3:189956728-190122437 | -1.12 | -2.17 | 4.25 | 46.00 | 1.19E-11 | 3.17E-09 | -8.50 |
| *VCL* | ENSG00000035403.18 | chr10:73995193-74121363 | -0.50 | -1.41 | 7.67 | 45.95 | 1.22E-11 | 3.24E-09 | -8.49 |
| *ZFHX4* | ENSG00000091656.19 | chr8:76681239-76867281 | -0.68 | -1.61 | 5.89 | 45.84 | 1.29E-11 | 3.42E-09 | -8.47 |
| *SULF1* | ENSG00000137573.14 | chr8:69466624-69660915 | -1.30 | -2.47 | 3.83 | 45.67 | 1.41E-11 | 3.71E-09 | -8.43 |
| *PLXNA2* | ENSG00000076356.7 | chr1:208022242-208244384 | -0.63 | -1.55 | 6.22 | 45.66 | 1.42E-11 | 3.72E-09 | -8.43 |
| *ATP11A* | ENSG00000068650.18 | chr13:112690329-112887168 | -0.67 | -1.59 | 5.95 | 45.63 | 1.43E-11 | 3.75E-09 | -8.43 |
| *CNTNAP2* | ENSG00000174469.23 | chr7:146116002-148420998 | -0.56 | -1.47 | 6.76 | 45.57 | 1.48E-11 | 3.85E-09 | -8.41 |
| *PDE4D* | ENSG00000113448.19 | chr5:58969038-60522120 | -0.45 | -1.37 | 8.40 | 45.50 | 1.53E-11 | 3.97E-09 | -8.40 |
| *ELFN1* | ENSG00000225968.7 | chr7:1688119-1747954 | 0.61 | 1.52 | 6.36 | 45.18 | 1.81E-11 | 4.63E-09 | -8.33 |
| *CEMIP* | ENSG00000103888.17 | chr15:80779343-80951776 | -1.22 | -2.32 | 3.99 | 44.86 | 2.13E-11 | 5.42E-09 | -8.27 |
| *OLFML2A* | ENSG00000185585.20 | chr9:124777133-124814885 | -0.61 | -1.53 | 6.34 | 44.78 | 2.21E-11 | 5.62E-09 | -8.25 |
| *COL3A1* | ENSG00000168542.16 | chr2:188974373-189012746 | -0.52 | -1.44 | 7.17 | 44.76 | 2.23E-11 | 5.64E-09 | -8.25 |
| *OSBPL11* | ENSG00000144909.8 | chr3:125528858-125595497 | -0.72 | -1.64 | 5.62 | 44.67 | 2.34E-11 | 5.90E-09 | -8.23 |
| *RAPGEF1* | ENSG00000107263.18 | chr9:131576770-131740074 | 0.44 | 1.36 | 8.58 | 44.57 | 2.46E-11 | 6.17E-09 | -8.21 |
| *PLEC* | ENSG00000178209.15 | chr8:143915147-143976734 | -0.41 | -1.33 | 9.21 | 44.52 | 2.53E-11 | 6.31E-09 | -8.20 |
| *ITGA3* | ENSG00000005884.18 | chr17:50055968-50090481 | 0.59 | 1.50 | 6.45 | 44.31 | 2.82E-11 | 7.00E-09 | -8.15 |
| *GLIPR1* | ENSG00000139278.10 | chr12:75480753-75503863 | -0.70 | -1.62 | 5.72 | 44.08 | 3.16E-11 | 7.82E-09 | -8.11 |
| *TTC3* | ENSG00000182670.13 | chr21:37073226-37203112 | -0.39 | -1.31 | 9.63 | 43.87 | 3.53E-11 | 8.70E-09 | -8.06 |
| *GRID1* | ENSG00000182771.19 | chr10:85599552-86366795 | 1.19 | 2.28 | 4.01 | 43.64 | 3.97E-11 | 9.76E-09 | -8.01 |
| *ACTN4* | ENSG00000130402.12 | chr19:38647649-38731589 | -0.45 | -1.36 | 8.28 | 43.29 | 4.74E-11 | 1.16E-08 | -7.94 |
| *SCG3* | ENSG00000104112.9 | chr15:51681492-51721026 | 0.60 | 1.52 | 6.31 | 43.19 | 4.99E-11 | 1.22E-08 | -7.91 |
| *KCNF1* | ENSG00000162975.5 | chr2:10911934-10914225 | -0.74 | -1.67 | 5.43 | 43.09 | 5.25E-11 | 1.27E-08 | -7.89 |
| *PRKCA* | ENSG00000154229.12 | chr17:66302613-66810743 | -0.59 | -1.50 | 6.41 | 43.07 | 5.30E-11 | 1.28E-08 | -7.89 |
| *PLXNA4* | ENSG00000221866.9 | chr7:132123332-132648688 | -0.45 | -1.36 | 8.27 | 42.97 | 5.58E-11 | 1.34E-08 | -7.87 |
| *PON2* | ENSG00000105854.13 | chr7:95404862-95435329 | -0.43 | -1.35 | 8.58 | 42.72 | 6.33E-11 | 1.52E-08 | -7.82 |
| *PPP1R15A* | ENSG00000087074.8 | chr19:48872421-48876058 | 0.53 | 1.45 | 6.83 | 42.49 | 7.12E-11 | 1.70E-08 | -7.77 |
| *MGLL* | ENSG00000074416.14 | chr3:127689062-128052190 | 0.93 | 1.90 | 4.69 | 42.10 | 8.72E-11 | 2.07E-08 | -7.68 |
| *LHFPL3* | ENSG00000187416.12 | chr7:104328700-104907232 | 0.85 | 1.81 | 4.95 | 41.69 | 1.07E-10 | 2.54E-08 | -7.60 |
| *DEPTOR* | ENSG00000155792.10 | chr8:119873717-120050918 | 1.09 | 2.12 | 4.17 | 41.61 | 1.12E-10 | 2.65E-08 | -7.58 |
| *VAT1L* | ENSG00000171724.3 | chr16:77788564-77980107 | -0.38 | -1.30 | 9.69 | 41.54 | 1.16E-10 | 2.73E-08 | -7.56 |
| *LDLR* | ENSG00000130164.14 | chr19:11089462-11133820 | -0.62 | -1.54 | 6.09 | 41.46 | 1.21E-10 | 2.83E-08 | -7.55 |
| *PMP22* | ENSG00000109099.16 | chr17:15229773-15272292 | -0.49 | -1.41 | 7.32 | 41.34 | 1.28E-10 | 2.99E-08 | -7.52 |
| *RBFOX1* | ENSG00000078328.22 | chr16:5239802-7713340 | 1.11 | 2.16 | 4.09 | 41.17 | 1.40E-10 | 3.25E-08 | -7.49 |
| *TGFBI* | ENSG00000120708.17 | chr5:136028988-136063818 | -1.03 | -2.04 | 4.31 | 40.98 | 1.55E-10 | 3.58E-08 | -7.45 |
| *C1orf61* | ENSG00000125462.18 | chr1:156404250-156456763 | 0.47 | 1.39 | 7.61 | 40.94 | 1.58E-10 | 3.64E-08 | -7.44 |
| *ANXA2* | ENSG00000182718.17 | chr15:60347134-60402883 | -0.59 | -1.51 | 6.25 | 40.72 | 1.76E-10 | 4.05E-08 | -7.39 |
| *RPLP1* | ENSG00000137818.12 | chr15:69452814-69456205 | 0.46 | 1.37 | 7.79 | 40.46 | 2.02E-10 | 4.62E-08 | -7.34 |
| *LZTS1* | ENSG00000061337.15 | chr8:20246165-20303963 | -0.62 | -1.54 | 5.99 | 40.15 | 2.36E-10 | 5.38E-08 | -7.27 |
| *H2BC21* | ENSG00000184678.10 | chr1:149884459-149886652 | 0.51 | 1.42 | 6.96 | 40.12 | 2.40E-10 | 5.46E-08 | -7.26 |
| *COBL* | ENSG00000106078.19 | chr7:51016212-51316818 | 1.64 | 3.12 | 2.95 | 40.05 | 2.48E-10 | 5.62E-08 | -7.25 |
| *FAM20C* | ENSG00000177706.9 | chr7:192571-260772 | 0.66 | 1.58 | 5.70 | 39.99 | 2.56E-10 | 5.78E-08 | -7.24 |
| *FAM89A* | ENSG00000182118.8 | chr1:231018958-231040254 | 0.77 | 1.70 | 5.21 | 39.88 | 2.71E-10 | 6.09E-08 | -7.22 |
| *LHX2* | ENSG00000106689.11 | chr9:124001670-124033301 | 0.47 | 1.39 | 7.37 | 38.96 | 4.34E-10 | 9.68E-08 | -7.01 |
| *PID1* | ENSG00000153823.19 | chr2:228850526-229271287 | 0.65 | 1.57 | 5.71 | 38.84 | 4.61E-10 | 1.02E-07 | -6.99 |
| *RPL37A* | ENSG00000197756.10 | chr2:216498825-216579180 | 0.39 | 1.31 | 9.16 | 38.73 | 4.89E-10 | 1.08E-07 | -6.97 |
| *STON2* | ENSG00000140022.13 | chr14:81260656-81436465 | -0.62 | -1.53 | 5.97 | 38.60 | 5.21E-10 | 1.15E-07 | -6.94 |
| *HK2* | ENSG00000159399.10 | chr2:74834127-74893359 | -1.04 | -2.06 | 4.19 | 38.60 | 5.23E-10 | 1.15E-07 | -6.94 |
| *CDH11* | ENSG00000140937.14 | chr16:64943753-65126112 | -0.70 | -1.62 | 5.44 | 38.52 | 5.45E-10 | 1.19E-07 | -6.92 |
| *CSGALNACT1* | ENSG00000147408.14 | chr8:19404161-19758029 | 1.59 | 3.02 | 2.96 | 38.51 | 5.47E-10 | 1.19E-07 | -6.92 |
| *DSP* | ENSG00000096696.14 | chr6:7541617-7586714 | -3.24 | -9.45 | 1.39 | 38.50 | 5.49E-10 | 1.19E-07 | -6.92 |
| *ISLR* | ENSG00000129009.13 | chr15:74173710-74176872 | -1.08 | -2.12 | 4.09 | 38.46 | 5.60E-10 | 1.21E-07 | -6.92 |
| *LAMA2* | ENSG00000196569.13 | chr6:128883138-129516566 | -1.04 | -2.05 | 4.21 | 38.46 | 5.59E-10 | 1.21E-07 | -6.92 |
| *AHNAK2* | ENSG00000185567.7 | chr14:104937244-104978374 | -0.49 | -1.40 | 7.15 | 38.47 | 5.58E-10 | 1.21E-07 | -6.92 |
| *FBLN1* | ENSG00000077942.19 | chr22:45502238-45601135 | -0.57 | -1.48 | 6.32 | 38.37 | 5.87E-10 | 1.25E-07 | -6.90 |
| *AMOTL2* | ENSG00000114019.14 | chr3:134355874-134375479 | -0.57 | -1.48 | 6.31 | 38.28 | 6.16E-10 | 1.31E-07 | -6.88 |
| *MGAT5* | ENSG00000152127.9 | chr2:134119983-134454621 | -0.47 | -1.39 | 7.35 | 38.26 | 6.22E-10 | 1.32E-07 | -6.88 |
| *HSPA2* | ENSG00000126803.9 | chr14:64535905-64546173 | -1.65 | -3.14 | 2.89 | 37.97 | 7.21E-10 | 1.52E-07 | -6.82 |
| *SERPING1* | ENSG00000149131.16 | chr11:57597387-57614848 | -0.41 | -1.33 | 8.57 | 37.84 | 7.70E-10 | 1.62E-07 | -6.79 |
| *EDIL3* | ENSG00000164176.13 | chr5:83940554-84384880 | 0.43 | 1.35 | 7.97 | 37.73 | 8.16E-10 | 1.71E-07 | -6.77 |
| *NPY1R* | ENSG00000164128.7 | chr4:163323962-163344832 | -1.16 | -2.23 | 3.88 | 37.71 | 8.23E-10 | 1.72E-07 | -6.76 |
| *PAX8* | ENSG00000125618.17 | chr2:113215997-113278921 | -1.40 | -2.63 | 3.30 | 37.46 | 9.37E-10 | 1.95E-07 | -6.71 |
| *KIRREL3* | ENSG00000149571.12 | chr11:126423358-127003460 | -0.64 | -1.56 | 5.74 | 37.30 | 1.02E-09 | 2.11E-07 | -6.68 |
| *ZFAND5* | ENSG00000107372.13 | chr9:72351413-72365235 | 0.46 | 1.38 | 7.45 | 37.29 | 1.02E-09 | 2.12E-07 | -6.67 |
| *ESPN* | ENSG00000187017.17 | chr1:6424776-6461367 | -0.65 | -1.57 | 5.62 | 37.07 | 1.14E-09 | 2.36E-07 | -6.63 |
| *MT2A* | ENSG00000125148.7 | chr16:56608584-56609497 | 0.56 | 1.48 | 6.26 | 36.97 | 1.20E-09 | 2.47E-07 | -6.61 |
| *TRH* | ENSG00000170893.4 | chr3:129974688-129977935 | -0.56 | -1.48 | 6.27 | 36.78 | 1.33E-09 | 2.72E-07 | -6.57 |
| *ENO1* | ENSG00000074800.16 | chr1:8861000-8879190 | -0.39 | -1.31 | 8.90 | 36.71 | 1.38E-09 | 2.81E-07 | -6.55 |
| *MYO5C* | ENSG00000128833.13 | chr15:52192322-52295804 | -0.87 | -1.83 | 4.68 | 36.66 | 1.41E-09 | 2.87E-07 | -6.54 |
| *CAMK2N2* | ENSG00000163888.4 | chr3:184259213-184261553 | 0.53 | 1.44 | 6.49 | 36.61 | 1.45E-09 | 2.94E-07 | -6.53 |
| *TANGO2* | ENSG00000183597.16 | chr22:20017014-20067164 | 0.66 | 1.58 | 5.52 | 36.43 | 1.59E-09 | 3.21E-07 | -6.49 |
| *ADGRV1* | ENSG00000164199.18 | chr5:90529344-91164437 | -0.49 | -1.40 | 6.97 | 36.37 | 1.64E-09 | 3.29E-07 | -6.48 |
| *ANGPTL2* | ENSG00000136859.10 | chr9:127087348-127122635 | 0.61 | 1.53 | 5.87 | 36.36 | 1.64E-09 | 3.30E-07 | -6.48 |
| *H1-4* | ENSG00000168298.7 | chr6:26156329-26157115 | 0.41 | 1.33 | 8.23 | 36.34 | 1.67E-09 | 3.33E-07 | -6.48 |
| *CENPF* | ENSG00000117724.13 | chr1:214603195-214664571 | -0.71 | -1.63 | 5.31 | 36.30 | 1.70E-09 | 3.38E-07 | -6.47 |
| *ITGA2* | ENSG00000164171.11 | chr5:52989340-53094779 | -2.98 | -7.88 | 1.48 | 36.18 | 1.81E-09 | 3.60E-07 | -6.44 |
| *CD44* | ENSG00000026508.20 | chr11:35138882-35232402 | 0.34 | 1.27 | 9.98 | 35.95 | 2.03E-09 | 4.03E-07 | -6.39 |
| *TENT5A* | ENSG00000112773.16 | chr6:81491439-81752774 | -0.55 | -1.46 | 6.30 | 35.90 | 2.09E-09 | 4.12E-07 | -6.38 |
| *CP* | ENSG00000047457.14 | chr3:149162410-149221829 | -1.29 | -2.44 | 3.49 | 35.78 | 2.21E-09 | 4.36E-07 | -6.36 |
| *ALDOA* | ENSG00000149925.22 | chr16:30064164-30070457 | 0.64 | 1.56 | 5.57 | 35.75 | 2.25E-09 | 4.42E-07 | -6.35 |
| *KLF9* | ENSG00000119138.4 | chr9:70384597-70414624 | -0.61 | -1.52 | 5.87 | 35.72 | 2.29E-09 | 4.48E-07 | -6.35 |
| *HYOU1* | ENSG00000149428.19 | chr11:119044188-119057227 | 0.41 | 1.33 | 8.22 | 35.66 | 2.36E-09 | 4.60E-07 | -6.34 |
| *PRKDC* | ENSG00000253729.8 | chr8:47773111-47960178 | -0.45 | -1.37 | 7.42 | 35.56 | 2.48E-09 | 4.82E-07 | -6.32 |
| *NAALAD2* | ENSG00000077616.11 | chr11:90131515-90192894 | -1.37 | -2.58 | 3.29 | 35.53 | 2.52E-09 | 4.88E-07 | -6.31 |
| *TIMP3* | ENSG00000100234.12 | chr22:32801705-32863041 | -0.57 | -1.49 | 6.11 | 35.44 | 2.64E-09 | 5.10E-07 | -6.29 |
| *HSPG2* | ENSG00000142798.20 | chr1:21822244-21937310 | -0.59 | -1.51 | 5.95 | 35.26 | 2.89E-09 | 5.56E-07 | -6.26 |
| *FABP5* | ENSG00000164687.11 | chr8:81280536-81284777 | 0.50 | 1.41 | 6.71 | 35.25 | 2.91E-09 | 5.58E-07 | -6.25 |
| *UBR4* | ENSG00000127481.15 | chr1:19074510-19210266 | -0.42 | -1.34 | 7.80 | 34.91 | 3.46E-09 | 6.62E-07 | -6.18 |
| *AFF1* | ENSG00000172493.21 | chr4:86935002-87141054 | -0.52 | -1.43 | 6.45 | 34.65 | 3.95E-09 | 7.54E-07 | -6.12 |
| *MCHR1* | ENSG00000128285.4 | chr22:40678750-40682814 | -0.48 | -1.40 | 6.87 | 34.62 | 4.03E-09 | 7.65E-07 | -6.12 |
| *SLC2A12* | ENSG00000146411.6 | chr6:133987581-134052624 | -0.99 | -1.99 | 4.17 | 34.61 | 4.04E-09 | 7.67E-07 | -6.12 |
| *GLRA3* | ENSG00000145451.13 | chr4:174636920-174829247 | 1.01 | 2.01 | 4.12 | 34.42 | 4.44E-09 | 8.39E-07 | -6.08 |
| *SORBS2* | ENSG00000154556.18 | chr4:185585444-185956652 | -0.49 | -1.41 | 6.67 | 34.27 | 4.80E-09 | 9.04E-07 | -6.04 |
| *GAREM2* | ENSG00000157833.13 | chr2:26173088-26189663 | 0.46 | 1.38 | 7.14 | 34.21 | 4.96E-09 | 9.31E-07 | -6.03 |
| *MGP* | ENSG00000111341.10 | chr12:14880864-14885857 | -1.17 | -2.25 | 3.69 | 33.85 | 5.97E-09 | 1.11E-06 | -5.95 |
| *CPD* | ENSG00000108582.12 | chr17:30378927-30469989 | -0.49 | -1.40 | 6.72 | 33.85 | 5.96E-09 | 1.11E-06 | -5.95 |
| *AGPAT4* | ENSG00000026652.15 | chr6:161129967-161274061 | -0.52 | -1.43 | 6.43 | 33.81 | 6.10E-09 | 1.13E-06 | -5.95 |
| *KIF5C* | ENSG00000168280.17 | chr2:148875227-149026759 | -0.34 | -1.27 | 9.73 | 33.80 | 6.12E-09 | 1.14E-06 | -5.94 |
| *KAT2B* | ENSG00000114166.8 | chr3:20040446-20154404 | 0.50 | 1.41 | 6.55 | 33.47 | 7.24E-09 | 1.33E-06 | -5.87 |
| *PBXIP1* | ENSG00000163346.17 | chr1:154944076-154956123 | -0.45 | -1.36 | 7.28 | 33.36 | 7.68E-09 | 1.41E-06 | -5.85 |
| *HUWE1* | ENSG00000086758.16 | chrX:53532096-53686728 | -0.37 | -1.29 | 8.83 | 33.31 | 7.89E-09 | 1.45E-06 | -5.84 |
| *REC8* | ENSG00000100918.13 | chr14:24171853-24180257 | -0.90 | -1.86 | 4.42 | 33.30 | 7.92E-09 | 1.45E-06 | -5.84 |
| *GPM6A* | ENSG00000150625.16 | chr4:175632934-176002664 | 0.36 | 1.28 | 9.25 | 33.29 | 7.97E-09 | 1.45E-06 | -5.84 |
| *SF3A2* | ENSG00000104897.10 | chr19:2236824-2248655 | 0.48 | 1.40 | 6.72 | 33.28 | 8.00E-09 | 1.45E-06 | -5.84 |
| *FBN2* | ENSG00000138829.12 | chr5:128257909-128659185 | -0.73 | -1.66 | 5.09 | 33.24 | 8.19E-09 | 1.48E-06 | -5.83 |
| *DUSP4* | ENSG00000120875.9 | chr8:29333064-29350684 | 0.43 | 1.35 | 7.51 | 33.23 | 8.21E-09 | 1.48E-06 | -5.83 |
| *FSTL1* | ENSG00000163430.12 | chr3:120392293-120450993 | -0.42 | -1.34 | 7.62 | 33.13 | 8.65E-09 | 1.56E-06 | -5.81 |
| *ASCL1* | ENSG00000139352.4 | chr12:102957674-102960513 | 0.99 | 1.99 | 4.09 | 32.90 | 9.73E-09 | 1.75E-06 | -5.76 |
| *TAF9B* | ENSG00000187325.5 | chrX:78129748-78139650 | -5.23 | -37.55 | 0.44 | 32.84 | 1.00E-08 | 1.79E-06 | -5.75 |
| *LGR4* | ENSG00000205213.14 | chr11:27365961-27472790 | -0.59 | -1.51 | 5.77 | 32.77 | 1.04E-08 | 1.85E-06 | -5.73 |
| *ELAVL3* | ENSG00000196361.10 | chr19:11451326-11481046 | 0.39 | 1.31 | 8.18 | 32.66 | 1.10E-08 | 1.96E-06 | -5.71 |
| *PBX3* | ENSG00000167081.18 | chr9:125747345-125967377 | 0.44 | 1.36 | 7.27 | 32.53 | 1.17E-08 | 2.08E-06 | -5.68 |
| *TP53BP1* | ENSG00000067369.14 | chr15:43403061-43510728 | -0.42 | -1.34 | 7.56 | 32.39 | 1.26E-08 | 2.23E-06 | -5.65 |
| *RPL38* | ENSG00000172809.13 | chr17:74203582-74210655 | 0.40 | 1.32 | 7.97 | 32.39 | 1.26E-08 | 2.23E-06 | -5.65 |
| *SPECC1* | ENSG00000128487.16 | chr17:20009344-20319026 | 0.39 | 1.31 | 8.21 | 32.22 | 1.38E-08 | 2.42E-06 | -5.62 |
| *MEGF10* | ENSG00000145794.17 | chr5:127290796-127465737 | 0.42 | 1.34 | 7.53 | 32.17 | 1.42E-08 | 2.49E-06 | -5.60 |
| *CLDN6* | ENSG00000184697.7 | chr16:3014712-3020071 | -1.73 | -3.32 | 2.55 | 32.06 | 1.50E-08 | 2.61E-06 | -5.58 |
| *WWC2* | ENSG00000151718.16 | chr4:183099257-183320777 | -0.59 | -1.51 | 5.73 | 31.92 | 1.61E-08 | 2.80E-06 | -5.55 |
| *ANGPTL1* | ENSG00000116194.13 | chr1:178849535-178871077 | -1.79 | -3.47 | 2.45 | 31.79 | 1.72E-08 | 2.99E-06 | -5.52 |
| *SMG8* | ENSG00000167447.12 | chr17:59209400-59215247 | -1.39 | -2.62 | 3.06 | 31.60 | 1.90E-08 | 3.28E-06 | -5.48 |
| *GPSM2* | ENSG00000121957.15 | chr1:108875350-108934545 | 0.76 | 1.69 | 4.86 | 31.50 | 2.00E-08 | 3.45E-06 | -5.46 |
| *CBX4* | ENSG00000141582.15 | chr17:79833156-79839440 | 0.46 | 1.37 | 6.87 | 31.45 | 2.06E-08 | 3.53E-06 | -5.45 |
| *ZIC4* | ENSG00000174963.18 | chr3:147386046-147406809 | -1.15 | -2.22 | 3.63 | 31.43 | 2.08E-08 | 3.55E-06 | -5.45 |
| *GGH* | ENSG00000137563.12 | chr8:63015079-63038806 | -0.60 | -1.52 | 5.62 | 31.43 | 2.07E-08 | 3.55E-06 | -5.45 |
| *MMD2* | ENSG00000136297.14 | chr7:4905989-4959213 | 1.95 | 3.86 | 2.16 | 31.32 | 2.19E-08 | 3.73E-06 | -5.43 |
| *MYCBP2* | ENSG00000005810.18 | chr13:77044657-77327094 | -0.43 | -1.35 | 7.34 | 31.30 | 2.22E-08 | 3.76E-06 | -5.42 |
| *MMP19* | ENSG00000123342.16 | chr12:55835433-55842966 | -2.02 | -4.06 | 2.09 | 31.28 | 2.23E-08 | 3.78E-06 | -5.42 |
| *CCDC85C* | ENSG00000205476.9 | chr14:99500190-99604207 | 0.48 | 1.40 | 6.56 | 31.15 | 2.40E-08 | 4.05E-06 | -5.39 |
| *RPL27A* | ENSG00000166441.13 | chr11:8682788-8714759 | 0.38 | 1.30 | 8.29 | 30.86 | 2.77E-08 | 4.68E-06 | -5.33 |
| *ID4* | ENSG00000172201.12 | chr6:19837370-19842197 | -0.39 | -1.31 | 7.84 | 30.79 | 2.88E-08 | 4.83E-06 | -5.32 |
| *NAMPT* | ENSG00000105835.12 | chr7:106248298-106286326 | -0.44 | -1.35 | 7.14 | 30.75 | 2.94E-08 | 4.92E-06 | -5.31 |
| *PTPRG* | ENSG00000144724.20 | chr3:61561569-62297609 | -0.47 | -1.39 | 6.61 | 30.74 | 2.95E-08 | 4.93E-06 | -5.31 |
| *DOCK10* | ENSG00000135905.19 | chr2:224765090-225042445 | -0.71 | -1.64 | 5.03 | 30.69 | 3.03E-08 | 5.04E-06 | -5.30 |
| *CD34* | ENSG00000174059.17 | chr1:207880972-207911402 | -1.92 | -3.79 | 2.21 | 30.68 | 3.05E-08 | 5.07E-06 | -5.29 |
| *DSG2* | ENSG00000046604.13 | chr18:31498177-31549008 | -1.12 | -2.17 | 3.67 | 30.64 | 3.11E-08 | 5.16E-06 | -5.29 |
| *CCND1* | ENSG00000110092.4 | chr11:69641156-69654474 | -0.41 | -1.33 | 7.45 | 30.26 | 3.78E-08 | 6.24E-06 | -5.20 |
| *C21orf62* | ENSG00000205929.11 | chr21:32790673-32813743 | -0.61 | -1.53 | 5.46 | 30.19 | 3.93E-08 | 6.47E-06 | -5.19 |
| *LRRN2* | ENSG00000170382.12 | chr1:204617170-204685738 | 0.44 | 1.35 | 7.05 | 30.09 | 4.14E-08 | 6.78E-06 | -5.17 |
| *CPXM2* | ENSG00000121898.13 | chr10:123706207-123940267 | 0.47 | 1.39 | 6.54 | 30.08 | 4.15E-08 | 6.78E-06 | -5.17 |
| *NBL1* | ENSG00000158747.15 | chr1:19596979-19658456 | 0.68 | 1.60 | 5.14 | 30.07 | 4.18E-08 | 6.81E-06 | -5.17 |
| *C1GALT1* | ENSG00000106392.11 | chr7:7156934-7248616 | -0.48 | -1.40 | 6.48 | 29.90 | 4.55E-08 | 7.37E-06 | -5.13 |
| *ADGRG1* | ENSG00000205336.13 | chr16:57610652-57665580 | 0.35 | 1.27 | 8.98 | 29.90 | 4.55E-08 | 7.37E-06 | -5.13 |
| *UBE2QL1* | ENSG00000215218.4 | chr5:6448859-6496723 | 0.63 | 1.55 | 5.35 | 29.90 | 4.55E-08 | 7.37E-06 | -5.13 |
| *PRKCB* | ENSG00000166501.14 | chr16:23835983-24220611 | -0.45 | -1.36 | 6.88 | 29.85 | 4.68E-08 | 7.54E-06 | -5.12 |
| *IL1R1* | ENSG00000115594.12 | chr2:102064544-102179874 | -0.92 | -1.89 | 4.19 | 29.78 | 4.85E-08 | 7.80E-06 | -5.11 |
| *DOCK5* | ENSG00000147459.18 | chr8:25184689-25418082 | -1.00 | -2.00 | 3.95 | 29.76 | 4.91E-08 | 7.88E-06 | -5.10 |
| *ARHGAP33* | ENSG00000004777.18 | chr19:35774532-35788822 | 0.40 | 1.32 | 7.59 | 29.72 | 5.00E-08 | 8.00E-06 | -5.10 |
| *H1-10* | ENSG00000184897.6 | chr3:129314771-129316286 | 0.47 | 1.38 | 6.58 | 29.68 | 5.11E-08 | 8.16E-06 | -5.09 |
| *EDNRB* | ENSG00000136160.17 | chr13:77895481-77975529 | 0.57 | 1.49 | 5.68 | 29.63 | 5.24E-08 | 8.34E-06 | -5.08 |
| *SLIT3* | ENSG00000184347.15 | chr5:168661733-169301139 | -1.27 | -2.41 | 3.22 | 29.56 | 5.43E-08 | 8.60E-06 | -5.07 |
| *RASA3* | ENSG00000185989.11 | chr13:113977783-114132623 | -0.46 | -1.37 | 6.68 | 29.56 | 5.44E-08 | 8.60E-06 | -5.07 |
| *TNC* | ENSG00000041982.16 | chr9:115019575-115118257 | -0.36 | -1.28 | 8.56 | 29.49 | 5.62E-08 | 8.85E-06 | -5.05 |
| *OCLN* | ENSG00000197822.11 | chr5:69492292-69558104 | -1.03 | -2.05 | 3.84 | 29.40 | 5.90E-08 | 9.26E-06 | -5.03 |
| *PDLIM7* | ENSG00000196923.14 | chr5:177483394-177497606 | -0.44 | -1.35 | 6.97 | 29.40 | 5.91E-08 | 9.26E-06 | -5.03 |
| *MARVELD3* | ENSG00000140832.10 | chr16:71626161-71642114 | -2.52 | -5.73 | 1.56 | 29.39 | 5.93E-08 | 9.27E-06 | -5.03 |
| *MGAT4C* | ENSG00000182050.13 | chr12:85955666-86838904 | -0.84 | -1.79 | 4.42 | 29.21 | 6.50E-08 | 1.01E-05 | -4.99 |
| *WASL* | ENSG00000106299.8 | chr7:123681943-123749003 | 0.43 | 1.35 | 7.04 | 29.08 | 6.96E-08 | 1.08E-05 | -4.97 |
| *OPCML* | ENSG00000183715.13 | chr11:132414977-133532519 | -0.55 | -1.47 | 5.80 | 29.07 | 7.00E-08 | 1.09E-05 | -4.96 |
| *GRIK1* | ENSG00000171189.17 | chr21:29536933-29940033 | 1.12 | 2.17 | 3.57 | 28.99 | 7.29E-08 | 1.13E-05 | -4.95 |
| *BTBD11* | ENSG00000151136.15 | chr12:107318421-107659642 | -0.75 | -1.68 | 4.78 | 28.97 | 7.36E-08 | 1.13E-05 | -4.95 |
| *DYNC1H1* | ENSG00000197102.12 | chr14:101964573-102056443 | -0.33 | -1.25 | 9.41 | 28.97 | 7.36E-08 | 1.13E-05 | -4.95 |
| *PXDN* | ENSG00000130508.11 | chr2:1631887-1744852 | -0.49 | -1.41 | 6.31 | 28.86 | 7.78E-08 | 1.20E-05 | -4.92 |
| *CCN1* | ENSG00000142871.18 | chr1:85580761-85584589 | -0.40 | -1.32 | 7.47 | 28.78 | 8.13E-08 | 1.24E-05 | -4.90 |
| *MAF* | ENSG00000178573.7 | chr16:79585843-79600737 | -0.44 | -1.36 | 6.75 | 28.74 | 8.30E-08 | 1.27E-05 | -4.90 |
| *SNCB* | ENSG00000074317.11 | chr5:176620082-176630556 | 0.56 | 1.48 | 5.67 | 28.68 | 8.55E-08 | 1.30E-05 | -4.88 |
| *PDPN* | ENSG00000162493.16 | chr1:13583465-13617957 | 0.40 | 1.32 | 7.53 | 28.66 | 8.66E-08 | 1.32E-05 | -4.88 |
| *CUX1* | ENSG00000257923.11 | chr7:101815904-102283958 | -0.41 | -1.33 | 7.30 | 28.39 | 9.92E-08 | 1.50E-05 | -4.82 |
| *HSPA5* | ENSG00000044574.8 | chr9:125234853-125241343 | 0.35 | 1.27 | 8.63 | 28.38 | 1.00E-07 | 1.51E-05 | -4.82 |
| *ARHGEF40* | ENSG00000165801.10 | chr14:21070273-21090248 | -0.42 | -1.34 | 7.06 | 28.26 | 1.06E-07 | 1.60E-05 | -4.80 |
| *TPX2* | ENSG00000088325.16 | chr20:31739271-31801805 | -0.69 | -1.62 | 4.98 | 28.08 | 1.16E-07 | 1.75E-05 | -4.76 |
| *PCDHGA11* | ENSG00000253873.6 | chr5:141421047-141512975 | 1.64 | 3.11 | 2.48 | 28.05 | 1.18E-07 | 1.77E-05 | -4.75 |
| *DCBLD2* | ENSG00000057019.16 | chr3:98795941-98901695 | -0.48 | -1.40 | 6.32 | 28.03 | 1.19E-07 | 1.78E-05 | -4.75 |
| *TLR4* | ENSG00000136869.16 | chr9:117704175-117724735 | -1.18 | -2.26 | 3.38 | 28.02 | 1.21E-07 | 1.80E-05 | -4.75 |
| *ABCB1* | ENSG00000085563.15 | chr7:87503017-87713323 | -3.37 | -10.34 | 0.90 | 27.97 | 1.24E-07 | 1.84E-05 | -4.74 |
| *SEMA3A* | ENSG00000075213.11 | chr7:83955777-84492724 | -0.54 | -1.45 | 5.85 | 27.84 | 1.32E-07 | 1.96E-05 | -4.71 |
| *CST3* | ENSG00000101439.9 | chr20:23626706-23638473 | 0.36 | 1.29 | 8.17 | 27.81 | 1.34E-07 | 1.97E-05 | -4.70 |
| *CAMK2N1* | ENSG00000162545.6 | chr1:20482391-20486210 | 0.38 | 1.30 | 7.70 | 27.78 | 1.36E-07 | 2.00E-05 | -4.70 |
| *FRZB* | ENSG00000162998.5 | chr2:182833275-182866637 | -1.15 | -2.22 | 3.42 | 27.77 | 1.37E-07 | 2.01E-05 | -4.70 |
| *GOLGB1* | ENSG00000173230.15 | chr3:121663199-121749767 | -0.41 | -1.33 | 7.17 | 27.69 | 1.43E-07 | 2.09E-05 | -4.68 |
| *SERINC2* | ENSG00000168528.12 | chr1:31409565-31434680 | 0.48 | 1.40 | 6.29 | 27.66 | 1.45E-07 | 2.11E-05 | -4.67 |
| *EIF1B* | ENSG00000114784.4 | chr3:40309707-40312424 | 0.49 | 1.41 | 6.19 | 27.64 | 1.46E-07 | 2.13E-05 | -4.67 |
| *TBCK* | ENSG00000145348.17 | chr4:106041599-106316683 | -0.61 | -1.53 | 5.32 | 27.52 | 1.56E-07 | 2.26E-05 | -4.65 |
| *HEPH* | ENSG00000089472.16 | chrX:66162549-66268867 | -0.43 | -1.35 | 6.74 | 27.42 | 1.64E-07 | 2.37E-05 | -4.62 |
| *PLEKHM2* | ENSG00000116786.13 | chr1:15684320-15734769 | 0.41 | 1.33 | 7.14 | 27.34 | 1.71E-07 | 2.47E-05 | -4.61 |
| *ADGRL2* | ENSG00000117114.20 | chr1:81306147-81992436 | -0.64 | -1.56 | 5.17 | 27.32 | 1.73E-07 | 2.49E-05 | -4.60 |
| *MDN1* | ENSG00000112159.12 | chr6:89642498-89819794 | -0.55 | -1.46 | 5.68 | 27.09 | 1.94E-07 | 2.80E-05 | -4.55 |
| *ATOH8* | ENSG00000168874.13 | chr2:85751344-85791383 | -0.66 | -1.58 | 5.09 | 27.05 | 1.99E-07 | 2.86E-05 | -4.54 |
| *GRINA* | ENSG00000178719.17 | chr8:143990056-143993415 | 0.33 | 1.26 | 8.95 | 27.01 | 2.03E-07 | 2.90E-05 | -4.54 |
| *INSYN1* | ENSG00000205363.5 | chr15:73735431-73752747 | 0.53 | 1.44 | 5.83 | 27.00 | 2.04E-07 | 2.91E-05 | -4.54 |
| *RAI14* | ENSG00000039560.14 | chr5:34656328-34832612 | -0.45 | -1.37 | 6.52 | 26.90 | 2.15E-07 | 3.06E-05 | -4.51 |
| *NRCAM* | ENSG00000091129.21 | chr7:108147623-108456717 | -0.34 | -1.26 | 8.72 | 26.85 | 2.20E-07 | 3.13E-05 | -4.50 |
| *ZNF585B* | ENSG00000245680.11 | chr19:37181579-37218153 | -0.80 | -1.74 | 4.42 | 26.84 | 2.22E-07 | 3.14E-05 | -4.50 |
| *KCNJ8* | ENSG00000121361.5 | chr12:21764955-21775600 | -0.71 | -1.63 | 4.86 | 26.83 | 2.23E-07 | 3.14E-05 | -4.50 |
| *ABCA8* | ENSG00000141338.14 | chr17:68867289-68955392 | -0.56 | -1.47 | 5.56 | 26.82 | 2.23E-07 | 3.14E-05 | -4.50 |
| *FOS* | ENSG00000170345.10 | chr14:75278826-75282230 | 0.60 | 1.52 | 5.32 | 26.82 | 2.23E-07 | 3.14E-05 | -4.50 |
| *EFHD1* | ENSG00000115468.12 | chr2:232606057-232682780 | 0.78 | 1.72 | 4.50 | 26.82 | 2.24E-07 | 3.14E-05 | -4.50 |
| *FREM1* | ENSG00000164946.19 | chr9:14734666-14910995 | -1.37 | -2.59 | 2.88 | 26.69 | 2.39E-07 | 3.32E-05 | -4.48 |
| *SSPN* | ENSG00000123096.12 | chr12:26121991-26299290 | -0.47 | -1.39 | 6.30 | 26.70 | 2.38E-07 | 3.32E-05 | -4.48 |
| *SLC1A4* | ENSG00000115902.11 | chr2:64988477-65023865 | 0.43 | 1.35 | 6.66 | 26.69 | 2.39E-07 | 3.32E-05 | -4.48 |
| *SCN2A* | ENSG00000136531.18 | chr2:165194993-165392310 | -0.47 | -1.38 | 6.34 | 26.69 | 2.40E-07 | 3.33E-05 | -4.48 |
| *IQGAP1* | ENSG00000140575.13 | chr15:90388242-90502239 | -0.41 | -1.33 | 7.06 | 26.68 | 2.40E-07 | 3.33E-05 | -4.48 |
| *DIPK2A* | ENSG00000181744.9 | chr3:143971823-144048719 | -0.56 | -1.47 | 5.56 | 26.66 | 2.43E-07 | 3.36E-05 | -4.47 |
| *EVX2* | ENSG00000174279.4 | chr2:176077472-176083913 | -0.93 | -1.90 | 3.99 | 26.62 | 2.47E-07 | 3.41E-05 | -4.47 |
| *PLPP3* | ENSG00000162407.9 | chr1:56494761-56645301 | 0.42 | 1.33 | 6.97 | 26.62 | 2.48E-07 | 3.41E-05 | -4.47 |
| *UNC5C* | ENSG00000182168.15 | chr4:95162504-95549206 | -0.95 | -1.93 | 3.93 | 26.58 | 2.53E-07 | 3.47E-05 | -4.46 |
| *NTRK3* | ENSG00000140538.16 | chr15:87859751-88256768 | -0.43 | -1.34 | 6.77 | 26.47 | 2.68E-07 | 3.67E-05 | -4.43 |
| *MSN* | ENSG00000147065.17 | chrX:65588377-65741931 | -0.32 | -1.25 | 9.00 | 26.39 | 2.79E-07 | 3.81E-05 | -4.42 |
| *MAZ* | ENSG00000103495.14 | chr16:29806106-29811164 | 0.38 | 1.31 | 7.48 | 26.36 | 2.83E-07 | 3.86E-05 | -4.41 |
| *MEIS1* | ENSG00000143995.20 | chr2:66433452-66573869 | -0.74 | -1.67 | 4.64 | 26.30 | 2.93E-07 | 3.98E-05 | -4.40 |
| *CNTFR* | ENSG00000122756.15 | chr9:34551432-34590140 | 0.37 | 1.29 | 7.68 | 26.30 | 2.93E-07 | 3.98E-05 | -4.40 |
| *FAM124A* | ENSG00000150510.17 | chr13:51222334-51284239 | -0.73 | -1.66 | 4.70 | 26.15 | 3.17E-07 | 4.29E-05 | -4.37 |
| *NPTX2* | ENSG00000106236.4 | chr7:98617285-98629869 | 0.67 | 1.59 | 4.97 | 26.03 | 3.37E-07 | 4.56E-05 | -4.34 |
| *GSE1* | ENSG00000131149.19 | chr16:85169525-85676204 | 0.42 | 1.34 | 6.74 | 25.96 | 3.49E-07 | 4.71E-05 | -4.33 |
| *S100A10* | ENSG00000197747.9 | chr1:151982915-151993859 | -0.47 | -1.39 | 6.24 | 25.92 | 3.56E-07 | 4.79E-05 | -4.32 |
| *NEBL* | ENSG00000078114.19 | chr10:20779973-21293011 | -0.41 | -1.33 | 6.96 | 25.90 | 3.60E-07 | 4.83E-05 | -4.32 |
| *KCTD13* | ENSG00000174943.11 | chr16:29905012-29926236 | 0.46 | 1.38 | 6.32 | 25.86 | 3.68E-07 | 4.93E-05 | -4.31 |
| *TLE5* | ENSG00000104964.15 | chr19:3052910-3063107 | 0.38 | 1.30 | 7.55 | 25.75 | 3.90E-07 | 5.21E-05 | -4.28 |
| *CSPG5* | ENSG00000114646.10 | chr3:47562238-47580792 | 0.39 | 1.31 | 7.35 | 25.71 | 3.97E-07 | 5.29E-05 | -4.28 |
| *ASPM* | ENSG00000066279.18 | chr1:197084127-197146694 | -1.17 | -2.25 | 3.23 | 25.66 | 4.08E-07 | 5.43E-05 | -4.27 |
| *ATG12* | ENSG00000145782.13 | chr5:115828200-115841837 | 0.53 | 1.44 | 5.72 | 25.50 | 4.43E-07 | 5.89E-05 | -4.23 |
| *LHX9* | ENSG00000143355.16 | chr1:197911902-197935478 | -0.52 | -1.43 | 5.82 | 25.44 | 4.57E-07 | 6.06E-05 | -4.22 |
| *PCDH10* | ENSG00000138650.9 | chr4:133149294-133208606 | 0.36 | 1.28 | 7.84 | 25.42 | 4.61E-07 | 6.09E-05 | -4.22 |
| *VCAN* | ENSG00000038427.16 | chr5:83471618-83582303 | -0.32 | -1.25 | 8.80 | 25.40 | 4.67E-07 | 6.16E-05 | -4.21 |
| *SYNE3* | ENSG00000176438.12 | chr14:95407266-95475836 | 0.94 | 1.92 | 3.86 | 25.39 | 4.68E-07 | 6.16E-05 | -4.21 |
| *KCNIP3* | ENSG00000115041.13 | chr2:95297327-95386077 | 0.71 | 1.64 | 4.72 | 25.30 | 4.91E-07 | 6.44E-05 | -4.19 |
| *TENM4* | ENSG00000149256.16 | chr11:78652829-79441030 | -0.40 | -1.32 | 7.12 | 25.22 | 5.12E-07 | 6.70E-05 | -4.17 |
| *ADAMTS1* | ENSG00000154734.15 | chr21:26835755-26845409 | -1.21 | -2.32 | 3.11 | 25.21 | 5.15E-07 | 6.72E-05 | -4.17 |
| *SDK1* | ENSG00000146555.19 | chr7:3301252-4269000 | -0.32 | -1.25 | 8.81 | 25.12 | 5.40E-07 | 7.03E-05 | -4.15 |
| *IMPA2* | ENSG00000141401.12 | chr18:11981025-12030877 | -1.59 | -3.02 | 2.42 | 25.03 | 5.66E-07 | 7.33E-05 | -4.13 |
| *TPPP3* | ENSG00000159713.11 | chr16:67389809-67393518 | -0.62 | -1.54 | 5.14 | 25.03 | 5.66E-07 | 7.33E-05 | -4.13 |
| *AMOTL1* | ENSG00000166025.18 | chr11:94706431-94876748 | -0.40 | -1.32 | 7.08 | 24.93 | 5.95E-07 | 7.69E-05 | -4.11 |
| *HSPB1* | ENSG00000106211.10 | chr7:76302673-76304295 | 0.35 | 1.27 | 8.06 | 24.92 | 5.98E-07 | 7.72E-05 | -4.11 |
| *S100A11* | ENSG00000163191.6 | chr1:152032506-152047907 | -0.54 | -1.46 | 5.53 | 24.90 | 6.05E-07 | 7.79E-05 | -4.11 |
| *FAT4* | ENSG00000196159.13 | chr4:125314955-125492932 | -0.53 | -1.44 | 5.67 | 24.89 | 6.06E-07 | 7.79E-05 | -4.11 |
| *RPL30* | ENSG00000156482.11 | chr8:98024851-98046469 | 0.35 | 1.27 | 7.97 | 24.83 | 6.25E-07 | 8.02E-05 | -4.10 |
| *AGAP3* | ENSG00000133612.19 | chr7:151085831-151144436 | 0.41 | 1.33 | 6.86 | 24.81 | 6.32E-07 | 8.08E-05 | -4.09 |
| *NET1* | ENSG00000173848.19 | chr10:5412557-5459056 | 0.58 | 1.50 | 5.30 | 24.81 | 6.32E-07 | 8.08E-05 | -4.09 |
| *PAK3* | ENSG00000077264.15 | chrX:110944285-111227361 | -0.37 | -1.29 | 7.58 | 24.75 | 6.55E-07 | 8.35E-05 | -4.08 |
| *ILDR2* | ENSG00000143195.13 | chr1:166895711-166975540 | -0.56 | -1.48 | 5.41 | 24.72 | 6.65E-07 | 8.46E-05 | -4.07 |
| *HS6ST2* | ENSG00000171004.18 | chrX:132626016-132961395 | 0.60 | 1.52 | 5.20 | 24.62 | 6.98E-07 | 8.86E-05 | -4.05 |
| *XKR4* | ENSG00000206579.9 | chr8:55102028-55542054 | -0.49 | -1.40 | 6.01 | 24.61 | 7.04E-07 | 8.92E-05 | -4.05 |
| *FZD1* | ENSG00000157240.4 | chr7:91264433-91271326 | -0.50 | -1.41 | 5.87 | 24.43 | 7.73E-07 | 9.75E-05 | -4.01 |
| *DHRS3* | ENSG00000162496.9 | chr1:12567910-12618210 | 0.59 | 1.51 | 5.23 | 24.39 | 7.86E-07 | 9.89E-05 | -4.00 |
| *MBNL2* | ENSG00000139793.18 | chr13:97221434-97394120 | -0.48 | -1.40 | 6.00 | 24.36 | 7.99E-07 | 1.00E-04 | -4.00 |
| *EFNB3* | ENSG00000108947.5 | chr17:7705202-7711372 | -0.36 | -1.28 | 7.65 | 24.36 | 7.99E-07 | 1.00E-04 | -4.00 |
| *DUSP1* | ENSG00000120129.6 | chr5:172768096-172771195 | -0.76 | -1.69 | 4.42 | 24.34 | 8.07E-07 | 1.01E-04 | -4.00 |
| *TMEM158* | ENSG00000249992.2 | chr3:45224466-45226287 | -0.41 | -1.33 | 6.73 | 24.34 | 8.07E-07 | 1.01E-04 | -4.00 |
| *IRS2* | ENSG00000185950.9 | chr13:109752695-109786583 | 0.34 | 1.26 | 8.25 | 24.27 | 8.40E-07 | 1.04E-04 | -3.98 |
| *GAD1* | ENSG00000128683.14 | chr2:170813213-170861151 | 0.66 | 1.58 | 4.89 | 24.26 | 8.44E-07 | 1.05E-04 | -3.98 |
| *RAP1GAP* | ENSG00000076864.19 | chr1:21596215-21669363 | 0.51 | 1.43 | 5.72 | 24.20 | 8.68E-07 | 1.07E-04 | -3.97 |
| *ITIH5* | ENSG00000123243.15 | chr10:7559270-7666998 | -0.52 | -1.44 | 5.63 | 24.15 | 8.94E-07 | 1.10E-04 | -3.96 |
| *SLC2A13* | ENSG00000151229.13 | chr12:39755025-40106089 | -0.55 | -1.47 | 5.43 | 24.06 | 9.34E-07 | 1.15E-04 | -3.94 |
| *TJP1* | ENSG00000104067.16 | chr15:29699367-29968865 | -0.34 | -1.27 | 8.08 | 24.01 | 9.58E-07 | 1.18E-04 | -3.93 |
| *ZFP28* | ENSG00000196867.8 | chr19:56538948-56556808 | -0.89 | -1.85 | 3.98 | 24.01 | 9.62E-07 | 1.18E-04 | -3.93 |
| *BTG1* | ENSG00000133639.6 | chr12:92140278-92145846 | 0.51 | 1.42 | 5.75 | 23.99 | 9.71E-07 | 1.19E-04 | -3.93 |
| *PRKAR1B* | ENSG00000188191.15 | chr7:549197-727650 | 0.51 | 1.43 | 5.70 | 23.88 | 1.03E-06 | 1.25E-04 | -3.90 |
| *PCSK1* | ENSG00000175426.11 | chr5:96390333-96434143 | -0.53 | -1.44 | 5.53 | 23.84 | 1.05E-06 | 1.28E-04 | -3.89 |
| *ZNF274* | ENSG00000171606.18 | chr19:58183029-58213562 | 0.65 | 1.56 | 4.93 | 23.75 | 1.10E-06 | 1.33E-04 | -3.88 |
| *RFX4* | ENSG00000111783.13 | chr12:106583004-106762803 | -0.37 | -1.29 | 7.34 | 23.65 | 1.16E-06 | 1.40E-04 | -3.85 |
| *GABRB2* | ENSG00000145864.14 | chr5:161288429-161549044 | -0.76 | -1.69 | 4.38 | 23.54 | 1.22E-06 | 1.48E-04 | -3.83 |
| *OLFML3* | ENSG00000116774.12 | chr1:113979391-114035572 | -0.91 | -1.88 | 3.86 | 23.45 | 1.28E-06 | 1.54E-04 | -3.81 |
| *RHOJ* | ENSG00000126785.13 | chr14:63204114-63293508 | -0.84 | -1.80 | 4.08 | 23.42 | 1.31E-06 | 1.57E-04 | -3.80 |
| *MRTFB* | ENSG00000186260.17 | chr16:14071319-14266773 | -0.44 | -1.36 | 6.29 | 23.38 | 1.33E-06 | 1.60E-04 | -3.80 |
| *NCAN* | ENSG00000130287.14 | chr19:19211958-19252233 | 0.33 | 1.26 | 8.25 | 23.37 | 1.34E-06 | 1.60E-04 | -3.80 |
| *JMJD8* | ENSG00000161999.13 | chr16:681670-684528 | -0.49 | -1.41 | 5.84 | 23.35 | 1.35E-06 | 1.62E-04 | -3.79 |
| *WFS1* | ENSG00000109501.14 | chr4:6269849-6303265 | 0.40 | 1.32 | 6.77 | 23.25 | 1.42E-06 | 1.69E-04 | -3.77 |
| *HMGA1* | ENSG00000137309.20 | chr6:34236873-34246231 | 0.33 | 1.26 | 8.22 | 23.03 | 1.60E-06 | 1.89E-04 | -3.72 |
| *LAMB1* | ENSG00000091136.14 | chr7:107923799-108003187 | -0.65 | -1.57 | 4.86 | 23.02 | 1.61E-06 | 1.90E-04 | -3.72 |
| *PDE10A* | ENSG00000112541.16 | chr6:165327287-165988078 | -0.36 | -1.28 | 7.52 | 23.00 | 1.62E-06 | 1.91E-04 | -3.72 |
| *CFI* | ENSG00000205403.13 | chr4:109740694-109802179 | -0.78 | -1.72 | 4.26 | 22.97 | 1.64E-06 | 1.93E-04 | -3.71 |
| *RNF152* | ENSG00000176641.11 | chr18:61808067-61894247 | -0.41 | -1.33 | 6.59 | 22.97 | 1.65E-06 | 1.93E-04 | -3.71 |
| *IGF2* | ENSG00000167244.21 | chr11:2129112-2141238 | 1.77 | 3.42 | 1.97 | 22.91 | 1.70E-06 | 1.99E-04 | -3.70 |
| *CAMSAP3* | ENSG00000076826.10 | chr19:7595863-7618304 | 0.46 | 1.37 | 6.13 | 22.88 | 1.72E-06 | 2.02E-04 | -3.69 |
| *PRSS23* | ENSG00000150687.12 | chr11:86791059-86952910 | -0.43 | -1.35 | 6.35 | 22.81 | 1.79E-06 | 2.09E-04 | -3.68 |
| *NPVF* | ENSG00000105954.2 | chr7:25224570-25228486 | -1.20 | -2.29 | 3.01 | 22.66 | 1.94E-06 | 2.26E-04 | -3.65 |
| *SLC17A6* | ENSG00000091664.9 | chr11:22338381-22379503 | -0.37 | -1.29 | 7.19 | 22.63 | 1.96E-06 | 2.29E-04 | -3.64 |
| *PDE7B* | ENSG00000171408.14 | chr6:135851701-136195574 | -0.73 | -1.66 | 4.42 | 22.62 | 1.98E-06 | 2.29E-04 | -3.64 |
| *PLEKHA5* | ENSG00000052126.14 | chr12:19129752-19376400 | -0.40 | -1.32 | 6.65 | 22.62 | 1.98E-06 | 2.29E-04 | -3.64 |
| *HLA-E* | ENSG00000204592.9 | chr6:30489509-30494194 | 0.34 | 1.27 | 7.75 | 22.62 | 1.98E-06 | 2.29E-04 | -3.64 |
| *GRAMD1A* | ENSG00000089351.14 | chr19:34994784-35026471 | 0.35 | 1.28 | 7.54 | 22.63 | 1.97E-06 | 2.29E-04 | -3.64 |
| *DPY19L1* | ENSG00000173852.15 | chr7:34928876-35038271 | 0.49 | 1.41 | 5.76 | 22.53 | 2.07E-06 | 2.39E-04 | -3.62 |
| *LTBP2* | ENSG00000119681.12 | chr14:74498183-74612378 | -0.85 | -1.80 | 4.02 | 22.53 | 2.08E-06 | 2.39E-04 | -3.62 |
| *BACE1* | ENSG00000186318.16 | chr11:117285207-117316259 | 0.36 | 1.28 | 7.43 | 22.52 | 2.08E-06 | 2.39E-04 | -3.62 |
| *ABHD4* | ENSG00000100439.10 | chr14:22598237-22613215 | 0.35 | 1.27 | 7.63 | 22.48 | 2.13E-06 | 2.44E-04 | -3.61 |
| *RPL21* | ENSG00000122026.10 | chr13:27251309-27256691 | 0.35 | 1.27 | 7.57 | 22.45 | 2.15E-06 | 2.46E-04 | -3.61 |
| *ITPK1* | ENSG00000100605.17 | chr14:92936914-93116320 | 0.43 | 1.34 | 6.38 | 22.46 | 2.15E-06 | 2.46E-04 | -3.61 |
| *CSRP1* | ENSG00000159176.14 | chr1:201483530-201509456 | -0.44 | -1.35 | 6.29 | 22.44 | 2.17E-06 | 2.47E-04 | -3.61 |
| *LRBA* | ENSG00000198589.14 | chr4:150264435-151015727 | -0.51 | -1.43 | 5.56 | 22.43 | 2.19E-06 | 2.48E-04 | -3.60 |
| *MID1* | ENSG00000101871.16 | chrX:10445310-10833654 | 0.36 | 1.28 | 7.40 | 22.41 | 2.21E-06 | 2.50E-04 | -3.60 |
| *CRABP2* | ENSG00000143320.9 | chr1:156699606-156705816 | -0.71 | -1.64 | 4.51 | 22.39 | 2.23E-06 | 2.52E-04 | -3.60 |
| *FAM117A* | ENSG00000121104.8 | chr17:49710332-49789180 | -0.60 | -1.51 | 5.09 | 22.32 | 2.32E-06 | 2.61E-04 | -3.58 |
| *MARK4* | ENSG00000007047.15 | chr19:45079288-45305284 | 0.34 | 1.27 | 7.71 | 22.27 | 2.37E-06 | 2.67E-04 | -3.57 |
| *TTYH2* | ENSG00000141540.11 | chr17:74213571-74262020 | 0.46 | 1.38 | 6.01 | 22.24 | 2.41E-06 | 2.71E-04 | -3.57 |
| *TBX2* | ENSG00000121068.14 | chr17:61399843-61409466 | 1.06 | 2.09 | 3.29 | 22.18 | 2.48E-06 | 2.78E-04 | -3.56 |
| *PLCH1* | ENSG00000114805.18 | chr3:155375580-155745071 | -0.56 | -1.48 | 5.25 | 22.14 | 2.54E-06 | 2.85E-04 | -3.55 |
| *PRDM16* | ENSG00000142611.17 | chr1:3069168-3438621 | 0.57 | 1.49 | 5.18 | 22.09 | 2.61E-06 | 2.91E-04 | -3.54 |
| *AKAP9* | ENSG00000127914.17 | chr7:91940862-92110673 | -0.38 | -1.30 | 6.88 | 22.08 | 2.61E-06 | 2.91E-04 | -3.54 |
| *HS3ST4* | ENSG00000182601.7 | chr16:25691959-26137685 | 8.68 | 411.10 | -0.65 | 22.08 | 2.62E-06 | 2.91E-04 | -3.54 |
| *PXDNL* | ENSG00000147485.13 | chr8:51319577-51809445 | -1.26 | -2.39 | 2.86 | 22.07 | 2.63E-06 | 2.91E-04 | -3.54 |
| *SPHKAP* | ENSG00000153820.13 | chr2:227979955-228181687 | 0.98 | 1.97 | 3.55 | 22.03 | 2.69E-06 | 2.97E-04 | -3.53 |
| *KCTD12* | ENSG00000178695.6 | chr13:76880175-76886405 | -0.38 | -1.30 | 6.96 | 21.96 | 2.78E-06 | 3.08E-04 | -3.51 |
| *HIVEP2* | ENSG00000010818.10 | chr6:142751469-142956698 | -0.39 | -1.31 | 6.80 | 21.86 | 2.94E-06 | 3.24E-04 | -3.49 |
| *GIT1* | ENSG00000108262.16 | chr17:29573475-29594054 | 0.35 | 1.27 | 7.53 | 21.85 | 2.95E-06 | 3.25E-04 | -3.49 |
| *NAT8L* | ENSG00000185818.8 | chr4:2059327-2069089 | 0.37 | 1.29 | 7.12 | 21.84 | 2.97E-06 | 3.26E-04 | -3.49 |
| *MCRIP1* | ENSG00000225663.8 | chr17:81822361-81833302 | 0.43 | 1.35 | 6.27 | 21.84 | 2.97E-06 | 3.26E-04 | -3.49 |
| *TSC22D1* | ENSG00000102804.15 | chr13:44432143-44577147 | 0.33 | 1.26 | 7.93 | 21.82 | 3.00E-06 | 3.29E-04 | -3.48 |
| *GLIS2* | ENSG00000126603.9 | chr16:4314761-4339597 | 0.37 | 1.29 | 7.14 | 21.80 | 3.03E-06 | 3.31E-04 | -3.48 |
| *NBEA* | ENSG00000172915.18 | chr13:34942287-35673022 | -0.35 | -1.27 | 7.48 | 21.72 | 3.16E-06 | 3.44E-04 | -3.46 |
| *STK38L* | ENSG00000211455.8 | chr12:27243968-27325959 | -0.60 | -1.51 | 5.04 | 21.70 | 3.19E-06 | 3.48E-04 | -3.46 |
| *BRSK1* | ENSG00000160469.17 | chr19:55282072-55312562 | 0.38 | 1.30 | 6.88 | 21.50 | 3.55E-06 | 3.85E-04 | -3.41 |
| *RPRML* | ENSG00000179673.5 | chr17:46978156-46979253 | -2.25 | -4.76 | 1.42 | 21.46 | 3.61E-06 | 3.92E-04 | -3.41 |
| *PLPPR4* | ENSG00000117600.12 | chr1:99263953-99309590 | 0.43 | 1.35 | 6.20 | 21.36 | 3.81E-06 | 4.12E-04 | -3.39 |
| *CUBN* | ENSG00000107611.16 | chr10:16823966-17129811 | -1.07 | -2.10 | 3.22 | 21.35 | 3.84E-06 | 4.14E-04 | -3.38 |
| *COX6B1* | ENSG00000126267.11 | chr19:35648323-35658782 | 0.37 | 1.29 | 6.98 | 21.22 | 4.09E-06 | 4.40E-04 | -3.36 |
| *JADE1* | ENSG00000077684.16 | chr4:128809700-128875224 | -0.65 | -1.57 | 4.75 | 21.21 | 4.11E-06 | 4.41E-04 | -3.36 |
| *SYT5* | ENSG00000129990.15 | chr19:55171196-55180289 | 0.38 | 1.30 | 6.82 | 21.19 | 4.17E-06 | 4.47E-04 | -3.35 |
| *PTPN14* | ENSG00000152104.12 | chr1:214348700-214552449 | -0.58 | -1.49 | 5.11 | 21.18 | 4.19E-06 | 4.48E-04 | -3.35 |
| *MYL6B* | ENSG00000196465.10 | chr12:56152256-56159647 | 0.47 | 1.38 | 5.81 | 21.09 | 4.39E-06 | 4.68E-04 | -3.33 |
| *C2orf72* | ENSG00000204128.6 | chr2:231037523-231049719 | 0.62 | 1.53 | 4.87 | 21.06 | 4.45E-06 | 4.72E-04 | -3.33 |
| *SORCS2* | ENSG00000184985.16 | chr4:7192538-7742836 | 0.36 | 1.28 | 7.14 | 21.04 | 4.50E-06 | 4.77E-04 | -3.32 |
| *SERPINF1* | ENSG00000132386.11 | chr17:1762029-1777565 | -0.90 | -1.86 | 3.75 | 21.02 | 4.55E-06 | 4.81E-04 | -3.32 |
| *TUBA1C* | ENSG00000167553.16 | chr12:49188736-49274600 | -0.53 | -1.44 | 5.36 | 20.99 | 4.62E-06 | 4.88E-04 | -3.31 |
| *EFNA5* | ENSG00000184349.13 | chr5:107376889-107670937 | -0.42 | -1.34 | 6.27 | 20.95 | 4.71E-06 | 4.97E-04 | -3.30 |
| *CTDNEP1* | ENSG00000175826.12 | chr17:7243591-7252491 | 0.37 | 1.29 | 6.93 | 20.95 | 4.72E-06 | 4.97E-04 | -3.30 |
| *THSD7A* | ENSG00000005108.16 | chr7:11370365-11832198 | -0.55 | -1.46 | 5.25 | 20.91 | 4.82E-06 | 5.06E-04 | -3.30 |
| *RAB3A* | ENSG00000105649.10 | chr19:18196784-18204042 | 0.34 | 1.26 | 7.62 | 20.85 | 4.97E-06 | 5.21E-04 | -3.28 |
| *ZSWIM6* | ENSG00000130449.6 | chr5:61332258-61546172 | -0.37 | -1.29 | 6.93 | 20.82 | 5.06E-06 | 5.28E-04 | -3.28 |
| *DDR2* | ENSG00000162733.19 | chr1:162631373-162787405 | -0.48 | -1.40 | 5.67 | 20.81 | 5.09E-06 | 5.30E-04 | -3.28 |
| *RAB12* | ENSG00000206418.4 | chr18:8609437-8639382 | 0.52 | 1.43 | 5.37 | 20.73 | 5.28E-06 | 5.48E-04 | -3.26 |
| *CHSY1* | ENSG00000131873.7 | chr15:101175727-101252048 | -0.42 | -1.33 | 6.31 | 20.70 | 5.38E-06 | 5.58E-04 | -3.25 |
| *PDZRN3* | ENSG00000121440.15 | chr3:73382431-73624941 | -0.48 | -1.39 | 5.69 | 20.69 | 5.40E-06 | 5.59E-04 | -3.25 |
| *SYNE1* | ENSG00000131018.24 | chr6:152121684-152637801 | -0.38 | -1.30 | 6.71 | 20.69 | 5.41E-06 | 5.59E-04 | -3.25 |
| *MAB21L1* | ENSG00000180660.8 | chr13:35473789-35476689 | -0.51 | -1.43 | 5.42 | 20.63 | 5.58E-06 | 5.76E-04 | -3.24 |
| *ITPRID2* | ENSG00000138434.17 | chr2:181891730-181930738 | 0.38 | 1.30 | 6.67 | 20.57 | 5.76E-06 | 5.94E-04 | -3.23 |
| *GALNT10* | ENSG00000164574.16 | chr5:154190730-154420984 | -0.42 | -1.34 | 6.22 | 20.45 | 6.11E-06 | 6.27E-04 | -3.20 |
| *PODXL2* | ENSG00000114631.11 | chr3:127629185-127672802 | 0.33 | 1.25 | 7.76 | 20.45 | 6.14E-06 | 6.29E-04 | -3.20 |
| *CNN1* | ENSG00000130176.8 | chr19:11538767-11550323 | -0.71 | -1.64 | 4.33 | 20.43 | 6.18E-06 | 6.31E-04 | -3.20 |
| *BCAR1* | ENSG00000050820.17 | chr16:75228181-75268053 | 0.35 | 1.27 | 7.25 | 20.43 | 6.18E-06 | 6.31E-04 | -3.20 |
| *MYO5B* | ENSG00000167306.20 | chr18:49822789-50195147 | -0.72 | -1.65 | 4.30 | 20.39 | 6.33E-06 | 6.43E-04 | -3.19 |
| *BIRC6* | ENSG00000115760.14 | chr2:32357028-32618899 | -0.37 | -1.29 | 6.92 | 20.35 | 6.45E-06 | 6.55E-04 | -3.18 |
| *PLEKHF1* | ENSG00000166289.6 | chr19:29665459-29675477 | 1.55 | 2.93 | 2.13 | 20.32 | 6.57E-06 | 6.65E-04 | -3.18 |
| *ATP1A3* | ENSG00000105409.19 | chr19:41966582-41997497 | 0.47 | 1.39 | 5.71 | 20.31 | 6.59E-06 | 6.66E-04 | -3.18 |
| *AP1S2* | ENSG00000182287.15 | chrX:15825806-15854931 | 0.37 | 1.29 | 6.82 | 20.30 | 6.62E-06 | 6.68E-04 | -3.18 |
| *PCDH9* | ENSG00000184226.15 | chr13:66302834-67230445 | -0.33 | -1.25 | 7.71 | 20.29 | 6.66E-06 | 6.71E-04 | -3.17 |
| *B3GAT2* | ENSG00000112309.11 | chr6:70856679-70957060 | -0.67 | -1.59 | 4.57 | 20.24 | 6.84E-06 | 6.87E-04 | -3.16 |
| *WSCD2* | ENSG00000075035.10 | chr12:108129288-108250537 | -0.85 | -1.80 | 3.85 | 20.11 | 7.32E-06 | 7.34E-04 | -3.13 |
| *SCN4B* | ENSG00000177098.9 | chr11:118133377-118152888 | 0.36 | 1.29 | 6.91 | 20.08 | 7.42E-06 | 7.42E-04 | -3.13 |
| *COL2A1* | ENSG00000139219.19 | chr12:47972967-48004554 | -1.65 | -3.14 | 1.99 | 20.04 | 7.58E-06 | 7.58E-04 | -3.12 |
| *ARHGAP28* | ENSG00000088756.13 | chr18:6729716-6915716 | -1.16 | -2.23 | 2.94 | 20.03 | 7.63E-06 | 7.60E-04 | -3.12 |
| *HYDIN* | ENSG00000157423.18 | chr16:70802084-71230722 | -0.59 | -1.51 | 4.93 | 20.03 | 7.62E-06 | 7.60E-04 | -3.12 |
| *UBE2D2* | ENSG00000131508.16 | chr5:139526431-139628434 | 0.36 | 1.28 | 6.94 | 19.99 | 7.78E-06 | 7.74E-04 | -3.11 |
| *HEPACAM* | ENSG00000165478.7 | chr11:124919205-124936412 | 0.37 | 1.29 | 6.79 | 19.98 | 7.82E-06 | 7.76E-04 | -3.11 |
| *CEMIP2* | ENSG00000135048.14 | chr9:71683366-71816690 | -0.60 | -1.52 | 4.87 | 19.95 | 7.95E-06 | 7.85E-04 | -3.11 |
| *LAMC1* | ENSG00000135862.6 | chr1:183023420-183145592 | -0.34 | -1.27 | 7.28 | 19.95 | 7.95E-06 | 7.85E-04 | -3.11 |
| *ZIC5* | ENSG00000139800.8 | chr13:99962964-99971909 | -0.59 | -1.50 | 4.96 | 19.93 | 8.04E-06 | 7.93E-04 | -3.10 |
| *LIX1* | ENSG00000145721.12 | chr5:97091867-97142753 | -0.36 | -1.28 | 6.92 | 19.91 | 8.12E-06 | 8.00E-04 | -3.10 |
| *MDGA1* | ENSG00000112139.16 | chr6:37630679-37699306 | -0.34 | -1.26 | 7.37 | 19.89 | 8.21E-06 | 8.07E-04 | -3.09 |
| *SZRD1* | ENSG00000055070.17 | chr1:16352575-16398145 | 0.36 | 1.28 | 6.90 | 19.82 | 8.50E-06 | 8.33E-04 | -3.08 |
| *EHD4* | ENSG00000103966.11 | chr15:41895933-41972557 | -0.89 | -1.85 | 3.68 | 19.77 | 8.76E-06 | 8.55E-04 | -3.07 |
| *SYNJ2* | ENSG00000078269.15 | chr6:157981863-158099176 | -0.49 | -1.41 | 5.45 | 19.68 | 9.16E-06 | 8.93E-04 | -3.05 |
| *ATAT1* | ENSG00000137343.18 | chr6:30626842-30646823 | 0.33 | 1.25 | 7.59 | 19.66 | 9.28E-06 | 9.03E-04 | -3.04 |
| *CLEC19A* | ENSG00000261210.8 | chr16:19285731-19322145 | -0.86 | -1.82 | 3.76 | 19.56 | 9.75E-06 | 9.47E-04 | -3.02 |
| *KCND3* | ENSG00000171385.9 | chr1:111770662-111989155 | 0.36 | 1.29 | 6.81 | 19.53 | 9.92E-06 | 9.61E-04 | -3.02 |
| *NHLH2* | ENSG00000177551.5 | chr1:115836377-115843917 | -0.92 | -1.90 | 3.55 | 19.52 | 9.98E-06 | 9.66E-04 | -3.02 |
| *UACA* | ENSG00000137831.15 | chr15:70654554-70763558 | -0.63 | -1.55 | 4.69 | 19.47 | 1.02E-05 | 9.90E-04 | -3.00 |
| *H2AC20* | ENSG00000184260.6 | chr1:149886918-149887411 | 0.44 | 1.36 | 5.92 | 19.46 | 1.03E-05 | 9.93E-04 | -3.00 |
| *FBN1* | ENSG00000166147.14 | chr15:48408313-48645721 | -0.80 | -1.74 | 3.97 | 19.43 | 1.04E-05 | 1.00E-03 | -3.00 |
| *TMEM108* | ENSG00000144868.13 | chr3:133038391-133397792 | -0.37 | -1.30 | 6.62 | 19.41 | 1.05E-05 | 1.01E-03 | -2.99 |
| *VWA1* | ENSG00000179403.12 | chr1:1434861-1442882 | -0.50 | -1.42 | 5.38 | 19.37 | 1.08E-05 | 1.04E-03 | -2.98 |
| *PLXDC2* | ENSG00000120594.17 | chr10:19816239-20289856 | -0.33 | -1.25 | 7.52 | 19.36 | 1.08E-05 | 1.04E-03 | -2.98 |
| *SPSB3* | ENSG00000162032.16 | chr16:1776712-1793700 | 0.57 | 1.48 | 4.99 | 19.34 | 1.10E-05 | 1.05E-03 | -2.98 |
| *CPNE2* | ENSG00000140848.17 | chr16:57092583-57148369 | 0.35 | 1.28 | 7.00 | 19.33 | 1.10E-05 | 1.05E-03 | -2.98 |
| *TSHZ3* | ENSG00000121297.8 | chr19:31149979-31349436 | -0.48 | -1.40 | 5.49 | 19.32 | 1.10E-05 | 1.05E-03 | -2.98 |
| *RARB* | ENSG00000077092.19 | chr3:25174332-25597932 | -1.33 | -2.52 | 2.53 | 19.26 | 1.14E-05 | 1.09E-03 | -2.96 |
| *COTL1* | ENSG00000103187.8 | chr16:84565596-84618078 | -0.33 | -1.26 | 7.35 | 19.19 | 1.19E-05 | 1.12E-03 | -2.95 |
| *B3GNT9* | ENSG00000237172.4 | chr16:67148104-67150998 | -0.76 | -1.70 | 4.07 | 19.16 | 1.20E-05 | 1.14E-03 | -2.94 |
| *C3* | ENSG00000125730.17 | chr19:6677704-6730562 | -1.81 | -3.50 | 1.74 | 19.11 | 1.23E-05 | 1.16E-03 | -2.93 |
| *S1PR1* | ENSG00000170989.10 | chr1:101236865-101243713 | 0.52 | 1.44 | 5.24 | 19.11 | 1.23E-05 | 1.16E-03 | -2.93 |
| *CHODL* | ENSG00000154645.14 | chr21:17901263-18267373 | -0.73 | -1.66 | 4.18 | 19.09 | 1.25E-05 | 1.17E-03 | -2.93 |
| *LOXL1* | ENSG00000129038.16 | chr15:73925989-73952137 | 0.51 | 1.42 | 5.33 | 19.06 | 1.26E-05 | 1.19E-03 | -2.93 |
| *PDGFA* | ENSG00000197461.13 | chr7:497258-520296 | 0.96 | 1.94 | 3.37 | 19.04 | 1.28E-05 | 1.20E-03 | -2.92 |
| *LETMD1* | ENSG00000050426.16 | chr12:51047962-51060424 | -0.80 | -1.74 | 3.93 | 19.02 | 1.29E-05 | 1.21E-03 | -2.92 |
| *TMPRSS7* | ENSG00000176040.13 | chr3:112034843-112081269 | -1.09 | -2.12 | 3.02 | 19.02 | 1.30E-05 | 1.21E-03 | -2.92 |
| *CD9* | ENSG00000010278.14 | chr12:6199715-6238271 | -0.48 | -1.39 | 5.49 | 19.01 | 1.30E-05 | 1.21E-03 | -2.92 |
| *ENHO* | ENSG00000168913.7 | chr9:34521043-34522990 | 0.43 | 1.34 | 6.01 | 18.97 | 1.33E-05 | 1.24E-03 | -2.91 |
| *CASKIN1* | ENSG00000167971.16 | chr16:2177180-2196605 | 0.35 | 1.27 | 7.02 | 18.96 | 1.34E-05 | 1.24E-03 | -2.91 |
| *EPHB1* | ENSG00000154928.18 | chr3:134597801-135260467 | 0.44 | 1.36 | 5.81 | 18.95 | 1.34E-05 | 1.24E-03 | -2.91 |
| *CIZ1* | ENSG00000148337.21 | chr9:128161251-128204383 | 0.33 | 1.26 | 7.30 | 18.92 | 1.36E-05 | 1.26E-03 | -2.90 |
| *MLST8* | ENSG00000167965.18 | chr16:2204248-2209453 | 0.40 | 1.32 | 6.27 | 18.91 | 1.37E-05 | 1.26E-03 | -2.90 |
| *HAPLN1* | ENSG00000145681.11 | chr5:83637805-83720855 | 0.36 | 1.28 | 6.81 | 18.90 | 1.38E-05 | 1.27E-03 | -2.90 |
| *FDFT1* | ENSG00000079459.13 | chr8:11795573-11839304 | -0.38 | -1.30 | 6.55 | 18.89 | 1.39E-05 | 1.28E-03 | -2.89 |
| *SRGAP1* | ENSG00000196935.9 | chr12:63844700-64162217 | -0.35 | -1.27 | 6.95 | 18.85 | 1.41E-05 | 1.30E-03 | -2.89 |
| *CASC3* | ENSG00000108349.17 | chr17:40140318-40172171 | 0.33 | 1.26 | 7.31 | 18.80 | 1.46E-05 | 1.33E-03 | -2.87 |
| *P3H3* | ENSG00000110811.20 | chr12:6828407-6839847 | -0.99 | -1.99 | 3.24 | 18.78 | 1.47E-05 | 1.34E-03 | -2.87 |
| *TRIM56* | ENSG00000169871.13 | chr7:101085481-101097967 | -0.45 | -1.37 | 5.71 | 18.78 | 1.47E-05 | 1.34E-03 | -2.87 |
| *GM2A* | ENSG00000196743.9 | chr5:151212150-151270440 | 0.35 | 1.28 | 6.83 | 18.61 | 1.60E-05 | 1.46E-03 | -2.84 |
| *PIP5K1B* | ENSG00000107242.20 | chr9:68705240-69009176 | -1.37 | -2.59 | 2.40 | 18.61 | 1.61E-05 | 1.46E-03 | -2.84 |
| *RFLNB* | ENSG00000183688.4 | chr17:439978-445939 | -0.84 | -1.80 | 3.74 | 18.58 | 1.63E-05 | 1.48E-03 | -2.83 |
| *MARCHF3* | ENSG00000173926.6 | chr5:126867714-127030558 | 0.89 | 1.85 | 3.57 | 18.57 | 1.64E-05 | 1.48E-03 | -2.83 |
| *SAMD9* | ENSG00000205413.8 | chr7:93099513-93118023 | -1.79 | -3.46 | 1.73 | 18.56 | 1.65E-05 | 1.49E-03 | -2.83 |
| *FAM219A* | ENSG00000164970.15 | chr9:34398184-34458570 | 0.33 | 1.26 | 7.27 | 18.45 | 1.74E-05 | 1.57E-03 | -2.80 |
| *ADRA1B* | ENSG00000170214.5 | chr5:159865080-159973012 | 1.14 | 2.21 | 2.82 | 18.37 | 1.82E-05 | 1.64E-03 | -2.79 |
| *ERF* | ENSG00000105722.10 | chr19:42247569-42255128 | 0.39 | 1.31 | 6.28 | 18.33 | 1.86E-05 | 1.67E-03 | -2.78 |
| *SCGN* | ENSG00000079689.14 | chr6:25652201-25701783 | 1.49 | 2.80 | 2.12 | 18.26 | 1.93E-05 | 1.72E-03 | -2.76 |
| *CLCN7* | ENSG00000103249.18 | chr16:1444934-1475084 | 0.35 | 1.27 | 6.82 | 18.25 | 1.94E-05 | 1.73E-03 | -2.76 |
| *PCDH17* | ENSG00000118946.12 | chr13:57631744-57729311 | -0.42 | -1.34 | 5.94 | 18.25 | 1.94E-05 | 1.73E-03 | -2.76 |
| *KRT10* | ENSG00000186395.8 | chr17:40818117-40822614 | 0.53 | 1.45 | 5.10 | 18.22 | 1.97E-05 | 1.76E-03 | -2.76 |
| *TENM2* | ENSG00000145934.16 | chr5:167284799-168264157 | -0.40 | -1.32 | 6.15 | 18.12 | 2.08E-05 | 1.85E-03 | -2.73 |
| *ZNF619* | ENSG00000177873.14 | chr3:40477113-40491053 | -1.41 | -2.66 | 2.27 | 18.08 | 2.12E-05 | 1.88E-03 | -2.73 |
| *SYNPO2* | ENSG00000172403.11 | chr4:118850688-119061247 | -1.17 | -2.24 | 2.78 | 18.04 | 2.17E-05 | 1.91E-03 | -2.72 |
| *GLI2* | ENSG00000074047.22 | chr2:120735623-120992653 | 0.49 | 1.40 | 5.34 | 18.03 | 2.17E-05 | 1.91E-03 | -2.72 |
| *MT3* | ENSG00000087250.9 | chr16:56589074-56591088 | 1.15 | 2.22 | 2.79 | 18.03 | 2.17E-05 | 1.91E-03 | -2.72 |
| *PCDHB16* | ENSG00000272674.3 | chr5:141181399-141186399 | 0.94 | 1.91 | 3.36 | 18.01 | 2.19E-05 | 1.93E-03 | -2.72 |
| *CACNA1C* | ENSG00000151067.22 | chr12:1970786-2697950 | -0.35 | -1.28 | 6.72 | 18.01 | 2.20E-05 | 1.93E-03 | -2.71 |
| *CYB561* | ENSG00000008283.16 | chr17:63432304-63446354 | -0.46 | -1.38 | 5.49 | 17.98 | 2.23E-05 | 1.95E-03 | -2.71 |
| *ZFHX3* | ENSG00000140836.17 | chr16:72782885-73891871 | -0.47 | -1.38 | 5.48 | 17.97 | 2.25E-05 | 1.97E-03 | -2.71 |
| *ARRDC4* | ENSG00000140450.9 | chr15:97960703-97973833 | -0.63 | -1.55 | 4.56 | 17.96 | 2.26E-05 | 1.97E-03 | -2.70 |
| *SORCS1* | ENSG00000108018.15 | chr10:106573663-107164534 | -0.60 | -1.51 | 4.73 | 17.93 | 2.29E-05 | 1.99E-03 | -2.70 |
| *GRB10* | ENSG00000106070.20 | chr7:50590063-50793462 | 0.45 | 1.36 | 5.64 | 17.93 | 2.29E-05 | 1.99E-03 | -2.70 |
| *CCDC167* | ENSG00000198937.9 | chr6:37482938-37499893 | 0.58 | 1.49 | 4.82 | 17.79 | 2.46E-05 | 2.13E-03 | -2.67 |
| *ELOVL2* | ENSG00000197977.4 | chr6:10980759-11044305 | 0.37 | 1.29 | 6.45 | 17.78 | 2.49E-05 | 2.14E-03 | -2.67 |
| *KIRREL1* | ENSG00000183853.18 | chr1:157993273-158100262 | -0.39 | -1.31 | 6.22 | 17.77 | 2.49E-05 | 2.14E-03 | -2.67 |
| *HEXIM1* | ENSG00000186834.4 | chr17:45148475-45152099 | -0.45 | -1.36 | 5.62 | 17.75 | 2.52E-05 | 2.17E-03 | -2.66 |
| *NCK2* | ENSG00000071051.14 | chr2:105744912-105894274 | -0.33 | -1.25 | 7.24 | 17.69 | 2.60E-05 | 2.23E-03 | -2.65 |
| *RNF19B* | ENSG00000116514.16 | chr1:32936445-32964685 | 0.47 | 1.39 | 5.43 | 17.68 | 2.62E-05 | 2.24E-03 | -2.65 |
| *TFPI* | ENSG00000003436.16 | chr2:187464230-187565760 | 0.48 | 1.40 | 5.36 | 17.65 | 2.65E-05 | 2.27E-03 | -2.64 |
| *C6orf136* | ENSG00000204564.12 | chr6:30647039-30653210 | 0.55 | 1.46 | 4.97 | 17.65 | 2.65E-05 | 2.27E-03 | -2.64 |
| *ZMAT4* | ENSG00000165061.15 | chr8:40530590-40897833 | -0.72 | -1.64 | 4.12 | 17.49 | 2.88E-05 | 2.46E-03 | -2.61 |
| *OPRM1* | ENSG00000112038.18 | chr6:154010496-154246867 | -1.29 | -2.44 | 2.48 | 17.38 | 3.07E-05 | 2.59E-03 | -2.59 |
| *EPHB3* | ENSG00000182580.3 | chr3:184561785-184582408 | 0.98 | 1.97 | 3.16 | 17.36 | 3.10E-05 | 2.61E-03 | -2.58 |
| *P2RX3* | ENSG00000109991.9 | chr11:57338352-57372396 | -0.93 | -1.91 | 3.32 | 17.34 | 3.12E-05 | 2.63E-03 | -2.58 |
| *SIX3* | ENSG00000138083.5 | chr2:44941702-44946071 | -0.89 | -1.85 | 3.47 | 17.31 | 3.18E-05 | 2.68E-03 | -2.57 |
| *GASK1B* | ENSG00000164125.15 | chr4:158124474-158173318 | 0.39 | 1.31 | 6.15 | 17.22 | 3.34E-05 | 2.80E-03 | -2.55 |
| *DTX1* | ENSG00000135144.7 | chr12:113056709-113098028 | 0.39 | 1.31 | 6.15 | 17.17 | 3.42E-05 | 2.86E-03 | -2.54 |
| *NUDT14* | ENSG00000183828.15 | chr14:105172938-105181323 | 0.71 | 1.63 | 4.10 | 17.10 | 3.55E-05 | 2.96E-03 | -2.53 |
| *C9orf16* | ENSG00000171159.5 | chr9:128160265-128163924 | 0.35 | 1.27 | 6.68 | 17.08 | 3.58E-05 | 2.98E-03 | -2.53 |
| *TOP2A* | ENSG00000131747.15 | chr17:40388525-40417896 | -0.55 | -1.46 | 4.92 | 17.07 | 3.61E-05 | 3.01E-03 | -2.52 |
| *HERC1* | ENSG00000103657.14 | chr15:63608618-63833948 | -0.33 | -1.26 | 6.99 | 17.06 | 3.62E-05 | 3.01E-03 | -2.52 |
| *SLC20A2* | ENSG00000168575.10 | chr8:42416475-42541926 | 0.38 | 1.30 | 6.30 | 17.02 | 3.70E-05 | 3.07E-03 | -2.51 |
| *RPUSD1* | ENSG00000007376.8 | chr16:784974-788397 | 0.77 | 1.71 | 3.85 | 17.02 | 3.71E-05 | 3.07E-03 | -2.51 |
| *RPE65* | ENSG00000116745.7 | chr1:68428822-68449954 | -0.44 | -1.35 | 5.64 | 16.90 | 3.94E-05 | 3.25E-03 | -2.49 |
| *MEX3A* | ENSG00000254726.3 | chr1:156072013-156082465 | 0.34 | 1.26 | 6.86 | 16.88 | 3.98E-05 | 3.28E-03 | -2.48 |
| *NID2* | ENSG00000087303.18 | chr14:52004803-52069228 | -1.86 | -3.64 | 1.53 | 16.83 | 4.09E-05 | 3.36E-03 | -2.47 |
| *GCN1* | ENSG00000089154.11 | chr12:120127202-120194715 | -0.39 | -1.31 | 6.17 | 16.77 | 4.22E-05 | 3.46E-03 | -2.46 |
| *PEAK1* | ENSG00000173517.10 | chr15:77100656-77420144 | -0.44 | -1.35 | 5.62 | 16.72 | 4.34E-05 | 3.56E-03 | -2.45 |
| *BMF* | ENSG00000104081.14 | chr15:40087890-40108892 | 1.16 | 2.24 | 2.69 | 16.72 | 4.35E-05 | 3.56E-03 | -2.45 |
| *DYNC2H1* | ENSG00000187240.16 | chr11:103109410-103479863 | -0.49 | -1.40 | 5.24 | 16.71 | 4.35E-05 | 3.56E-03 | -2.45 |
| *TOMM7* | ENSG00000196683.10 | chr7:22812628-22822852 | 0.40 | 1.32 | 5.94 | 16.68 | 4.43E-05 | 3.61E-03 | -2.44 |
| *RUNX3* | ENSG00000020633.19 | chr1:24899511-24965121 | 8.37 | 330.95 | -0.95 | 16.65 | 4.50E-05 | 3.66E-03 | -2.44 |
| *PIEZO1* | ENSG00000103335.22 | chr16:88715338-88785220 | -0.49 | -1.40 | 5.23 | 16.63 | 4.54E-05 | 3.68E-03 | -2.43 |
| *PLEKHG3* | ENSG00000126822.17 | chr14:64704102-64750249 | -0.45 | -1.36 | 5.50 | 16.57 | 4.70E-05 | 3.81E-03 | -2.42 |
| *TFAP2C* | ENSG00000087510.7 | chr20:56629306-56639283 | -1.50 | -2.83 | 1.97 | 16.53 | 4.78E-05 | 3.86E-03 | -2.41 |
| *ASS1* | ENSG00000130707.18 | chr9:130444961-130501274 | -0.46 | -1.38 | 5.39 | 16.47 | 4.94E-05 | 3.98E-03 | -2.40 |
| *HERC2* | ENSG00000128731.18 | chr15:28111040-28322179 | -0.32 | -1.25 | 7.04 | 16.47 | 4.96E-05 | 3.99E-03 | -2.40 |
| *SLC4A4* | ENSG00000080493.17 | chr4:71062667-71572087 | 0.33 | 1.26 | 6.85 | 16.41 | 5.11E-05 | 4.10E-03 | -2.39 |
| *CNFN* | ENSG00000105427.10 | chr19:42387019-42390297 | -0.95 | -1.93 | 3.17 | 16.37 | 5.23E-05 | 4.17E-03 | -2.38 |
| *APOLD1* | ENSG00000178878.13 | chr12:12725917-12829975 | -1.21 | -2.32 | 2.56 | 16.33 | 5.32E-05 | 4.24E-03 | -2.37 |
| *STEAP3* | ENSG00000115107.20 | chr2:119223831-119265652 | -0.50 | -1.42 | 5.12 | 16.30 | 5.42E-05 | 4.31E-03 | -2.37 |
| *RPL23A* | ENSG00000198242.14 | chr17:28719985-28724359 | 0.33 | 1.25 | 6.92 | 16.28 | 5.47E-05 | 4.35E-03 | -2.36 |
| *APCDD1* | ENSG00000154856.13 | chr18:10454635-10489949 | 0.35 | 1.28 | 6.47 | 16.23 | 5.61E-05 | 4.44E-03 | -2.35 |
| *CDCP1* | ENSG00000163814.8 | chr3:45082277-45146422 | -0.95 | -1.93 | 3.16 | 16.21 | 5.67E-05 | 4.48E-03 | -2.35 |
| *SCAF11* | ENSG00000139218.18 | chr12:45919131-45992120 | -0.39 | -1.31 | 6.02 | 16.21 | 5.67E-05 | 4.48E-03 | -2.35 |
| *UBTD1* | ENSG00000165886.5 | chr10:97498924-97571206 | 0.40 | 1.32 | 5.94 | 16.21 | 5.68E-05 | 4.48E-03 | -2.35 |
| *SLC38A3* | ENSG00000188338.15 | chr3:50205246-50221486 | 0.41 | 1.33 | 5.86 | 16.15 | 5.84E-05 | 4.60E-03 | -2.34 |
| *SLC4A3* | ENSG00000114923.17 | chr2:219627394-219641980 | 0.32 | 1.25 | 6.96 | 16.13 | 5.91E-05 | 4.64E-03 | -2.33 |
| *REST* | ENSG00000084093.19 | chr4:56907876-56966808 | -0.40 | -1.32 | 5.98 | 16.09 | 6.06E-05 | 4.75E-03 | -2.32 |
| *MAMLD1* | ENSG00000013619.14 | chrX:150361422-150514178 | 0.37 | 1.29 | 6.30 | 16.06 | 6.13E-05 | 4.80E-03 | -2.32 |
| *IPO8* | ENSG00000133704.10 | chr12:30628988-30695869 | -0.50 | -1.42 | 5.10 | 16.06 | 6.14E-05 | 4.80E-03 | -2.32 |
| *PGAP1* | ENSG00000197121.15 | chr2:196833004-196927796 | -0.34 | -1.27 | 6.57 | 16.01 | 6.30E-05 | 4.90E-03 | -2.31 |
| *MAPK15* | ENSG00000181085.15 | chr8:143716340-143722458 | -0.87 | -1.83 | 3.40 | 15.99 | 6.37E-05 | 4.96E-03 | -2.30 |
| *MGA* | ENSG00000174197.16 | chr15:41621224-41773081 | -0.37 | -1.29 | 6.23 | 15.94 | 6.54E-05 | 5.07E-03 | -2.29 |
| *FILIP1L* | ENSG00000168386.18 | chr3:99830141-100114513 | -0.65 | -1.57 | 4.26 | 15.92 | 6.61E-05 | 5.11E-03 | -2.29 |
| *TBKBP1* | ENSG00000198933.9 | chr17:47694081-47712050 | 0.57 | 1.48 | 4.67 | 15.86 | 6.82E-05 | 5.25E-03 | -2.28 |
| *FAM167A* | ENSG00000154319.16 | chr8:11421476-11475908 | -0.47 | -1.39 | 5.26 | 15.85 | 6.88E-05 | 5.29E-03 | -2.28 |
| *MID1IP1* | ENSG00000165175.15 | chrX:38801432-38806537 | 0.43 | 1.35 | 5.52 | 15.81 | 7.00E-05 | 5.37E-03 | -2.27 |
| *ECM2* | ENSG00000106823.12 | chr9:92493554-92536655 | -0.93 | -1.90 | 3.17 | 15.80 | 7.05E-05 | 5.40E-03 | -2.27 |
| *BBC3* | ENSG00000105327.18 | chr19:47220822-47232766 | 0.69 | 1.62 | 4.05 | 15.80 | 7.05E-05 | 5.40E-03 | -2.27 |
| *ABI3BP* | ENSG00000154175.18 | chr3:100749156-100993515 | -0.78 | -1.72 | 3.72 | 15.79 | 7.08E-05 | 5.41E-03 | -2.27 |
| *PANX2* | ENSG00000073150.14 | chr22:50170731-50180295 | 0.58 | 1.50 | 4.57 | 15.74 | 7.27E-05 | 5.54E-03 | -2.26 |
| *PLAT* | ENSG00000104368.18 | chr8:42174718-42207676 | -0.34 | -1.27 | 6.52 | 15.74 | 7.29E-05 | 5.55E-03 | -2.26 |
| *TAFA5* | ENSG00000219438.9 | chr22:48489553-48850912 | 0.44 | 1.36 | 5.44 | 15.71 | 7.40E-05 | 5.63E-03 | -2.25 |
| *SOX21* | ENSG00000125285.6 | chr13:94709622-94712545 | 0.35 | 1.28 | 6.41 | 15.68 | 7.50E-05 | 5.68E-03 | -2.25 |
| *PLXDC1* | ENSG00000161381.14 | chr17:39063313-39154394 | -0.79 | -1.73 | 3.69 | 15.68 | 7.51E-05 | 5.68E-03 | -2.25 |
| *PDE1C* | ENSG00000154678.18 | chr7:31751179-32428131 | -0.58 | -1.49 | 4.60 | 15.60 | 7.81E-05 | 5.88E-03 | -2.23 |
| *GDAP1L1* | ENSG00000124194.17 | chr20:44247099-44280947 | 0.48 | 1.39 | 5.21 | 15.60 | 7.85E-05 | 5.90E-03 | -2.23 |
| *NR1D1* | ENSG00000126368.6 | chr17:40092793-40100589 | 0.43 | 1.35 | 5.48 | 15.58 | 7.91E-05 | 5.93E-03 | -2.23 |
| *NKD2* | ENSG00000145506.14 | chr5:1008802-1038943 | -0.45 | -1.37 | 5.35 | 15.57 | 7.95E-05 | 5.95E-03 | -2.23 |
| *FAM107A* | ENSG00000168309.18 | chr3:58564117-58627610 | 0.60 | 1.51 | 4.48 | 15.57 | 7.95E-05 | 5.95E-03 | -2.23 |
| *KCNMA1* | ENSG00000156113.24 | chr10:76869601-77638369 | -0.38 | -1.30 | 6.08 | 15.54 | 8.07E-05 | 6.02E-03 | -2.22 |
| *TNFRSF11B* | ENSG00000164761.9 | chr8:118923557-118951885 | -0.73 | -1.66 | 3.88 | 15.54 | 8.09E-05 | 6.02E-03 | -2.22 |
| *SLC47A2* | ENSG00000180638.18 | chr17:19678288-19718979 | 0.58 | 1.49 | 4.60 | 15.54 | 8.10E-05 | 6.03E-03 | -2.22 |
| *RAB5C* | ENSG00000108774.15 | chr17:42124978-42155044 | 0.39 | 1.31 | 5.99 | 15.53 | 8.14E-05 | 6.05E-03 | -2.22 |
| *RAC3* | ENSG00000169750.9 | chr17:82031678-82034204 | 0.40 | 1.32 | 5.83 | 15.44 | 8.53E-05 | 6.32E-03 | -2.20 |
| *ZNF385D* | ENSG00000151789.12 | chr3:21412218-22373321 | 0.43 | 1.35 | 5.51 | 15.42 | 8.61E-05 | 6.37E-03 | -2.20 |
| *CLVS2* | ENSG00000146352.13 | chr6:122996235-123072925 | 0.48 | 1.39 | 5.19 | 15.39 | 8.75E-05 | 6.46E-03 | -2.19 |
| *NIBAN1* | ENSG00000135842.17 | chr1:184790724-184974508 | -1.38 | -2.59 | 2.09 | 15.38 | 8.80E-05 | 6.48E-03 | -2.19 |
| *NUMBL* | ENSG00000105245.9 | chr19:40665905-40690972 | 0.42 | 1.34 | 5.61 | 15.37 | 8.82E-05 | 6.49E-03 | -2.19 |
| *NUDT4* | ENSG00000173598.14 | chr12:93377883-93408146 | 0.36 | 1.28 | 6.30 | 15.37 | 8.84E-05 | 6.49E-03 | -2.19 |
| *MED29* | ENSG00000063322.15 | chr19:39391303-39400641 | 0.36 | 1.28 | 6.29 | 15.36 | 8.88E-05 | 6.50E-03 | -2.19 |
| *CEND1* | ENSG00000184524.6 | chr11:787115-790113 | 0.48 | 1.40 | 5.16 | 15.33 | 9.02E-05 | 6.58E-03 | -2.18 |
| *MXI1* | ENSG00000119950.21 | chr10:110207605-110287365 | 0.37 | 1.29 | 6.17 | 15.30 | 9.17E-05 | 6.67E-03 | -2.18 |
| *SERPINB1* | ENSG00000021355.13 | chr6:2832332-2841959 | -2.56 | -5.92 | 0.72 | 15.27 | 9.31E-05 | 6.76E-03 | -2.17 |
| *LMNA* | ENSG00000160789.21 | chr1:156082573-156140089 | -0.39 | -1.31 | 5.91 | 15.20 | 9.66E-05 | 7.00E-03 | -2.15 |
| *ARHGAP42* | ENSG00000165895.19 | chr11:100687288-100993941 | 0.41 | 1.33 | 5.66 | 15.18 | 9.79E-05 | 7.08E-03 | -2.15 |
| *PCOLCE* | ENSG00000106333.13 | chr7:100602363-100608175 | -0.64 | -1.56 | 4.22 | 15.16 | 9.86E-05 | 7.11E-03 | -2.15 |
| *PCDH18* | ENSG00000189184.11 | chr4:137518918-137532494 | 0.61 | 1.53 | 4.34 | 15.17 | 9.85E-05 | 7.11E-03 | -2.15 |
| *UBAC1* | ENSG00000130560.9 | chr9:135932969-135961373 | 0.57 | 1.49 | 4.58 | 15.16 | 9.89E-05 | 7.12E-03 | -2.15 |
| *BAG3* | ENSG00000151929.10 | chr10:119651380-119677819 | 0.32 | 1.25 | 6.69 | 15.14 | 1.00E-04 | 7.19E-03 | -2.14 |
| *H1-0* | ENSG00000189060.5 | chr22:37805093-37807436 | -0.34 | -1.26 | 6.52 | 15.13 | 1.00E-04 | 7.21E-03 | -2.14 |
| *KSR1* | ENSG00000141068.14 | chr17:27456470-27626438 | 0.36 | 1.29 | 6.19 | 15.11 | 1.02E-04 | 7.29E-03 | -2.14 |
| *TNFAIP8L1* | ENSG00000185361.9 | chr19:4639516-4655568 | -0.94 | -1.92 | 3.09 | 15.09 | 1.02E-04 | 7.33E-03 | -2.14 |
| *SMOC2* | ENSG00000112562.19 | chr6:168441151-168673445 | -1.22 | -2.34 | 2.41 | 15.08 | 1.03E-04 | 7.36E-03 | -2.13 |
| *IFITM10* | ENSG00000244242.2 | chr11:1732406-1750595 | 0.97 | 1.96 | 2.98 | 15.08 | 1.03E-04 | 7.36E-03 | -2.13 |
| *TNS3* | ENSG00000136205.17 | chr7:47275154-47582558 | -0.42 | -1.34 | 5.53 | 15.05 | 1.05E-04 | 7.47E-03 | -2.13 |
| *FAM181B* | ENSG00000182103.5 | chr11:82729940-82733864 | 0.57 | 1.48 | 4.60 | 15.04 | 1.06E-04 | 7.51E-03 | -2.12 |
| *CHMP1B* | ENSG00000255112.3 | chr18:11851413-11854444 | -0.39 | -1.31 | 5.89 | 15.01 | 1.07E-04 | 7.58E-03 | -2.12 |
| *NR4A3* | ENSG00000119508.18 | chr9:99821855-99866891 | -0.38 | -1.30 | 5.98 | 15.01 | 1.07E-04 | 7.58E-03 | -2.12 |
| *CARMIL1* | ENSG00000079691.18 | chr6:25279078-25620530 | 0.42 | 1.33 | 5.56 | 15.00 | 1.07E-04 | 7.61E-03 | -2.12 |
| *RALYL* | ENSG00000184672.12 | chr8:84182787-84921844 | 0.66 | 1.58 | 4.10 | 14.95 | 1.10E-04 | 7.79E-03 | -2.11 |
| *PLA2R1* | ENSG00000153246.13 | chr2:159932006-160062615 | -0.96 | -1.94 | 3.02 | 14.92 | 1.12E-04 | 7.92E-03 | -2.10 |
| *TNFRSF12A* | ENSG00000006327.14 | chr16:3018445-3022383 | -0.39 | -1.31 | 5.85 | 14.90 | 1.14E-04 | 8.00E-03 | -2.10 |
| *KAT5* | ENSG00000172977.13 | chr11:65711996-65719604 | 0.44 | 1.36 | 5.37 | 14.89 | 1.14E-04 | 8.00E-03 | -2.10 |
| *TSHZ1* | ENSG00000179981.11 | chr18:75210755-75289950 | 0.44 | 1.36 | 5.35 | 14.89 | 1.14E-04 | 8.00E-03 | -2.10 |
| *TNFAIP3* | ENSG00000118503.15 | chr6:137867214-137883312 | -0.72 | -1.64 | 3.89 | 14.83 | 1.18E-04 | 8.25E-03 | -2.08 |
| *ART3* | ENSG00000156219.17 | chr4:76011184-76112802 | -1.25 | -2.38 | 2.32 | 14.82 | 1.18E-04 | 8.28E-03 | -2.08 |
| *SEMA3E* | ENSG00000170381.14 | chr7:83363238-83649139 | 2.30 | 4.92 | 0.84 | 14.81 | 1.19E-04 | 8.29E-03 | -2.08 |
| *PRG4* | ENSG00000116690.13 | chr1:186296279-186314567 | -1.97 | -3.93 | 1.25 | 14.78 | 1.21E-04 | 8.41E-03 | -2.08 |
| *RPRM* | ENSG00000177519.4 | chr2:153477338-153478762 | 0.83 | 1.78 | 3.40 | 14.78 | 1.21E-04 | 8.42E-03 | -2.07 |
| *GPANK1* | ENSG00000204438.11 | chr6:31661228-31666283 | 0.92 | 1.90 | 3.09 | 14.74 | 1.24E-04 | 8.57E-03 | -2.07 |
| *EN1* | ENSG00000163064.7 | chr2:118842171-118847648 | 1.09 | 2.13 | 2.68 | 14.74 | 1.23E-04 | 8.57E-03 | -2.07 |
| *TET1* | ENSG00000138336.9 | chr10:68560337-68694487 | -0.38 | -1.30 | 5.97 | 14.73 | 1.24E-04 | 8.62E-03 | -2.06 |
| *FZD7* | ENSG00000155760.2 | chr2:202034587-202038445 | -0.51 | -1.42 | 4.94 | 14.72 | 1.25E-04 | 8.62E-03 | -2.06 |
| *BRPF3* | ENSG00000096070.19 | chr6:36196744-36232790 | 0.33 | 1.26 | 6.51 | 14.70 | 1.26E-04 | 8.71E-03 | -2.06 |
| *MT1F* | ENSG00000198417.7 | chr16:56657731-56660698 | 0.71 | 1.64 | 3.87 | 14.70 | 1.26E-04 | 8.71E-03 | -2.06 |
| *RBP4* | ENSG00000138207.14 | chr10:93591687-93601744 | -1.04 | -2.05 | 2.81 | 14.67 | 1.28E-04 | 8.82E-03 | -2.05 |
| *GRK2* | ENSG00000173020.11 | chr11:67266473-67286556 | 0.36 | 1.28 | 6.20 | 14.63 | 1.31E-04 | 9.01E-03 | -2.05 |
| *GAREM1* | ENSG00000141441.16 | chr18:32124877-32470882 | -0.39 | -1.31 | 5.84 | 14.61 | 1.32E-04 | 9.08E-03 | -2.04 |
| *RDH10* | ENSG00000121039.10 | chr8:73294602-73325281 | -0.86 | -1.81 | 3.30 | 14.56 | 1.36E-04 | 9.32E-03 | -2.03 |
| *LSM14B* | ENSG00000149657.20 | chr20:62122461-62135374 | 0.38 | 1.30 | 5.94 | 14.51 | 1.39E-04 | 9.54E-03 | -2.02 |
| *ABCA13* | ENSG00000179869.15 | chr7:48171458-48647497 | -1.42 | -2.68 | 1.94 | 14.50 | 1.40E-04 | 9.56E-03 | -2.02 |
| *GPAT3* | ENSG00000138678.11 | chr4:83535914-83605875 | 8.22 | 298.45 | -1.08 | 14.50 | 1.40E-04 | 9.56E-03 | -2.02 |
| *SH3BP5* | ENSG00000131370.16 | chr3:15254353-15341368 | -0.36 | -1.29 | 6.11 | 14.50 | 1.40E-04 | 9.56E-03 | -2.02 |
| *PDK3* | ENSG00000067992.16 | chrX:24465270-24550466 | -0.38 | -1.30 | 5.92 | 14.49 | 1.41E-04 | 9.60E-03 | -2.02 |
| *RAMP1* | ENSG00000132329.11 | chr2:237858893-237912106 | -0.49 | -1.41 | 5.01 | 14.48 | 1.42E-04 | 9.66E-03 | -2.01 |
| *GPR50* | ENSG00000102195.10 | chrX:151176584-151181465 | 0.79 | 1.73 | 3.53 | 14.47 | 1.42E-04 | 9.68E-03 | -2.01 |
| *ZNF667* | ENSG00000198046.12 | chr19:56439325-56478065 | -0.94 | -1.92 | 3.03 | 14.47 | 1.43E-04 | 9.70E-03 | -2.01 |
| *PROS1* | ENSG00000184500.16 | chr3:93873051-93980003 | -0.49 | -1.40 | 5.04 | 14.46 | 1.43E-04 | 9.72E-03 | -2.01 |
| *GPR176* | ENSG00000166073.11 | chr15:39799008-39920266 | -0.41 | -1.32 | 5.62 | 14.45 | 1.44E-04 | 9.75E-03 | -2.01 |
| *CSRP2* | ENSG00000175183.10 | chr12:76858709-76879023 | -0.43 | -1.35 | 5.41 | 14.42 | 1.47E-04 | 9.91E-03 | -2.00 |
| *TP53I3* | ENSG00000115129.14 | chr2:24077433-24085861 | -0.68 | -1.60 | 4.00 | 14.41 | 1.47E-04 | 9.91E-03 | -2.00 |
| *HELLS* | ENSG00000119969.15 | chr10:94501434-94613905 | -0.94 | -1.92 | 3.03 | 14.40 | 1.48E-04 | 9.94E-03 | -2.00 |
| *C3orf70* | ENSG00000187068.3 | chr3:185076838-185153060 | 0.38 | 1.30 | 5.92 | 14.39 | 1.48E-04 | 9.94E-03 | -2.00 |
| *PLCXD3* | ENSG00000182836.10 | chr5:41306952-41510628 | 0.59 | 1.51 | 4.36 | 14.39 | 1.48E-04 | 9.94E-03 | -2.00 |
| *SKIDA1* | ENSG00000180592.17 | chr10:21513475-21526368 | -0.50 | -1.41 | 4.98 | 14.35 | 1.52E-04 | 1.01E-02 | -1.99 |
| *PNCK* | ENSG00000130822.16 | chrX:153669733-153689010 | 0.52 | 1.44 | 4.78 | 14.33 | 1.53E-04 | 1.02E-02 | -1.99 |
| *MAL* | ENSG00000172005.11 | chr2:95025677-95053992 | -0.61 | -1.53 | 4.26 | 14.33 | 1.54E-04 | 1.03E-02 | -1.99 |
| *BTBD17* | ENSG00000204347.4 | chr17:74356416-74361868 | 0.88 | 1.84 | 3.17 | 14.31 | 1.55E-04 | 1.03E-02 | -1.99 |
| *HES4* | ENSG00000188290.11 | chr1:998962-1000172 | 0.41 | 1.33 | 5.51 | 14.27 | 1.59E-04 | 1.05E-02 | -1.98 |
| *PATL1* | ENSG00000166889.14 | chr11:59636716-59669037 | 0.36 | 1.28 | 6.13 | 14.25 | 1.60E-04 | 1.06E-02 | -1.97 |
| *INTU* | ENSG00000164066.13 | chr4:127623271-127726737 | -0.57 | -1.49 | 4.47 | 14.21 | 1.64E-04 | 1.08E-02 | -1.97 |
| *TINCR* | ENSG00000223573.7 | chr19:5558167-5578349 | 1.04 | 2.06 | 2.74 | 14.18 | 1.66E-04 | 1.10E-02 | -1.96 |
| *NAPA* | ENSG00000105402.8 | chr19:47487637-47515091 | 0.37 | 1.29 | 6.02 | 14.13 | 1.71E-04 | 1.12E-02 | -1.95 |
| *H2AC21* | ENSG00000184270.6 | chr1:149887469-149887965 | 0.65 | 1.57 | 4.07 | 14.10 | 1.73E-04 | 1.14E-02 | -1.94 |
| *H2AW* | ENSG00000181218.5 | chr1:228456979-228457873 | -0.68 | -1.60 | 3.97 | 14.10 | 1.74E-04 | 1.14E-02 | -1.94 |
| *POU4F1* | ENSG00000152192.8 | chr13:78598362-78603552 | 0.74 | 1.67 | 3.71 | 14.06 | 1.77E-04 | 1.16E-02 | -1.94 |
| *EPPK1* | ENSG00000261150.3 | chr8:143857324-143878467 | -0.97 | -1.96 | 2.93 | 14.05 | 1.78E-04 | 1.16E-02 | -1.94 |
| *LTBP4* | ENSG00000090006.18 | chr19:40592883-40629818 | -0.34 | -1.27 | 6.34 | 14.03 | 1.80E-04 | 1.17E-02 | -1.93 |
| *SFMBT2* | ENSG00000198879.12 | chr10:7158624-7411486 | -0.56 | -1.47 | 4.53 | 14.00 | 1.83E-04 | 1.19E-02 | -1.93 |
| *SLC25A22* | ENSG00000177542.11 | chr11:790475-798281 | 0.40 | 1.32 | 5.61 | 13.99 | 1.84E-04 | 1.19E-02 | -1.92 |
| *THBS4* | ENSG00000113296.14 | chr5:79991311-80083287 | 0.79 | 1.72 | 3.50 | 13.94 | 1.89E-04 | 1.23E-02 | -1.91 |
| *SHC1* | ENSG00000160691.19 | chr1:154962298-154974395 | -0.37 | -1.29 | 5.94 | 13.91 | 1.92E-04 | 1.24E-02 | -1.91 |
| *MPC1* | ENSG00000060762.19 | chr6:166364919-166383013 | 0.45 | 1.36 | 5.24 | 13.89 | 1.94E-04 | 1.25E-02 | -1.90 |
| *FOXC1* | ENSG00000054598.9 | chr6:1609915-1613897 | -0.85 | -1.80 | 3.27 | 13.88 | 1.95E-04 | 1.25E-02 | -1.90 |
| *AJUBA* | ENSG00000129474.16 | chr14:22971177-22982551 | -0.52 | -1.44 | 4.75 | 13.87 | 1.96E-04 | 1.26E-02 | -1.90 |
| *DSC2* | ENSG00000134755.17 | chr18:31058840-31102522 | -1.28 | -2.42 | 2.16 | 13.84 | 1.99E-04 | 1.28E-02 | -1.89 |
| *OCIAD2* | ENSG00000145247.12 | chr4:48885019-48906937 | -1.22 | -2.33 | 2.29 | 13.81 | 2.02E-04 | 1.29E-02 | -1.89 |
| *LRRC8C* | ENSG00000171488.15 | chr1:89633072-89769903 | -0.82 | -1.77 | 3.35 | 13.81 | 2.03E-04 | 1.29E-02 | -1.89 |
| *RNF141* | ENSG00000110315.7 | chr11:10511673-10541230 | 0.36 | 1.28 | 6.09 | 13.81 | 2.02E-04 | 1.29E-02 | -1.89 |
| *IGSF21* | ENSG00000117154.12 | chr1:18107798-18378483 | 0.62 | 1.53 | 4.20 | 13.81 | 2.03E-04 | 1.29E-02 | -1.89 |
| *HEY2* | ENSG00000135547.9 | chr6:125747664-125761269 | 0.48 | 1.39 | 5.00 | 13.79 | 2.05E-04 | 1.30E-02 | -1.88 |
| *NEUROD2* | ENSG00000171532.5 | chr17:39603536-39609777 | -0.65 | -1.57 | 4.07 | 13.77 | 2.07E-04 | 1.32E-02 | -1.88 |
| *ELN* | ENSG00000049540.17 | chr7:74027789-74069907 | 0.36 | 1.28 | 6.07 | 13.75 | 2.09E-04 | 1.33E-02 | -1.88 |
| *DCHS2* | ENSG00000197410.14 | chr4:154231742-154491799 | -1.05 | -2.07 | 2.69 | 13.71 | 2.13E-04 | 1.35E-02 | -1.87 |
| *KLHL42* | ENSG00000087448.11 | chr12:27780048-27803040 | -0.36 | -1.28 | 6.09 | 13.70 | 2.14E-04 | 1.35E-02 | -1.87 |
| *PPP1R17* | ENSG00000106341.11 | chr7:31687215-31708455 | -1.12 | -2.18 | 2.51 | 13.65 | 2.21E-04 | 1.39E-02 | -1.86 |
| *PTPRU* | ENSG00000060656.20 | chr1:29236516-29326813 | -0.38 | -1.30 | 5.79 | 13.65 | 2.21E-04 | 1.39E-02 | -1.86 |
| *WNT4* | ENSG00000162552.15 | chr1:22117313-22143969 | 0.86 | 1.82 | 3.15 | 13.62 | 2.24E-04 | 1.41E-02 | -1.85 |
| *FKBP11* | ENSG00000134285.11 | chr12:48921518-48926474 | -0.94 | -1.92 | 2.95 | 13.61 | 2.26E-04 | 1.42E-02 | -1.85 |
| *FP565260.3* | ENSG00000277117.5 | chr21:5022493-5040666 | 1.64 | 3.11 | 1.54 | 13.60 | 2.26E-04 | 1.42E-02 | -1.85 |
| *FGFR3* | ENSG00000068078.19 | chr4:1793293-1808872 | 0.54 | 1.45 | 4.61 | 13.59 | 2.28E-04 | 1.43E-02 | -1.85 |
| *ADCY9* | ENSG00000162104.10 | chr16:3953387-4116442 | -0.35 | -1.27 | 6.15 | 13.58 | 2.29E-04 | 1.43E-02 | -1.84 |
| *SLC1A6* | ENSG00000105143.12 | chr19:14950034-15022990 | 0.66 | 1.58 | 3.98 | 13.58 | 2.29E-04 | 1.43E-02 | -1.84 |
| *SLC66A1* | ENSG00000040487.13 | chr1:19312326-19329300 | 0.57 | 1.48 | 4.39 | 13.54 | 2.34E-04 | 1.46E-02 | -1.84 |
| *WDR74* | ENSG00000133316.15 | chr11:62832342-62841809 | 0.73 | 1.65 | 3.69 | 13.53 | 2.34E-04 | 1.46E-02 | -1.84 |
| *SUSD6* | ENSG00000100647.8 | chr14:69611596-69715144 | -0.38 | -1.30 | 5.78 | 13.53 | 2.35E-04 | 1.46E-02 | -1.83 |
| *FILIP1* | ENSG00000118407.15 | chr6:75291859-75493800 | -0.88 | -1.84 | 3.11 | 13.52 | 2.36E-04 | 1.46E-02 | -1.83 |
| *GTF3C4* | ENSG00000125484.12 | chr9:132670035-132694953 | -0.35 | -1.28 | 6.08 | 13.52 | 2.36E-04 | 1.46E-02 | -1.83 |
| *TANC1* | ENSG00000115183.15 | chr2:158968640-159232659 | -0.35 | -1.27 | 6.16 | 13.52 | 2.36E-04 | 1.46E-02 | -1.83 |
| *G3BP1* | ENSG00000145907.15 | chr5:151771045-151812785 | -0.35 | -1.27 | 6.18 | 13.50 | 2.39E-04 | 1.48E-02 | -1.83 |
| *CTTNBP2* | ENSG00000077063.11 | chr7:117710651-117874139 | -0.90 | -1.86 | 3.07 | 13.48 | 2.41E-04 | 1.49E-02 | -1.83 |
| *ZC3H3* | ENSG00000014164.7 | chr8:143437659-143541447 | 0.45 | 1.37 | 5.13 | 13.47 | 2.42E-04 | 1.49E-02 | -1.83 |
| *CERS5* | ENSG00000139624.14 | chr12:50129289-50167533 | -0.41 | -1.33 | 5.41 | 13.46 | 2.43E-04 | 1.50E-02 | -1.82 |
| *LGMN* | ENSG00000100600.15 | chr14:92703807-92748679 | -0.38 | -1.30 | 5.70 | 13.42 | 2.49E-04 | 1.53E-02 | -1.82 |
| *IL11RA* | ENSG00000137070.18 | chr9:34650702-34661902 | -0.87 | -1.83 | 3.12 | 13.42 | 2.49E-04 | 1.53E-02 | -1.82 |
| *DIAPH1* | ENSG00000131504.17 | chr5:141515016-141619055 | -0.43 | -1.34 | 5.32 | 13.42 | 2.49E-04 | 1.53E-02 | -1.82 |
| *MSX2* | ENSG00000120149.9 | chr5:174724582-174730896 | -1.86 | -3.64 | 1.26 | 13.41 | 2.51E-04 | 1.54E-02 | -1.81 |
| *RARRES2* | ENSG00000106538.10 | chr7:150338317-150341662 | -1.34 | -2.52 | 1.98 | 13.40 | 2.52E-04 | 1.54E-02 | -1.81 |
| *E2F5* | ENSG00000133740.11 | chr8:85177154-85217158 | 0.48 | 1.39 | 4.96 | 13.40 | 2.52E-04 | 1.54E-02 | -1.81 |
| *KDELR3* | ENSG00000100196.11 | chr22:38468078-38483447 | -0.74 | -1.67 | 3.62 | 13.39 | 2.54E-04 | 1.55E-02 | -1.81 |
| *TXNRD2* | ENSG00000184470.21 | chr22:19875517-19941820 | -2.02 | -4.04 | 1.07 | 13.38 | 2.55E-04 | 1.55E-02 | -1.81 |
| *NEUROD6* | ENSG00000164600.7 | chr7:31337465-31340726 | -0.56 | -1.48 | 4.41 | 13.36 | 2.57E-04 | 1.57E-02 | -1.81 |
| *OXR1* | ENSG00000164830.18 | chr8:106359476-106752694 | 0.36 | 1.29 | 5.91 | 13.34 | 2.60E-04 | 1.58E-02 | -1.80 |
| *ZMYND10* | ENSG00000004838.14 | chr3:50341110-50345732 | -0.75 | -1.68 | 3.60 | 13.33 | 2.62E-04 | 1.59E-02 | -1.80 |
| *SP140L* | ENSG00000185404.16 | chr2:230327184-230403732 | 0.79 | 1.73 | 3.39 | 13.30 | 2.66E-04 | 1.62E-02 | -1.79 |
| *SYNE2* | ENSG00000054654.19 | chr14:63761899-64226433 | -0.36 | -1.29 | 5.92 | 13.29 | 2.67E-04 | 1.62E-02 | -1.79 |
| *SLC12A5* | ENSG00000124140.14 | chr20:46021690-46060150 | 0.45 | 1.36 | 5.16 | 13.28 | 2.68E-04 | 1.62E-02 | -1.79 |
| *TBR1* | ENSG00000136535.15 | chr2:161416297-161425870 | -2.16 | -4.46 | 0.90 | 13.27 | 2.70E-04 | 1.64E-02 | -1.79 |
| *UBXN6* | ENSG00000167671.12 | chr19:4444999-4457794 | 0.36 | 1.28 | 6.00 | 13.25 | 2.73E-04 | 1.65E-02 | -1.78 |
| *VAX1* | ENSG00000148704.13 | chr10:117128521-117138301 | -0.84 | -1.79 | 3.21 | 13.24 | 2.73E-04 | 1.65E-02 | -1.78 |
| *ANXA1* | ENSG00000135046.14 | chr9:73151865-73170393 | -0.39 | -1.31 | 5.57 | 13.22 | 2.77E-04 | 1.67E-02 | -1.78 |
| *C1orf21* | ENSG00000116667.15 | chr1:184387029-184629019 | 0.36 | 1.29 | 5.92 | 13.18 | 2.84E-04 | 1.71E-02 | -1.77 |
| *COPS3* | ENSG00000141030.13 | chr17:17246616-17281273 | 0.55 | 1.47 | 4.42 | 13.13 | 2.91E-04 | 1.75E-02 | -1.76 |
| *CPT1C* | ENSG00000169169.15 | chr19:49690898-49713731 | -0.33 | -1.26 | 6.30 | 13.11 | 2.94E-04 | 1.76E-02 | -1.75 |
| *KY* | ENSG00000174611.12 | chr3:134599923-134651636 | 5.04 | 32.99 | -0.66 | 13.06 | 3.02E-04 | 1.81E-02 | -1.74 |
| *PDCD5* | ENSG00000105185.12 | chr19:32581190-32587453 | 0.53 | 1.45 | 4.57 | 13.01 | 3.10E-04 | 1.85E-02 | -1.73 |
| *SLC25A33* | ENSG00000171612.7 | chr1:9539465-9585173 | 0.69 | 1.62 | 3.77 | 13.01 | 3.11E-04 | 1.85E-02 | -1.73 |
| *MFSD4B* | ENSG00000173214.7 | chr6:111259327-111445354 | -0.63 | -1.54 | 4.07 | 13.00 | 3.11E-04 | 1.85E-02 | -1.73 |
| *TENT5C* | ENSG00000183508.5 | chr1:117606048-117628389 | -1.02 | -2.03 | 2.69 | 12.99 | 3.13E-04 | 1.86E-02 | -1.73 |
| *CPT1A* | ENSG00000110090.13 | chr11:68754620-68844410 | -0.82 | -1.76 | 3.27 | 12.97 | 3.16E-04 | 1.88E-02 | -1.73 |
| *STON1* | ENSG00000243244.7 | chr2:48529383-48598513 | -0.51 | -1.42 | 4.73 | 12.97 | 3.16E-04 | 1.88E-02 | -1.73 |
| *APOH* | ENSG00000091583.11 | chr17:66212033-66256525 | -2.05 | -4.13 | 0.99 | 12.97 | 3.17E-04 | 1.88E-02 | -1.73 |
| *TNK2* | ENSG00000061938.19 | chr3:195863364-195911945 | 0.38 | 1.31 | 5.60 | 12.96 | 3.18E-04 | 1.88E-02 | -1.73 |
| *BRINP1* | ENSG00000078725.13 | chr9:119153458-119369435 | -0.34 | -1.27 | 6.16 | 12.95 | 3.20E-04 | 1.89E-02 | -1.72 |
| *SNPH* | ENSG00000101298.15 | chr20:1266280-1309328 | 0.35 | 1.28 | 5.97 | 12.95 | 3.20E-04 | 1.89E-02 | -1.72 |
| *AGBL4* | ENSG00000186094.17 | chr1:48532854-50023954 | 1.61 | 3.06 | 1.50 | 12.94 | 3.21E-04 | 1.90E-02 | -1.72 |
| *C16orf74* | ENSG00000154102.11 | chr16:85690084-85751129 | 1.46 | 2.75 | 1.73 | 12.93 | 3.24E-04 | 1.91E-02 | -1.72 |
| *ZIC3* | ENSG00000156925.12 | chrX:137566127-137577691 | -1.21 | -2.31 | 2.21 | 12.91 | 3.27E-04 | 1.93E-02 | -1.71 |
| *RTN4RL2* | ENSG00000186907.8 | chr11:57460528-57477534 | 0.51 | 1.42 | 4.72 | 12.90 | 3.29E-04 | 1.94E-02 | -1.71 |
| *POLR2J* | ENSG00000005075.15 | chr7:102473118-102478907 | 0.41 | 1.33 | 5.39 | 12.89 | 3.30E-04 | 1.94E-02 | -1.71 |
| *CBLN1* | ENSG00000102924.12 | chr16:49277917-49281838 | 0.81 | 1.76 | 3.25 | 12.88 | 3.32E-04 | 1.95E-02 | -1.71 |
| *ZNF326* | ENSG00000162664.17 | chr1:89995110-90035533 | -0.39 | -1.31 | 5.50 | 12.86 | 3.36E-04 | 1.97E-02 | -1.71 |
| *RYR3* | ENSG00000198838.14 | chr15:33310962-33866121 | -0.45 | -1.37 | 5.09 | 12.82 | 3.43E-04 | 2.01E-02 | -1.70 |
| *FAM184A* | ENSG00000111879.20 | chr6:118959763-119149387 | -0.33 | -1.26 | 6.23 | 12.81 | 3.44E-04 | 2.01E-02 | -1.70 |
| *EEPD1* | ENSG00000122547.11 | chr7:36153254-36301538 | 0.63 | 1.55 | 4.00 | 12.80 | 3.47E-04 | 2.03E-02 | -1.69 |
| *NFATC2* | ENSG00000101096.20 | chr20:51386957-51562831 | -1.69 | -3.24 | 1.41 | 12.74 | 3.58E-04 | 2.08E-02 | -1.68 |
| *AJM1* | ENSG00000232434.2 | chr9:136844415-136848801 | 0.37 | 1.29 | 5.76 | 12.72 | 3.63E-04 | 2.11E-02 | -1.68 |
| *EPHA4* | ENSG00000116106.12 | chr2:221418027-221574202 | -0.68 | -1.60 | 3.80 | 12.70 | 3.65E-04 | 2.12E-02 | -1.67 |
| *ENTPD5* | ENSG00000187097.12 | chr14:73958010-74019399 | 0.42 | 1.34 | 5.24 | 12.65 | 3.75E-04 | 2.17E-02 | -1.66 |
| *REPS1* | ENSG00000135597.19 | chr6:138903493-138988261 | 0.34 | 1.27 | 6.08 | 12.64 | 3.78E-04 | 2.18E-02 | -1.66 |
| *CHI3L1* | ENSG00000133048.13 | chr1:203178931-203186704 | 0.88 | 1.84 | 3.00 | 12.62 | 3.82E-04 | 2.21E-02 | -1.66 |
| *KCNJ6* | ENSG00000157542.11 | chr21:37607373-38121345 | -0.46 | -1.38 | 4.98 | 12.61 | 3.84E-04 | 2.21E-02 | -1.66 |
| *NWD2* | ENSG00000174145.8 | chr4:37244743-37449463 | -0.90 | -1.86 | 2.98 | 12.60 | 3.85E-04 | 2.22E-02 | -1.65 |
| *C7orf61* | ENSG00000185955.5 | chr7:100456620-100464260 | 8.06 | 267.29 | -1.23 | 12.58 | 3.91E-04 | 2.24E-02 | -1.65 |
| *FHIT* | ENSG00000189283.10 | chr3:59747277-61251459 | -1.12 | -2.18 | 2.39 | 12.57 | 3.92E-04 | 2.25E-02 | -1.65 |
| *LCA5* | ENSG00000135338.14 | chr6:79484991-79537458 | -0.56 | -1.47 | 4.35 | 12.56 | 3.94E-04 | 2.26E-02 | -1.65 |
| *ESRRA* | ENSG00000173153.16 | chr11:64305497-64316743 | 0.89 | 1.86 | 2.97 | 12.54 | 3.99E-04 | 2.28E-02 | -1.64 |
| *RBPMS* | ENSG00000157110.16 | chr8:30384511-30572256 | -0.84 | -1.79 | 3.12 | 12.53 | 4.01E-04 | 2.29E-02 | -1.64 |
| *PHF24* | ENSG00000122733.12 | chr9:34957608-34982544 | 0.37 | 1.29 | 5.73 | 12.51 | 4.04E-04 | 2.30E-02 | -1.64 |
| *RGS7* | ENSG00000182901.17 | chr1:240775514-241357230 | 0.68 | 1.61 | 3.75 | 12.45 | 4.17E-04 | 2.37E-02 | -1.63 |
| *TGFB3* | ENSG00000119699.8 | chr14:75958097-75983011 | -0.76 | -1.70 | 3.42 | 12.41 | 4.28E-04 | 2.42E-02 | -1.62 |
| *SOCS7* | ENSG00000274211.6 | chr17:38351844-38405593 | 0.34 | 1.26 | 6.11 | 12.41 | 4.27E-04 | 2.42E-02 | -1.62 |
| *NECTIN3* | ENSG00000177707.11 | chr3:111070071-111275563 | -0.38 | -1.30 | 5.52 | 12.40 | 4.29E-04 | 2.43E-02 | -1.62 |
| *TSC22D3* | ENSG00000157514.16 | chrX:107713221-107777342 | 0.49 | 1.40 | 4.80 | 12.40 | 4.29E-04 | 2.43E-02 | -1.62 |
| *C2orf50* | ENSG00000150873.12 | chr2:11133128-11151317 | -0.67 | -1.59 | 3.80 | 12.35 | 4.42E-04 | 2.49E-02 | -1.60 |
| *TNFRSF10C* | ENSG00000173535.15 | chr8:23102921-23117445 | 1.37 | 2.59 | 1.81 | 12.31 | 4.51E-04 | 2.53E-02 | -1.60 |
| *ARHGEF2* | ENSG00000116584.20 | chr1:155946851-156007070 | 0.38 | 1.30 | 5.59 | 12.30 | 4.52E-04 | 2.54E-02 | -1.60 |
| *NFKB1* | ENSG00000109320.13 | chr4:102501331-102617302 | -0.56 | -1.47 | 4.31 | 12.25 | 4.65E-04 | 2.60E-02 | -1.58 |
| *COA1* | ENSG00000106603.19 | chr7:43608456-43729717 | -0.57 | -1.48 | 4.25 | 12.21 | 4.77E-04 | 2.66E-02 | -1.58 |
| *NME3* | ENSG00000103024.7 | chr16:1770286-1771730 | -0.80 | -1.74 | 3.23 | 12.20 | 4.77E-04 | 2.66E-02 | -1.57 |
| *CHST2* | ENSG00000175040.6 | chr3:143119771-143124014 | -0.39 | -1.31 | 5.42 | 12.20 | 4.79E-04 | 2.66E-02 | -1.57 |
| *SRSF1* | ENSG00000136450.13 | chr17:58000919-58007346 | -0.63 | -1.54 | 3.98 | 12.18 | 4.82E-04 | 2.68E-02 | -1.57 |
| *ADAMTS3* | ENSG00000156140.10 | chr4:72280969-72569221 | 0.43 | 1.35 | 5.13 | 12.18 | 4.83E-04 | 2.68E-02 | -1.57 |
| *UQCR10* | ENSG00000184076.13 | chr22:29767369-29770413 | 0.43 | 1.35 | 5.13 | 12.18 | 4.84E-04 | 2.68E-02 | -1.57 |
| *CEP41* | ENSG00000106477.20 | chr7:130393771-130442433 | -0.38 | -1.30 | 5.50 | 12.16 | 4.88E-04 | 2.70E-02 | -1.57 |
| *SLC25A16* | ENSG00000122912.15 | chr10:68477998-68527523 | 0.59 | 1.50 | 4.16 | 12.16 | 4.90E-04 | 2.71E-02 | -1.57 |
| *IQCA1* | ENSG00000132321.17 | chr2:236324147-236507535 | -2.53 | -5.78 | 0.50 | 12.13 | 4.96E-04 | 2.74E-02 | -1.56 |
| *LRP1B* | ENSG00000168702.18 | chr2:140231423-142131016 | -0.51 | -1.43 | 4.59 | 12.13 | 4.97E-04 | 2.74E-02 | -1.56 |
| *PAQR6* | ENSG00000160781.17 | chr1:156243320-156248117 | 0.39 | 1.31 | 5.43 | 12.13 | 4.97E-04 | 2.74E-02 | -1.56 |
| *CD109* | ENSG00000156535.15 | chr6:73695785-73828316 | -0.55 | -1.47 | 4.31 | 12.09 | 5.08E-04 | 2.79E-02 | -1.55 |
| *SERINC5* | ENSG00000164300.17 | chr5:80111651-80256048 | 0.34 | 1.26 | 6.04 | 12.09 | 5.07E-04 | 2.79E-02 | -1.55 |
| *PLD5* | ENSG00000180287.17 | chr1:242082986-242524697 | 0.68 | 1.61 | 3.71 | 12.08 | 5.11E-04 | 2.80E-02 | -1.55 |
| *SCRT1* | ENSG00000261678.3 | chr8:144330565-144336482 | 0.46 | 1.37 | 4.93 | 12.07 | 5.12E-04 | 2.81E-02 | -1.55 |
| *RAB39B* | ENSG00000155961.5 | chrX:155258235-155264491 | 0.42 | 1.34 | 5.18 | 12.00 | 5.31E-04 | 2.90E-02 | -1.54 |
| *ZNF37A* | ENSG00000075407.18 | chr10:38094334-38150293 | -0.62 | -1.53 | 4.01 | 11.98 | 5.39E-04 | 2.93E-02 | -1.53 |
| *ACAT2* | ENSG00000120437.9 | chr6:159762045-159779112 | -0.44 | -1.35 | 5.08 | 11.95 | 5.46E-04 | 2.97E-02 | -1.53 |
| *GPR137B* | ENSG00000077585.14 | chr1:236142505-236221865 | 0.34 | 1.26 | 6.01 | 11.95 | 5.47E-04 | 2.97E-02 | -1.53 |
| *C5orf38* | ENSG00000186493.13 | chr5:2752131-2755397 | 0.75 | 1.68 | 3.40 | 11.95 | 5.48E-04 | 2.97E-02 | -1.53 |
| *POTEF* | ENSG00000196604.13 | chr2:130073535-130129222 | -0.79 | -1.73 | 3.21 | 11.87 | 5.71E-04 | 3.08E-02 | -1.51 |
| *RSPO2* | ENSG00000147655.12 | chr8:107899316-108083642 | -2.94 | -7.68 | 0.20 | 11.86 | 5.72E-04 | 3.09E-02 | -1.51 |
| *CACNG3* | ENSG00000006116.4 | chr16:24256335-24362412 | -0.52 | -1.43 | 4.52 | 11.86 | 5.75E-04 | 3.09E-02 | -1.51 |
| *GPX8* | ENSG00000164294.14 | chr5:55160167-55167297 | -0.69 | -1.61 | 3.67 | 11.81 | 5.90E-04 | 3.17E-02 | -1.50 |
| *BCAT1* | ENSG00000060982.15 | chr12:24810024-24949101 | -0.34 | -1.26 | 5.98 | 11.81 | 5.90E-04 | 3.17E-02 | -1.50 |
| *ZNF638* | ENSG00000075292.19 | chr2:71276561-71435069 | -0.32 | -1.25 | 6.19 | 11.79 | 5.95E-04 | 3.19E-02 | -1.50 |
| *LSR* | ENSG00000105699.16 | chr19:35248330-35267964 | -1.62 | -3.07 | 1.42 | 11.78 | 5.98E-04 | 3.20E-02 | -1.49 |
| *LRRC8D* | ENSG00000171492.14 | chr1:89821014-89936611 | -0.34 | -1.27 | 5.96 | 11.78 | 6.00E-04 | 3.21E-02 | -1.49 |
| *DOK1* | ENSG00000115325.13 | chr2:74549026-74557554 | 0.83 | 1.78 | 3.06 | 11.76 | 6.06E-04 | 3.24E-02 | -1.49 |
| *PRR14* | ENSG00000156858.12 | chr16:30650717-30656413 | 0.34 | 1.27 | 5.91 | 11.73 | 6.15E-04 | 3.28E-02 | -1.48 |
| *C4orf50* | ENSG00000181215.15 | chr4:5897373-6200555 | -0.91 | -1.88 | 2.84 | 11.70 | 6.25E-04 | 3.31E-02 | -1.48 |
| *C4orf19* | ENSG00000154274.15 | chr4:37453925-37623495 | -0.50 | -1.41 | 4.63 | 11.70 | 6.26E-04 | 3.31E-02 | -1.48 |
| *FAM204A* | ENSG00000165669.14 | chr10:118297925-118342328 | 0.54 | 1.46 | 4.30 | 11.70 | 6.25E-04 | 3.31E-02 | -1.48 |
| *ADAMTS15* | ENSG00000166106.3 | chr11:130448974-130476641 | 0.53 | 1.45 | 4.36 | 11.67 | 6.37E-04 | 3.37E-02 | -1.47 |
| *STIM2* | ENSG00000109689.17 | chr4:26857601-27025381 | 0.32 | 1.25 | 6.15 | 11.66 | 6.37E-04 | 3.37E-02 | -1.47 |
| *ZNF865* | ENSG00000261221.3 | chr19:55605405-55617269 | 0.33 | 1.26 | 6.03 | 11.65 | 6.42E-04 | 3.39E-02 | -1.47 |
| *YPEL3* | ENSG00000090238.12 | chr16:30092314-30096915 | 0.42 | 1.33 | 5.17 | 11.62 | 6.51E-04 | 3.43E-02 | -1.46 |
| *ZC3H12C* | ENSG00000149289.11 | chr11:110093392-110171841 | -0.35 | -1.27 | 5.80 | 11.61 | 6.55E-04 | 3.44E-02 | -1.46 |
| *FOXB1* | ENSG00000171956.7 | chr15:60004234-60061730 | 0.67 | 1.59 | 3.72 | 11.60 | 6.59E-04 | 3.46E-02 | -1.46 |
| *KCNC4* | ENSG00000116396.15 | chr1:110210314-110283100 | 0.40 | 1.32 | 5.30 | 11.60 | 6.59E-04 | 3.46E-02 | -1.46 |
| *ATM* | ENSG00000149311.19 | chr11:108222832-108369102 | -0.37 | -1.29 | 5.59 | 11.59 | 6.62E-04 | 3.47E-02 | -1.46 |
| *MSRB3* | ENSG00000174099.12 | chr12:65278643-65491430 | -0.51 | -1.42 | 4.54 | 11.59 | 6.64E-04 | 3.48E-02 | -1.46 |
| *MRPL41* | ENSG00000182154.8 | chr9:137551879-137552555 | 0.47 | 1.39 | 4.78 | 11.58 | 6.66E-04 | 3.48E-02 | -1.46 |
| *NHSL2* | ENSG00000204131.9 | chrX:71910818-72161750 | -0.45 | -1.37 | 4.92 | 11.54 | 6.82E-04 | 3.56E-02 | -1.45 |
| *ITPRIP* | ENSG00000148841.17 | chr10:104309698-104338465 | -0.57 | -1.48 | 4.18 | 11.51 | 6.92E-04 | 3.60E-02 | -1.44 |
| *UHMK1* | ENSG00000152332.16 | chr1:162497251-162529631 | -0.35 | -1.28 | 5.73 | 11.51 | 6.92E-04 | 3.60E-02 | -1.44 |
| *DMTN* | ENSG00000158856.18 | chr8:22048995-22082527 | 0.38 | 1.30 | 5.44 | 11.46 | 7.10E-04 | 3.68E-02 | -1.43 |
| *NBEAL2* | ENSG00000160796.18 | chr3:46979666-47009704 | -0.51 | -1.43 | 4.47 | 11.46 | 7.13E-04 | 3.68E-02 | -1.43 |
| *COL6A3* | ENSG00000163359.16 | chr2:237324003-237414207 | -0.35 | -1.28 | 5.69 | 11.45 | 7.14E-04 | 3.69E-02 | -1.43 |
| *NBAS* | ENSG00000151779.13 | chr2:15166914-15561334 | -0.33 | -1.26 | 6.05 | 11.43 | 7.23E-04 | 3.72E-02 | -1.43 |
| *RELN* | ENSG00000189056.14 | chr7:103471784-103989658 | 0.35 | 1.28 | 5.70 | 11.41 | 7.33E-04 | 3.76E-02 | -1.42 |
| *AMIGO2* | ENSG00000139211.6 | chr12:47075707-47079951 | -0.96 | -1.94 | 2.67 | 11.38 | 7.43E-04 | 3.80E-02 | -1.42 |
| *PPP2R5B* | ENSG00000068971.14 | chr11:64917553-64934475 | 0.33 | 1.26 | 6.00 | 11.37 | 7.46E-04 | 3.82E-02 | -1.42 |
| *TSPYL5* | ENSG00000180543.5 | chr8:97273488-97277928 | -8.03 | -261.07 | -1.19 | 11.36 | 7.50E-04 | 3.83E-02 | -1.42 |
| *DLX6* | ENSG00000006377.11 | chr7:97005553-97011040 | 1.24 | 2.37 | 1.93 | 11.36 | 7.51E-04 | 3.83E-02 | -1.42 |
| *DNAJC18* | ENSG00000170464.10 | chr5:139408588-139444491 | 0.33 | 1.25 | 6.04 | 11.34 | 7.57E-04 | 3.85E-02 | -1.41 |
| *OSGIN1* | ENSG00000140961.14 | chr16:83931311-83966332 | 1.36 | 2.57 | 1.74 | 11.34 | 7.59E-04 | 3.86E-02 | -1.41 |
| *MINAR1* | ENSG00000169330.9 | chr15:79432336-79472304 | 0.38 | 1.30 | 5.43 | 11.30 | 7.75E-04 | 3.93E-02 | -1.41 |
| *SPAG6* | ENSG00000077327.16 | chr10:22345445-22454224 | -1.61 | -3.06 | 1.37 | 11.29 | 7.80E-04 | 3.95E-02 | -1.40 |
| *SGTB* | ENSG00000197860.10 | chr5:65665928-65723035 | 0.33 | 1.26 | 5.98 | 11.25 | 7.97E-04 | 4.03E-02 | -1.40 |
| *IL17RA* | ENSG00000177663.14 | chr22:17084954-17115694 | -0.42 | -1.34 | 5.08 | 11.23 | 8.06E-04 | 4.07E-02 | -1.39 |
| *SYNPR* | ENSG00000163630.11 | chr3:63228315-63616924 | 0.95 | 1.94 | 2.66 | 11.23 | 8.07E-04 | 4.07E-02 | -1.39 |
| *PDRG1* | ENSG00000088356.6 | chr20:31944337-31952046 | 0.49 | 1.40 | 4.62 | 11.22 | 8.08E-04 | 4.07E-02 | -1.39 |
| *TMEM132B* | ENSG00000139364.10 | chr12:125186836-125662377 | -0.68 | -1.60 | 3.62 | 11.20 | 8.17E-04 | 4.10E-02 | -1.39 |
| *FSTL3* | ENSG00000070404.10 | chr19:676392-683392 | -0.47 | -1.39 | 4.72 | 11.21 | 8.15E-04 | 4.10E-02 | -1.39 |
| *ABHD6* | ENSG00000163686.15 | chr3:58237532-58295693 | -0.45 | -1.37 | 4.87 | 11.21 | 8.14E-04 | 4.10E-02 | -1.39 |
| *MYLIP* | ENSG00000007944.15 | chr6:16129086-16148248 | -0.54 | -1.46 | 4.26 | 11.18 | 8.26E-04 | 4.13E-02 | -1.38 |
| *NRN1* | ENSG00000124785.9 | chr6:5997999-6007605 | -0.45 | -1.37 | 4.85 | 11.16 | 8.38E-04 | 4.17E-02 | -1.38 |
| *RANBP3L* | ENSG00000164188.9 | chr5:36246913-36302114 | 0.73 | 1.66 | 3.37 | 11.14 | 8.46E-04 | 4.20E-02 | -1.38 |
| *COX7A2L* | ENSG00000115944.15 | chr2:42333546-42425088 | 0.34 | 1.26 | 5.83 | 11.11 | 8.59E-04 | 4.25E-02 | -1.37 |
| *KCNQ3* | ENSG00000184156.17 | chr8:132120859-132481095 | -0.33 | -1.26 | 5.94 | 11.09 | 8.68E-04 | 4.28E-02 | -1.37 |
| *RNF150* | ENSG00000170153.11 | chr4:140859807-141212877 | -0.37 | -1.30 | 5.42 | 11.07 | 8.79E-04 | 4.33E-02 | -1.36 |
| *LSM7* | ENSG00000130332.15 | chr19:2321520-2328611 | 0.65 | 1.57 | 3.74 | 11.06 | 8.82E-04 | 4.34E-02 | -1.36 |
| *JPH3* | ENSG00000154118.13 | chr16:87601835-87698156 | 0.34 | 1.27 | 5.78 | 11.05 | 8.85E-04 | 4.35E-02 | -1.36 |
| *USP30* | ENSG00000135093.13 | chr12:109023089-109088023 | 0.44 | 1.36 | 4.93 | 11.05 | 8.88E-04 | 4.36E-02 | -1.36 |
| *ADGRA3* | ENSG00000152990.14 | chr4:22345071-22516066 | 0.38 | 1.30 | 5.37 | 11.03 | 8.99E-04 | 4.40E-02 | -1.36 |
| *PITPNM3* | ENSG00000091622.16 | chr17:6451263-6556555 | 0.32 | 1.25 | 6.02 | 11.00 | 9.11E-04 | 4.45E-02 | -1.35 |
| *CCR10* | ENSG00000184451.5 | chr17:42678889-42683917 | 1.01 | 2.01 | 2.48 | 11.00 | 9.12E-04 | 4.45E-02 | -1.35 |
| *ITPRIPL2* | ENSG00000205730.6 | chr16:19113932-19121629 | -0.40 | -1.32 | 5.20 | 11.00 | 9.13E-04 | 4.45E-02 | -1.35 |
| *PAQR3* | ENSG00000163291.14 | chr4:78887127-78939438 | 0.37 | 1.29 | 5.45 | 10.99 | 9.14E-04 | 4.46E-02 | -1.35 |
| *KCNT2* | ENSG00000162687.18 | chr1:196225779-196609225 | -0.64 | -1.56 | 3.76 | 10.96 | 9.30E-04 | 4.52E-02 | -1.34 |
| *DDX58* | ENSG00000107201.10 | chr9:32455302-32526208 | -0.80 | -1.74 | 3.08 | 10.95 | 9.38E-04 | 4.54E-02 | -1.34 |
| *CROT* | ENSG00000005469.12 | chr7:87345664-87399794 | -0.75 | -1.68 | 3.27 | 10.93 | 9.49E-04 | 4.59E-02 | -1.34 |
| *SHISAL2A* | ENSG00000182183.15 | chr1:52633168-52669683 | 3.81 | 14.05 | -0.44 | 10.90 | 9.63E-04 | 4.65E-02 | -1.33 |
| *PCSK6* | ENSG00000140479.17 | chr15:101297142-101525202 | -0.65 | -1.57 | 3.69 | 10.89 | 9.66E-04 | 4.66E-02 | -1.33 |
| *TRIM4* | ENSG00000146833.15 | chr7:99876958-99919600 | -7.96 | -249.77 | -1.23 | 10.88 | 9.74E-04 | 4.70E-02 | -1.33 |
| *TGFBR2* | ENSG00000163513.19 | chr3:30606601-30694142 | -0.86 | -1.81 | 2.89 | 10.85 | 9.86E-04 | 4.74E-02 | -1.32 |
| *GDPD2* | ENSG00000130055.14 | chrX:70423031-70433390 | -0.38 | -1.31 | 5.31 | 10.85 | 9.86E-04 | 4.74E-02 | -1.32 |
| *TIMM13* | ENSG00000099800.8 | chr19:2425625-2427586 | 0.38 | 1.30 | 5.32 | 10.85 | 9.86E-04 | 4.74E-02 | -1.32 |
| *IFFO2* | ENSG00000169991.11 | chr1:18904280-18956676 | -0.34 | -1.27 | 5.77 | 10.82 | 1.00E-03 | 4.80E-02 | -1.32 |
| *ENOPH1* | ENSG00000145293.16 | chr4:82430590-82461177 | 0.33 | 1.26 | 5.90 | 10.81 | 1.01E-03 | 4.83E-02 | -1.32 |
| *BDNF* | ENSG00000176697.20 | chr11:27654893-27722058 | -0.40 | -1.32 | 5.21 | 10.78 | 1.03E-03 | 4.89E-02 | -1.31 |
| *CDC14A* | ENSG00000079335.20 | chr1:100345001-100520277 | -0.75 | -1.68 | 3.23 | 10.76 | 1.04E-03 | 4.93E-02 | -1.31 |
| *NR3C1* | ENSG00000113580.15 | chr5:143277931-143435512 | -0.49 | -1.40 | 4.53 | 10.76 | 1.04E-03 | 4.94E-02 | -1.31 |
